# Drowning in a sandy ocean: Epiarenic growth of *Tillandsia* in the hyperarid Atacama Desert

**DOI:** 10.64898/2026.02.23.707457

**Authors:** Robin Schweikert, Ron Eric Stein, Neil Bogs, Olaf Bubenzer, Camilo del Río, Dörte Harpke, Simon Matthias May, Alexander Siegmund, Alexandra Stoll, Dietmar Quandt, Marcus A. Koch

**Author notes:** Correspondence to:* Marcus A. Koch. contributed equal to the work (Robin Schweikert, Ron Eric Stein).

## Abstract

The Atacama Desert hosts a unique ecosystem formed by the sand-dwelling *Tillandsia landbeckii*, which extends over hundreds of square kilometers. This vegetation relies primarily on fog as its main water source; however, aeolian sand also plays a crucial role in the long-term persistence of both the species and the overall plant community. The terrain is sloped and exposed to the prevailing wind direction. *Tillandsia* forms regular banding patterns oriented orthogonally to these landscape features. In this study, we aim to elucidate the abiotic–biotic interactions between sand properties and vegetation characteristics through a comparative approach. Three populations - Caldera, Oyarbide and Arica -, each spanning several square kilometers in the southern, central, and northern regions of the Chilean Atacama Desert, were selected to compare wind regimes, terrain structure, sand and substrate properties, and vegetation structure in order to identify common principles that maintain vegetation integrity. Data were collected from six climate stations, 1,246 substrate samples, population genomic data from 718 individuals, as well as satellite imagery and digital terrain models. Our findings demonstrate that regional wind systems transport sand from distant source areas, while near the ground, *Tillandsia* vegetation reduces wind velocity and traps sand, leading to the formation of moderately sorted sandy substrates that are similar across all three populations. Sites lacking or containing dead *Tillandsia* individuals often differ significantly in substrate characteristics. Genetic analyses indicate that *Tillandsia* populations exhibit strong spatial structure albeit recruiting high genetic diversity and an excess of heterozygosity, reflecting adaptation to the dynamic environmental conditions. We conclude that sand represents an essential component of this ecosystem, while *Tillandsia*, as the dominant biotic factor, actively shapes and maintains this distinctive desert environment.

**Graphical abstract:** 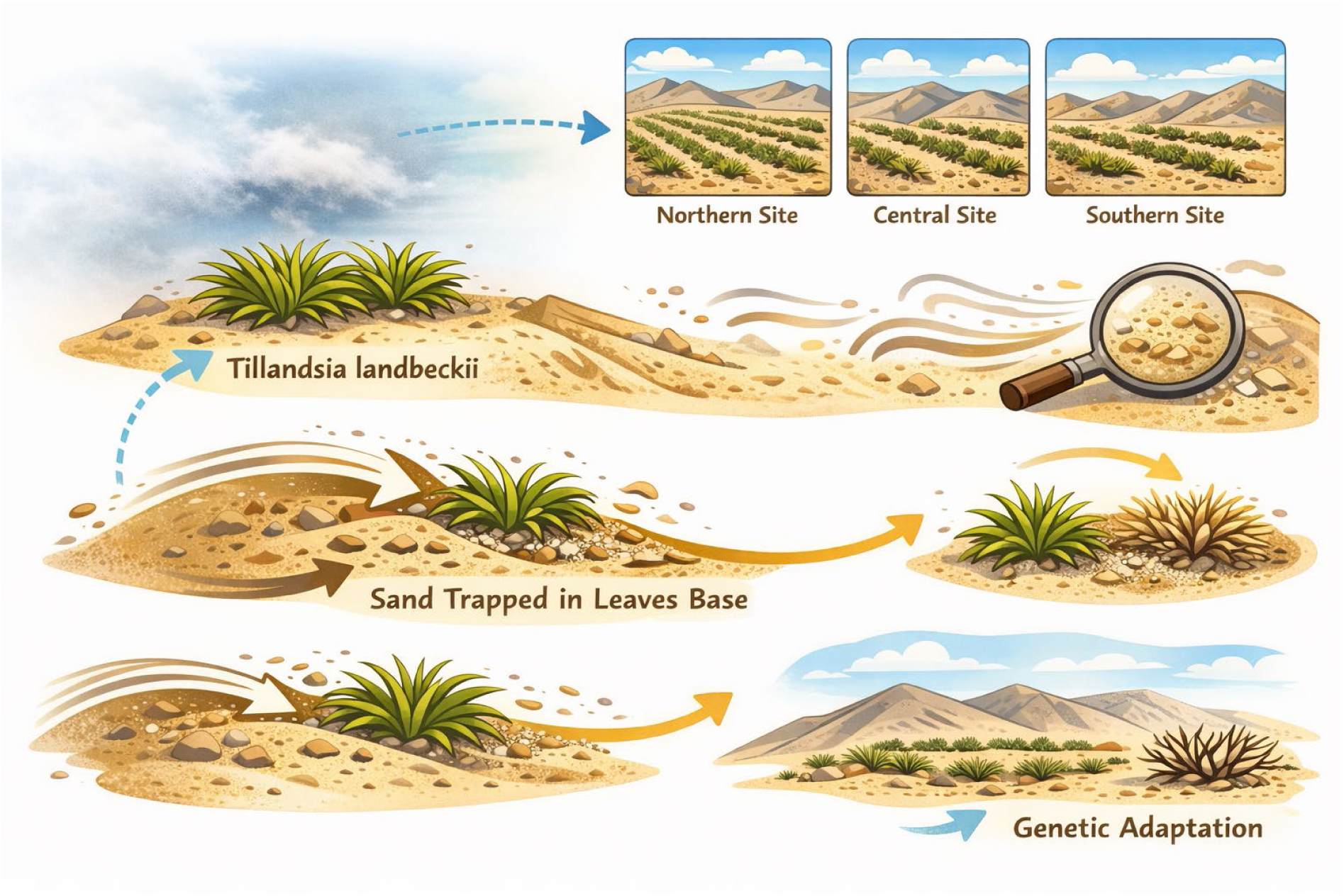

Generated based on own drawings and iterative improvements using ChatGPT while providing own peer-reviewed research contributions as input and baseline information (MAK).

**Short summary:** We exemplify unimodal regional wind systems facilitating sand transport toward *Tillandsiales. Tillandsiales* show a low-energy wind system allowing sand accumulation of predominant grain sizes available at each site. Thereby *Tillandsia landbeckii* modifies and maintains its own microenvironment. Genomic data reveal high clonality and excess of heterozygosity promoting fitness in a hyperarid environment, and abiotic factors drive the selection of diverse and adaptive *Tillandsia* phenotypes.

## 1 Introduction

Plant populations are subjected to numerous biotic and abiotic parameters driving their evolutionary fate in a spatio-temporal context. Under changing environmental conditions, the respective evolutionary unit, the species, may be confronted with scenarios of local adaptation (mitigation), extinction, range expansion or range shifts with the latter two cases requiring migration (Morris et al., 2020; Urban et al., 2016). However, ultimately, evolutionary dynamics may not only result in those spatial shifts, but may result also in newly created biological entities and diversification into new species (Willi et al., 2022). Most often plants are occurring as assemblages and floristic mosaics forming plant communities and vegetation units, and biotic and abiotic parameters jointly shape the structure and functioning of these higher-level units (Li and Prentice, 2024; Qian et al., 2023;). Monospecific vegetation, however, refers to plant communities in which a single species overwhelmingly dominates the available space, biomass, or canopy cover within a given area (Kremer et al., 2025). Such communities are characterized by low taxonomic and functional diversity and may occur in both natural and anthropogenically modified ecosystems (Chiappero et al., 2024; Liu et al., 2018;). From a community–ecological perspective, monospecific stands may represent either relatively stable endpoints under specific environmental filters or transient stages within successional trajectories following disturbance, invasion, or intensive management (Cubino et al., 2022; Long et al., 2025; Mendieta-Leiva et al., 2022). Operationally, they are often defined by high relative abundance or cover of a single species, with other species present only at low frequencies or entirely absent and is documented in a wide range of habitats often representing one or the other extreme of environmental parameters including agricultural landscapes (e.g., Lenné and Wood, 2022; Marani et al., 2013; Nuñez and Paritsis, 2018;).

The development and persistence of monospecific vegetation are governed by the interaction of abiotic factors, biotic interactions, and disturbance regimes. Abiotic factors such as extreme edaphic / sedimentological conditions (including high salinity, unusual pH, or limiting nutrient profiles) restrict the regional species pool to a subset of stress–tolerant taxa, thereby facilitating dominance by a single, highly adapted species (Frenette-Dussault et al., 2012; Grime, 1977). Hydrological regimes, including permanent waterlogging, seasonal flooding, or chronic soil drought, can similarly constrain community composition and favor particular life forms that are able to exploit these conditions, leading to monospecific stands in wetlands and drylands alike (Anderegg et al., 2016; Ihaddaden et al., 2013). At broader spatial scales, climate and topographic context modulate temperature, radiation, and moisture availability, further shaping the pool of species capable of persisting and potentially dominating under given environmental conditions (Moura et al., 2016). Biotic mechanisms are critical in reinforcing and maintaining monospecific dominance once established. Strong competitive asymmetries, arising from traits such as rapid growth, high propagule production, dense canopy structure, or extensive root systems, enable a dominant species to preempt limiting resources and suppress recruitment of co–occurring species (Nathan et al., 2015). Positive feedback, e.g. mediated by soil biota and mutualistic interactions, including specialized mycorrhizal associations or shifts in microbial communities, can enhance nutrient acquisition and stress tolerance of the dominant species while creating unfavorable conditions for potential competitors (Nuñez and Paritsis, 2018). Recurrent disturbances such as fire, grazing, mowing, and flooding can impose strong selective filters by favoring species with particular life–history traits (e.g., resprouting ability, rapid post–disturbance growth, or persistent seed banks), thereby enabling a single disturbance–tolerant species to achieve and maintain dominance under specific disturbance frequencies and intensities (Jaime et al., 2025; Wang et al., 2025). Collectively, these abiotic, biotic, and disturbance–driven processes interact to promote the establishment, consolidation, and long–term persistence of monospecific vegetation.

In arid and semi–arid environments, these mechanisms are accentuated by chronic water limitation, high evaporative demand, and strong temporal variability in rainfall, which severely constrain germination, establishment, and survival for many species. Under such conditions, species possessing traits such as deep or extensive root systems, pronounced drought tolerance, physiological plasticity, and efficient use or preemption of scarce resources can come to dominate extensive areas as monospecific stands, often composed of shrubs or perennial grasses (Wang et al., 2021). In areas with higher wind speeds and loose sand, vegetation is often damaged or even prevented by wind erosion (abrasion) (Goudie, 2022). Disturbances typical of drylands, including overgrazing, soil degradation, and altered fire regimes, may further simplify communities and provide a competitive advantage to these stress–tolerant dominants, thereby reinforcing and spatially expanding monospecific vegetation in arid landscapes (Wang et al., 2025).

There are several examples in arid and semiarid environments showing that these interactions in monospecific vegetation cause noticeable spatial structures. This fascinating phenomenon is often seen as an avenue to study complex abiotic-biotic interaction in simplified (one species) models, which often cover and form landscapes on km²–scale. Along bioclimatic gradients from humid to arid it has been proposed that barren gaps, such as fairy circles, in otherwise continuous vegetation are followed by linear vegetation pattern, and finally vegetation develops into highly isolated patches in extremely arid landscapes (Meron, 2019). A major focus has been given to banding patterns, which are formed by various plant species and sets of species in arid regions across continents. Banded vegetation in drylands (often called tiger bush, mogotes, or vegetation bands) consists of stripes of dense plants alternating with bare ground on gentle slopes in arid and semi–arid regions worldwide. The phenomenon is widespread, but published species lists are surprisingly incomplete and region–specific (Jürgens et al., 2022). In water–limited landscapes, small differences in infiltration and runoff create feedbacks where upslope bare zones shed water and sediment into downslope vegetated bands, which then maintain higher soil moisture, organic matter, and infiltration capacity. These eco–hydrological feedbacks lead to self–organized, quasi–periodic bands oriented roughly along contour lines, and similar banded patterns occur in Africa, Australia, the Americas, and parts of Asia on gentle slopes with sparse initial cover. Banded systems are ecologically important because they concentrate productivity, act as sinks for resources and hotspots of biodiversity, and their degradation (e. g. band thinning or collapse) is closely linked to desertification risk and global environmental change.

Past research on banded vegetation has focused far more on pattern formation theory and topographic mechanisms (Gandhi et al., 2018, Kästner et al., 2025), hydrological parameters (Paschalis et al. 2016; Wang & Zang 2019) or catastrophic responses (Rietkerk et al., 2004) than on feedback loops of biotic and abiotic interactions (Hidalgo-Ogalde et al., 2024). As a result, only a subset of species forming banded vegetation is documented explicitly, and there is no global synthesis that catalogues plant species known to occur or dominate in banded vegetation stripes. In the Chilean-Peruvian arid and hyperarid core of the Atacama Desert banded vegetation is prominently built up by only a few species of the genus *Tillandsia* from the bromeliad family (Koch et al., 2019; Koch et al., 2022; Möbus et al., 2021). The most prominent species is the fog-dependent *Tillandsia landbeckii*, that forms extensive monospecific “Tillandsia lomas” on coastal dune systems of the hyperarid Atacama Desert (Rundel et al., 1997), where mean annual rainfall is typically below 2 mm and the aridity index can fall at or below 0.0125, placing it at the dry limit for vascular plant life. Ecologically, these stands grow in striking linear bands oriented orthogonally to the slope, a pattern driven by unidirectional growth toward incoming fog, sand trapping around shoots, and progressive dieback and burial of older ramets, so that the vegetation itself helps shape coppice dunes (Nebkha) and stabilizes otherwise mobile substrates (Rundel et al., 1997). Biologically, *T. landbeckii* is an extreme atmospheric plant that lacks functional roots for water uptake, relying instead on leaf trichomes and CAM photosynthesis to harvest and efficiently use fog moisture under large daily temperature and radiation fluctuations (Raux et al., 2020); populations are long-lived, clonal yet genetically structured over tens to hundreds of kilometers, indicating both strong local adaptation (Jabbusch and Koch, 2025; Koch et al., 2020) and historical connectivity in this fragmented desert landscape (Koch et al., 2022; Latorre et al., 2011; Stein et al., 2023). In the hyperarid core, each plant also acts as a discrete microbial refugium: the phyllosphere and laimosphere harbor specialized, compartment-specific bacterial communities that persist where surrounding soils are almost devoid of life (Alfaro et al., 2021; Hakobyan et al., 2023; Jaeschke et al., 2024), highlighting the species’ dual role as a primary producer and as a keystone habitat for microorganisms at the very edge of terrestrial habitability.

The main factors determining *Tillandsia landbeckii* individual survival and vegetation integrity are regular occurrence of fog as the main source of water, aeolian sand as mobile but also stabilizing substrate, a sloped terrain allowing dune formation, wind systems transporting the fog and sand and serving as motor for environmental dynamics and a predominant and acrotonous vegetative growth form of a long-lived perennial plant. There were contributions on various parameters presented in the past (e.g. Del Rio et al., 2021; Garcia et al., 2021), but here we provide for the first time a study focusing on sand properties as well as the local wind system and combine and compare this information with vegetation and individual plant species characteristics on a fine-scale.

Herein we explore the hypotheses that *Tillandsia* builds its own “microhabitat” promoting growth and fitness while capturing sand in a continuous process. We hypothesize (H1) that a viable biotic system prioritizes and captures a defined sand grain size, and sand deposition and accumulation result in homogeneous substrate allowing acrotonal and branching growth with die-back and adequate oversanding at the base, thereby stabilizing plant architecture and vegetation integrity over decades. Accordingly, the range of captured sand grain sizes may reflect wind characteristics defining sand transport from regional source regions and fog from the Pacific coast. We also hypothesize (H2) that the biotic system responds collectively, enhancing the above-described processes, which is in return supporting clonal growth and vegetative propagation and should be manifested in a defined spatio-temporal population genetic structure.

For our study we selected three representative populations in the North, Center and South of the *Tillansia landbeckii* distribution range in the Chilean Atacama Desert, and we combine population genomic data with spatially matching substrate analyses, high resolution digital terrain models and weather station data of wind and temperature characteristics (Fig. 1).

**Figure 1.**
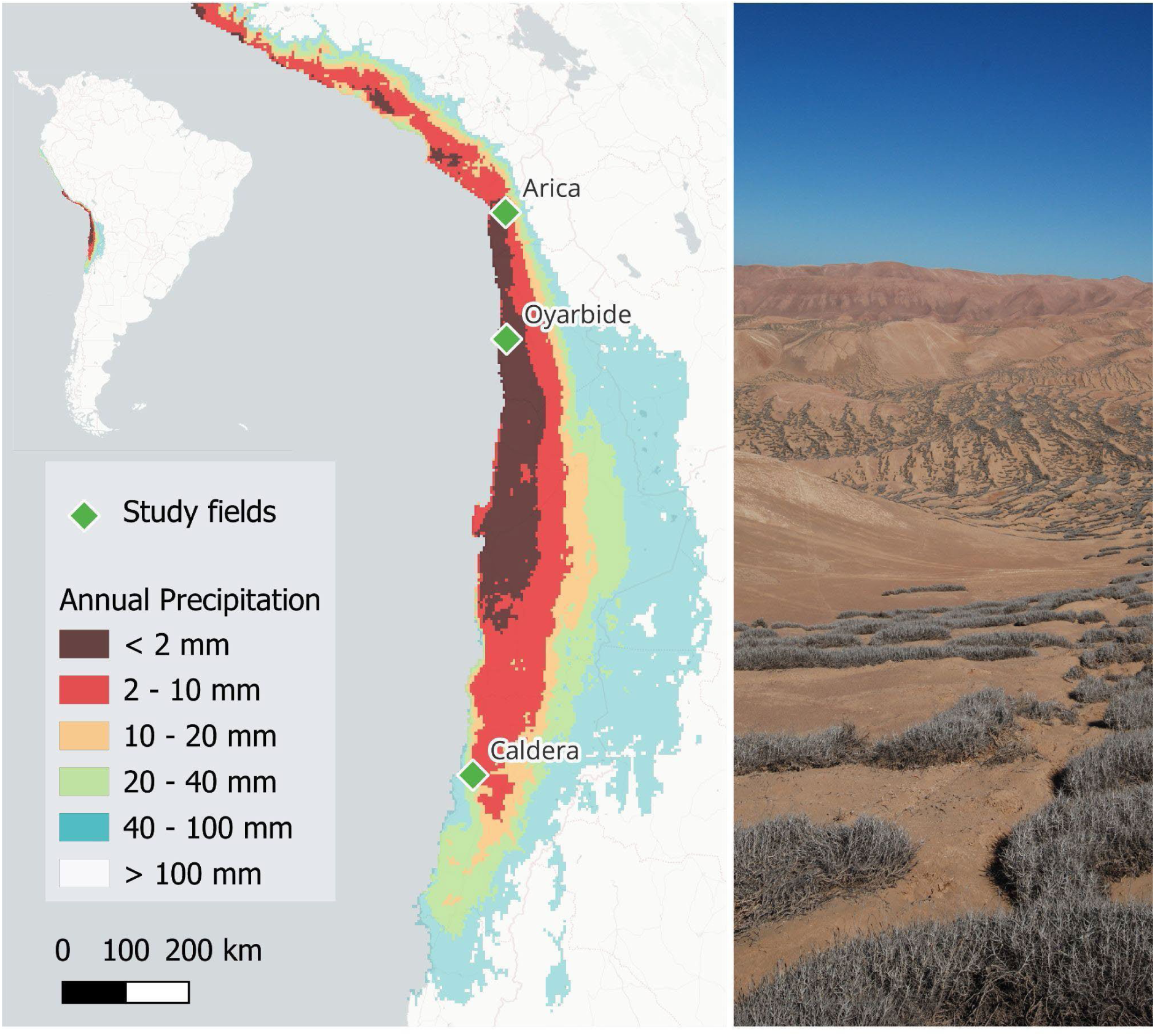
Map of *Tillandsia landbeckii* study sites in Chile. Overall distribution of *T. landbeckii* largely follows coastal regions within the very arid to hyperarid regions of the Atacama Desert (< 20 mm annual precipitation). Three study fields have been chosen to represent northern (Arica), central (Oyarbide) and southern (Caldera) populations. Color-codes show annual precipitation, based on WorldClim data (Bio12, https://www.worldclim.org/data/bioclim.html). Arica and Oyarbide are located within the “hyperarid core” (< 2 mm), Caldera is positioned in the less extreme transition zone with a more arid climate. The photograph shows part of the Oyarbide study field (taken by MAK).

## 2 Material and methods

### 2.1 Study area and comparative study site design

Our study area follows the biogeographic distribution of *Tillandsia landbeckii* in the hyperarid core of the Atacama Desert c. 10–25 km towards the inland along the Chilean Pacific coast. In particular in the hyperarid core of the northern Atacama Desert ecosystems experience less than 2 mm of precipitation (Mikulane et al., 2022), and average annual day and night temperature are c. 18.5 °C and 14 °C at about 700–1350 m a.s.l., respectively, with seasonal fluctuations (e.g., Schween et al., 2020; Schween, 2025). While the surface soils of the Atacama Desert are often dominated by fine-grained dusts of gypsum and other salts (Arens et al., 2021; Voigt et al., 2020), the occurrence of *Tillandsia landbeckii* is linked to the presence of sand. Based on our previous bio- and phylogeographic studies (Koch et al., 2022; Merklinger et al., 2020), distribution and ecological niche modelling (Stein et al., 2023) as well as population genetic analyses (Koch et al., 2019; Koch et al., 2020) three populations have been selected from the southern edge of the distribution range (hereafter named Caldera), the hyperarid core near Iquique (hereafter named Oyarbide) and the northernmost Chilean populations as represented by herein named population Arica (Fig. 1). This area stretches from 27.07° S to 18.48° S over a geographical distance of almost 1000 km. We did not consider the extinct southernmost and only historically described outpost of *Tillandsia landbeckii* from the Coquimbo region (31.65° S) (Smith and Downs, 1977; Till, 1992) as well as the recently discovered and explored populations in Peru (Stein et al., 2023). All three selected study sites are characterized with well-developed unispecific *Tillandsia landbeckii* lomas. However, in contrast to the Oyarbide population with *T. landbeckii* as the only flowering vascular plant species, there are few additional species found at the Arica population (e.g. including *T. marconae*) and even some higher plant diversity at Caldera (cacti, shrubs and annuals) indicating severe bioclimatic differences among populations. Genetically, all three populations are part of a coherent genepool, but each population differs significantly (Koch et al., 2022; Merklinger et al., 2020) for its genetic assignment.

All three study sites are located at a distance of approximately 10 km (Oyarbide) to 25 km (Arica and Caldera) from the Pacific coast and fully depend on fog or possibly dew as the only source of water for plant growth, and all sites are characterized and dominated by wind-transported (aeolian) sand, mainly composed of quartz and silicate minerals of igneous origin (amphibole, pyroxene and epidote), allowing to build up respective dune systems. The Atacama Desert is situated on the eastern flank of the subtropical Southeast Pacific anticyclone and the cold Humboldt Current and coastal upwelling cool the low troposphere, strengthening the inversion and supporting an extensive deck of marine stratocumulus clouds (Takahashi and Battisti, 2007). West to west-southwesterly wind directions prevail as a combination of geostrophic wind including the embedded low level coastal jet and a superimposed daily sea-breaze (Schween et al., 2020), which also drives fog penetration inland. Here, thermal induced weaker easterly winds dominate during the nights. The annual frequency of fog events or low clouds is 23 %, 13 %, and 20 % out of 365 days for Arica, Oyarbide, and Caldera, respectively, and fog distribution is mainly controlled by the horizontal and vertical dynamics of the marine boundary layer (Del Río et al., 2025).

Both, from a geological and a geomorphological point of view, there are different settings. According to the geological map 1:1 mio. (Sernageomin, 2003), the Arica site in the north is dominated be Lower-Middle Miocene sedimentary sequences of alluvial fans, pediments, or fluvial deposits composed of gravel, sand, and silt with interbedded ignimbrites as possible sediment sources for the Tillandsia sites. In addition, Pliocene-Pleistocene mass removal deposits of polymictic breccias with sand/silt matrix are less widespread. In Oyarbide, Pleistocene-Holocene sands and alluvial deposits above Jurassic and Tertiary rocks are providing, most probably, the source material, whereas in Caldera, again Pleistocene-Holocene alluvial and subordinated colluvial and lacustrine gravels, sands and silts are distributed. Whereas the orographies of Arica and Caldera are dominated and structured by valleys (Quebrada del Diabolo east of Arica and a slightly incised valley east of Caldera), the Oyarbide site is located above a steep coastal cliff on a steadily rising slope.

### 2.2 Experimental design and data sets

All three study sites were overlaid with a georeferenced grid system of 100 × 100 m. These grids were aligned to the linear arrangement of the vegetation. Coordinates were extracted for all grid centres and used further as way-points to navigate and collect samples. Total area sizes – stretching over *Tillandsia* loma vegetation – were 3.68 km² (368 grid cells) at Oyarbide, 4.91 km² (491 grid cells) at Caldera and 4.47 km² (447 grid cells) at Arica. For the three study sites elevation increases from 430–798 m a.s.l., 822–1168 m a.s.l., and 1116–1253 m a.s.l. at Caldera, Arica and Oyarbide, respectively.

At all three study population sites climate stations have been installed at the bottom and at the top area to collect weather data. For this study we gathered respective data on wind speed and wind direction for >12 months to determine parameters relevant for sand dynamics.

To calculate further geomorphological parameters relevant for sand transportation (elevation, exposition and slope) we utilized our earlier presented high resolution Digital Terrain Models (DTMs) of Arica (Stein et al., 2024a), Caldera (Stein et al., 2024b) and Oyarbide (Mikulane et al., 2022) complemented by freely available satellite data (COPDEM GLO-30, https://doi.org/10.5270/ESA-c5d3d65).

### 2.3 Sand and substrate analysis

#### 2.3.1 Sand and substrate collection

Within each of the study populations, substrate samples were taken from the centre of a 100 m x 100 m grid cell directly uphill and in front of a *Tillandsia landbeckii* plant. In case of missing plants near the grid cell centre we selected the nearest position for sampling, or if there have been no *Tillandsia*s in the entire grid cell substrate samples have been taken from the centre of the grid cell, too. 50 mL substrate samples were collected each from the upper 2–10 cm substrate layer in Falcon tubes for subsequent processing in the lab. Accordingly, we intended to avoid any recent and occasional surface disturbances. In total we collected 1,246 substrate samples (413, 492 and 388 samples, for Arica, Caldera, and Oyarbide, respectively). Sample codes, sample information, elevation and exact coordinates are provided with Supplementary material S1.

#### 2.3.2 Grain size determination by laser diffractometry

Prior to grain-size analysis, samples were subjected to mechanical and chemical preparation. Approximately 15 g (±1 g) of raw material was dry-sieved with a mesh-size of 2 mm, and particles larger than 2 mm were discarded. Based on estimated sample composition, weighted aliquots of the < 2 mm fraction were transferred into 12 mL PE reaction tubes: 0.6–0.8 g for coarse sand detection (0.63–2 mm), 0.4–0.6 g for medium sand detection (0.063–0.2 mm), and 0.15–0.3 g for silt- and clay-rich samples (< 0.063 mm).

Reaction tubes were filled one-third with dH₂O, treated with 1 mL of 12 % HCl and incubated overnight. After removing the supernatant, 1 mL of 15 % H₂O₂ was added and samples were incubated for three days at room temperature to dissolve any organic material. Samples were then washed three times with full tube volume of dH₂O followed by centrifugation (2500 rpm for 5 min). The final pellet was resuspended in two-thirds tube volume of dH₂O with 1 mL sodium pyrophosphate (Na₂HPO₃) and placed on an overhead shaker for thorough mixing until use for analysis for at least 12 hours. Prior immediate analysis samples were decanted again and filled with two-third with dH₂O. Particle grain-size distributions were measured using a laser particle analyzer (ANALYSETTE 22 NeXT Nano; FRITSCH GmbH, Idar-Oberstein, Germany). Each sample was run in triplicate, and mean values were used for subsequent analyses.

#### 2.3.3 Data processing and analysis

Grain size data were processed using the Microsoft Excel add-on GRADISTAT Version 8.1 (Blott and Pye, 2001). For each sample, grain size distribution and the proportions of clay, silt, and sand were calculated. Sand fractions were further subdivided into five GRADISTAT-defined categories (Blott and Pye, 2001) ranging from “very fine” (63–125 µm) to “very coarse” (500–2000 µm). Sorting indices as a measure of disturbance were calculated using the Folk and Ward method (Folk and Ward, 1957) representing the standard deviation around the grain-size mean, and sorting degrees into seven categories were assigned following GRADISTAT classification (Blott and Pye, 2001) ranging from “very well sorted” (grain size deviation < 1.27 µm) to “extremely poorly sorted” (grain size deviation > 16 µm).

Statistical analyses were performed in RStudio (Posit team, 2025). The package ggpubr Version 0.6.2 (Kasambra, 2020) was used to conduct Wilcoxon rank-sum tests for unpaired samples under the assumption of non-normal data distribution. *P*-values were adjusted using the Holm–Bonferroni method and significance levels were annotated using ggpubr Version 0.6.2 (Kasambra, 2020). Effect size *r* and η² (Kruskall-Wallis test) was calculated using rstatix Version 0.7.3 (Kasambra, 2020). Figures and data tidying were performed using ggplot2 Version 3.4.1 (Whickham, 2016) and the tidyverse collection Version 2.2.0 (Whickham, 2016).

### 2.4 Population genetic analysis

#### 2.4.1 Sample collection for DNA analysis

Leaf samples were collected in the field close to the centroid for any grid cell. Leaves were placed on silica gel for rapid ultra-drying to avoid any subsequent degradation of DNA. Samples were stored accordingly on newly replaced silica gel until DNA extraction. Sample codes, sample information, elevation and exact coordinates are provided with Supplementary material S2. In total we sampled and analysed 877 plants, with 201, 457, and 219 individuals originated from Arica, Caldera, and Oyarbide, respectively. The average collection distance between individuals of about 100 m should maximize the chance to sample individuals, which did not originate from each other via clonal reproduction or via sexual offsprings. We have already shown at Oyarbide that linearly arranged *Tillandsia* show high levels of clonality along vegetation banding patterns (Koch et al., 2019), and percentages of clonal individuals over the entire study area may exceed 16 % (Jabbusch and Koch, 2025).

#### 2.4.2 DNA extraction, library preparation and genotyping-by-sequencing (GBS)

DNA extraction followed different protocols ensuring optimal results. (1) Leaf material was homogenized by grinding with a pestle in liquid nitrogen. The Invisorb Spin Plant Mini kit was used for DNA extraction following the manufacturer’s instructions (Invitek Diagnostics, Berlin, Germany). The following modifications were applied to the protocol: DNA pellets were washed twice with 70 % ethanol and then dissolved in 100 µL TE buffer (10 mM Tris-HCl, 1 mM EDTA, pH 7.5) supplemented with 2 units of RNase A; RNA digestion was performed at 37 °C for 1 hr. (2) DNA was extracted from dried leaf material using a modified version of a published CTAB protocol (Tel-zur et al., 1999), designed to improve extraction from material with high viscous polysaccharide content. The integrity of the extracted DNA was determined by running 2 μL of the final eluate on a 1 % agarose gel using GelRed staining (Merck/Sigma–Aldrich, Taufkirchen, Germany). DNA quantity was measured using a NanoDrop Spectrophotometer (ND-1000) and an Invitrogen Qubit 2.0 fluorometer kit (Thermo Fisher, Freiburg, Germany).

GBS libraries were generated involving *Pst*I and *Msp*I restriction enzymes (library size range: 150–600 bp) and were sequenced (single-end: 107 cycles, index read: 8 cycles) initially using the Illumina HiSeq2500 device (random clustering, Illumina Inc., San Diego, CA, United States) as described previously (Wendler et al., 2014). The majority of the GBS libraries was sequenced using the Illumina NovaSeq6000 instrument, which used patterned flowcells requiring a narrower size-range of the GBS libraries than the HiSeq2500 device. Therefore, the library construction protocol was adapted accordingly (library size range: 400–600 bp; Zhang et al., 2024). Sequencing of GBS libraries prepared from *Pst*I and *Msp*I fragments involved a custom sequencing primer (Wendler et al., 2014) and single-end sequencing with 122 cycles followed by two index reads (8 cycles each) according to the manufacturer’s instructions (Illumina Inc., San Diego, CA, United States). Additionally, samples were sequenced on a NovaSeq X Plus platform ((Illumina Inc., San Diego, CA, United States) differing in the library preparation from the NovaSeq6000 by using standard Illumina primers.

#### 2.4.3 GBS data analysis

GBS reads were processed using the dDocent pipeline (Puritz et al., 2014). For the assembly of a *de-novo* reference we processed raw reads of 20 randomly selected samples. We first filtered for the *Pst*1 restriction site, and second, we selected only those reads with very high sequence identity (> 0.99) and coverage (> 10) across at least six individuals. BWA (Li, 2013) mapping was done using conservative parameters (-A 2 -B 5 -O 12) and SNP calling was performed using FreeBayes (Garrison and Marth, 2012). SNPs were filtered using vcftools (Danecek et al., 2011) and vcffilter (Garrison et al., 2022), using the following parameters: no-indel, MAC 3, minQ 30, minDP 3, MAF 0.01, max-missing 0.8, min-mean-DP 5, max-mean-DP 120, AB 0.25–0.75, MQM/MQMR ratio 0.85–1.10, QUAL > 20, removed samples with > 0.3 missing data from analysis, leaving a total of 2726 SNPs across 718 individual samples. The plink software version 1.90b6.26 (Purcell et al., 2007) was used with additional filters and linkage pruning (maf 0.02, geno 0.2, indep-pairwise 50 5 0.2). After LD-pruning a total of 1127 SNPs remained. The dataset comprised a total of 72 duplicates to estimate a mean experimental error rate of the entire data set across all GBS experiments following. The genotyping error rate was calculated by counting the number of mismatched genotypes among replicates per total number of SNPs (Mastretta-Yanes et al., 2015).

A respective dataset without duplicates was used as final input data for genetic assignment tests using ADMIXTURE version 1.3.0 (Alexander et al., 2009). ADMIXTURE analysis was run for *K*=2 to 20 for the entire dataset, and optimal *K* was chosen using CV error examination. In addition, respective genetic assignment tests were also performed for all three study fields with split datasets to ensure that historical genetic differentiation processes are not biasing respective analyses. For the split datasets optimal *K* was chosen using CV error examination, too. To quantify how much genotyping errors propagate into cluster assignments, scaled Manhattan-distance between Q-vectors of duplicate samples was calculated, defined as the sum of absolute differences across all *K* ancestry components, where 0 indicates identical ancestry profiles and 1 represents the maximum possible divergence.

The following descriptive parameters describing population genetic differentiation within and among populations has been estimated: observed and expected heterozygosity, number of unique and shared alleles among study sites and the inbreeding coefficient (F_IS_) according to Nei (1987) using the hierfstat 0.5-11 R script (Goudet and Jombart, 2024). Since we have ample evidence for clonal reproduction that might affect F_IS_ calculations, we also estimated the proportion of clonal individuals within respective study fields. We defined the number of different multilocus genotypes (MLG) using the absolute genetic distance and counting the number of differing alleles between individuals with a threshold of 5 % for any single comparison. The analyses were run using the R-function mlg.filter in the poppr package (version 2.9.7) (Kamvar et al., 2014). An AMOVA was run to assign total genetic variation to between and within study field variation as well as individuals to study further the impact of clonality on the overall distribution of genetic diversity using the R-function poppr.amova in proppr (version 2.9.7) using default options (Kamvar et al., 2014). Final vcf files and R codes/scripts are accessible via Zenodo (doi: 10.5281/zenodo.18635524).

### 2.5 Climate data recording and data processing

#### 2.5.1 Set-up of weather stations

In the Caldera and Arica study sites, Onset Hobo weather stations are operated from March 2022 (Caldera) and March 2023 (Arica). The stations are aligned along elevational transects stretching between 405 m a.s.l. (MSC3) and 732 m a.s.l. (MSC1) at Caldera and between 982 m a.s.l. (WSA1) and 1098 m a.s.l. (WSA2) at Arica in order to detect elevation-related climatic differences (e.g., wind fields, fog frequency, humidity) potentially entailing changing ecological conditions. All weather stations are equipped with a HOBO U30-NRC data logger (HOBO Data Loggers, Onset Computer Corporation, Bourne, Massachusetts, USA), a relative air humidity and air temperature sensor (Onset/HOBO S-THB-M002 sensor, installed in 2 m height), a rain gauge (S-RGB-M002), a leaf wetness sensor (S-LWA-M003), and a sensor for wind speed and direction (S-WDA-M003, S-WSB-M003, 2 m height). In addition, both soil moisture and soil temperature (S-SMD-M005, S-TMB-M006) were measured 3–4 cm below *Tillandsia* stands and below adjacent bare surfaces between the *Tillandsia* dunes. A standard fog collector (SFC; Schemenauer and Cereceda, 1994) was placed in 5 m distance from each weather station. Fog water was captured by a 1 x 1 HDPE mesh exposed to dominant wind directions in 2–3 m height and was directed into a second similar rain gauge by a PVC pipe mounted below the mesh. Fog water yields were recorded in rain gauge bucket tips and converted to ml using formula developed for the specific sensor type. Logging was set to ring buffer and 15 minutes intervals to guarantee data storage for at least one year. Due to (i) cable damage by rodent or bird bites affecting several sensors of some of the weather stations, and (ii) limited accessibility, datasets are discontinuous. However, for each weather station (except WSC2), at least one year of data is available. Data were recorded and stored locally and were read out regularly on site. At Oyarbide study site, data have been obtained from two stations, OYA1a in 1128 m a.s.l. and OYA1 in 1211 m a.s.l., covering the most relevant parts of the area, related to fog-ecosystems (https://klima.rgeo.de/en/). Weather stations are linked to a DL-16 THIES data logger (THIES CLIMA, Adolf Thies GmbH & Co. KG, Göttingen, Germany), a temperature and relative air humidity sensor (Hygro-thermo transmitter compact, 1.1005.64.000, within a weather and thermal radiation shield compact, installed in 2 m height), a rain gauge (precipitation transmitter, 5.4032.35.007), a leaf wetness sensor (1.0225.50.000), and a sensor for wind speed and direction (wind transmitter compact, 4.3519.00.000/4.3129.00.000), 2 m height). Like the other weather stations, they have an SFC connected to a rain gauge (precipitation transmitter, 5.4032.35.007), measuring fog water collection volumes. The data are transferred via satellite or mobile network to a central computer in Heidelberg to enable continuous data control and near real-time data analysis.

#### 2.5.2 Data collection and processing

The stations recorded wind speeds (in ms^−1^) and wind directions every 10 min (Oyarbide) or 15 min (Arica and Caldera) in 22.5 ° classes. This results in 16 possible directions, each designated by the abbreviation of the corresponding compass point (N, E, S, W) or by a combination thereof (NNE, NE, ENE, etc.). We analysed measurements at heights of 2.0 m above the ground. All raw data (Supplementary material S3), as well as the calculations and custom R scripts (v4.3.2; R Core Team, 2021) (Supplemenary material S4) described in the following section are also available on ZENODO (doi: 10.5281/zenodo.18635524). The sand drift potential was calculated according to Fryberger and Dean (1979) and Kok et. al. (2012) based on the Lettau equation for sand drift (McKee, 1979). Following Ludwig Prandtl’s derivation (Pichler, 1986), wind speeds were also interpolated to a height of 10 m. Surface roughness factor was selected for shrub vegetation - which can be applied with 5 cm (z0 = 0.05) according to Bagnold (1954) and, for the homogeneous, smooth terrain occurring in the study areas (Wieringa, 1993). The values were grouped according to the 16 wind directions and, depending on velocity, assigned to classes. For each of these 16 categories, the drift potential was calculated using equations and respective vector units were derived for each class. The total vector units accumulated for the respective wind direction.

From the sand roses two parameters can be derived: the drift potential (DP), which represents the total amount of sand capable of movement in all directions, and the resultant drift potential (RDP), which is the sum of all vectors and thus describes the net sand transport in one prevailing direction – the resultant drift direction (RDD). The ratio of RDP to DP serves as an index of wind direction variability. Values close to 1 indicate a unimodal wind system, whereas smaller values point to increasing directional variability (Kok et al., 2012; McKee, 1979).

Meteorological time series of weather stations were aggregated into monthly and yearly means in R. Observations were classified into “day” (07:00-19:00 local time) or “night” (19:00-07:00 local time) and monthly statistics were calculated for each period and region. To ensure robust estimates, only months with high data coverage (> 25 days) were retained. Monthly values were averaged over available years to compute yearly means to avoid bias from uneven data coverage.

### 2.6 Geomorphological data analysis

#### 2.6.1 Digital terrain models and satellite data

High-resolution Digital Terrain Models (DTM) have been processed for the *Tillandsia landbeckii* study site at Arica and Caldera. The original source data have been obtained from drone flight images in 2023. The final resolution of the DTMs is about 14 cm and freely available (Stein et al., 2024a; Stein et al., 2024b). The DTM of Oyarbide has been introduced earlier (Mikulane et al., 2022) at c. 0.30 m resolution. These DTMs served as a blueprint to arrange and position the grid system over the entire area, to exactly locate samples with its coordinates and elevation, and these DTMs will serve in future studies to extract further relevant information such as vegetation cover or vegetation structure. Geomorphological data such as elevation above sea level and mean directional exposition of hillslopes used for calculations in this study has been extracted from satellite data (COPDEM GLO-30, https://doi.org/10.5270/ESA-c5d3d65) provided as Digital Surface Model (DSM) at 30 m resolution using the QGIS software (v3.34.1).

#### 2.6.2 Statistical analyses of geomorphological data

To visualize slope exposition across study sites, we constructed “exposition roses” using custom R scripts (codes and scripting available from Supplementary material S4. For these aspect values (exposition at each centroid, 30 m lateral resolution) were binned into 16 compass sectors (each 22.5°), analogous as described for the wind roses, and aggregated per study field and categorized by plant occurrence (living, dead, absent). From aspect angles of each centroid the resultant mean exposition vector for each study field was calculated using trigonometric transformation. Arrows indicating resulting vector direction are scaled in relation to the maximum sector frequency.

### 2.7 Combined statistical analyses

Results of the comparative analysis of mean exposition of sand dunes (based on 30 m grids) and mean wind direction illustrate that there is a mean effective vector along sand particles are being transported and deposited depending on wind speed (at lower and also at upper elevation of respective fields) (Hereher, 2018). Accordingly, we test if there are gradients of grain size distribution and/or sortation differences across the three study fields along this vector that may be simply attributed to elevation and respective distance being passed. If there are no such patterns, we may conclude that observed sand particle variation within a study field should be therefore the result of other interacting parameters such as presence/absence of *Tillandsia* vegetation.

To assess their relationship, pairwise Spearman correlation coefficients between geomorphological (longitude, latitude, aspect, elevation at sampling point) and sediment variables (mean grain size, sorting index, relative fractions of clay, silt and sand) have been calculated for each study site. Results were visualized as correlation ellipse plots using the R corrplot package (Wei and Simko, 2024).

## 3. Results

### 3.1 Living plant occurrence follows defined local substrate characteristics at sampling sites

To evaluate correlation between the occurrence of *T. landbeckii* and substrate characteristics, we analyzed mean grain size (µm) and the sorting index across the three sampling fields (Arica, Oyarbide, Caldera) and three categories of plant occurrence: living plants (Plant), dead plants (Dead) and plant-free sampling locations (No Plant) (Fig. 2).

**Figure 2.**
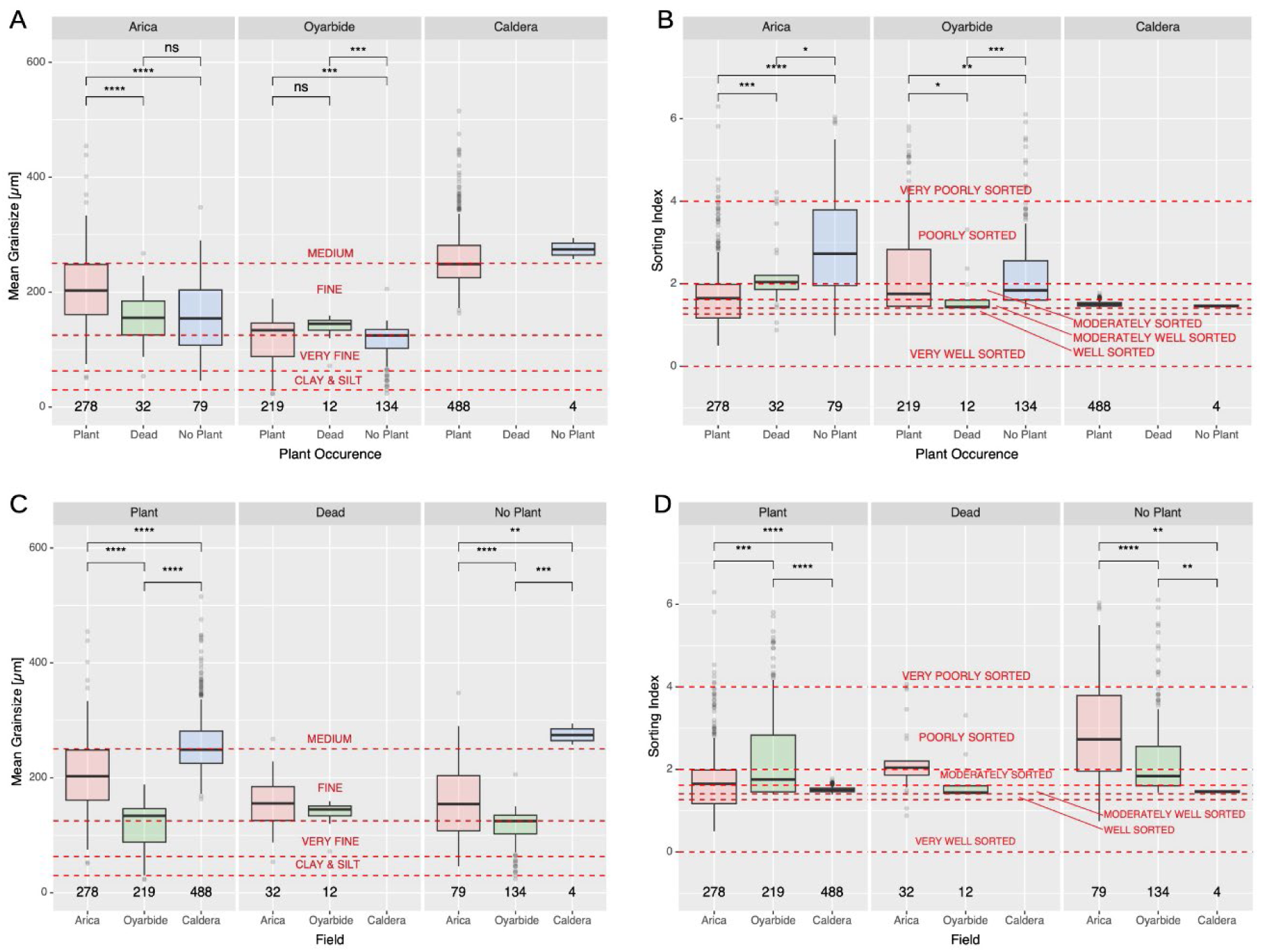
Summary statistics of grain size and sorting index distribution of substrate samples from the three study fields Arica, Oyarbide and Caldera. Data are grouped according to study fields for grain size (**a**) and sorting index (**b**), as well as according to plant occurrence (plant, dead plant, no plant) for grain size (**c**) and sorting index (**d**).

Overall, Arica showed a mean grain size of 189.8 µm and a sorting index of 2.04. Oyarbide and Caldera showed a mean grain size of 117.6 µm and a sorting index of 2.24 and 259.2 µm and 1.51, respectively. These results characterize overall substrate properties as sand of the grain size categories “very fine” (Oyarbide) and “fine” (Arica and Caldera), which are typical for wind transported (aeolian) sand, and “moderately well sorted” (Caldera), and “poorly sorted” sorting indices (Arica and Oyarbide).

Mean grain size and sorting index also varied among sites and categories of plant occurrence (Fig. 2). At Arica mean grain sizes of the substrate were significantly coarser around living plants relative to those around dead plants and no-plant areas (Fig. 2a; *p* = 3.4e-04; r = 0.2290; *p* = 2.8e-05; r = 0.2440). However, no significant difference in mean grain sizes between dead and no-plant substrates has been observed (Fig. 2a; *p* = 8.3e-01; r = 0.0204). In Caldera, the very small sample size for substrate at locations with dead or no plants limits statistically robust interpretation (Fig. 2a). In Oyarbide, mean grain size was also significantly coarser around living plants compared to plant-free sampling locations (*p* = 3.1e-03; r = 0.1790), and dead plant associated soils were also coarser than no plant substrates (*p* = 2.6e-03; r = 0.2870). No significant differences were observed between living and dead plant substrates, again statistical comparison is limited due to low sample size substrates near dead plants (*p* = 1.6e-01; r = 0.1280).

Sorting index patterns showed similar trends across the three fields (Fig. 2b). In Arica, substrates around living plants were significantly better sorted – resulting in a lower sorting index – than those around dead plants (*p* = 9.4e-04; r = 0.215) and no plants (*p* = 6.9e-14; r = 0.410). In Caldera, small sample size for dead-plant soils limited statistically robust comparison, but substrates near living plants again tended towards lower sorting indices. In Oyarbide, sorting indices were significantly lower in substrates near living plants than at plant-free sampling locations (*p* = 6.9e-03; r = 0.167), whereas substrates near dead plants are the most poorly sorted – based on limited sample size (*p* = 4.8e-02; r = 0.159; Fig. 2b).

When grouped by plant-occurrence category across fields, the same patterns persisted. Overall, substrates showed significantly coarser mean grain sizes and better sorting near living plants relative to both dead-plant and no-plant substrates aside from comparison affected by low sample size (Fig. 2c, d). Dead-plant substrates generally indicated intermediate grain sizes and sorting values, whereas plant-free substrates displayed the finest and most poorly sorted material neglecting the batch of low sample size from Caldera Fig. 2c, d). Among fields, substrates around living plants differed significantly, with Caldera showing the coarsest grain size and Iquique the finest (*p* < 0.001 across all comparisons, η² > 0.584 across all comparisons).

In summary, the results demonstrate that the presence of living *T. landbeckii* is generally associated with coarser and better-sorted aeolian sand, while in the absence of living plants finer and more poorly sorted grain-size distributions are accumulated. The strength of these trends varies between sites and are less obvious at Oyarbide and Caldera. This coincides well with the different mean grain sizes found at the three locations and transported from distant unknown regional source regions. Accordingly, trends as found at Arica may account for field-specific sand property characteristics enhancing the above-described effects.

### 3.2 Genetic data analysis highlights accelerated heterozygosity, vegetative propagation and severe spatial population structure

The estimated error rate of GBS experiments across all library preparations and sequencing experiments is 11.85 %. This value is comparatively high and does not allow detailed genetic analysis. As a result of this error rate, the mean difference between duplicate samples in ADMIXTURE cluster assignments (Manhattan-Distance) is 7.49 %. The reason may be, e.g., a change of the sequencing platform and library preparation protocols. However, comparisons across all sites and being compared with past analyses (Jabbusch & Koch, 2025; Stein et al., 2023) show the same signal and confirm robustness and reliability of the data.

Population genetic parameters demonstrate for all three populations high levels and an excess of heterozygosity. This leads to respective negative F_IS_ values. However, this is not attributed to increased outbreeding, rather than preferential clonal propagation of heterozygote individuals as highlighted by high levels of clonality between 37 % and 54 % across all three populations (Table 1). AMOVA shows a consistent finding attributing 90.8% of genetic variation within individuals and 35.3 % to genetic variation between populations. Accordingly, genetic variation between individuals within populations has a negative value of −25.9 %. This also strongly supports vegetative reproduction disturbing expectations from Hardy-Weinberg equilibrium and is also indicating a strong positive selection towards heterozygote individuals.

**Table 1.** Genetic diversity statistics for the three study fields Arica, Oyarbide and Caldera (H_e_= expected heterozygosity, H_o_=observed heterozygosity, F_IS_=inbreeding coefficient. *Multi-Locus-Genotypes (MLG) at 0.05 Hamming distance (bitwise) threshold Clonality-Index: 0 – 1: 0 = all one clone, 1 = all separate genotypes.

| population | samples | total alleles | private alleles | Shared alleles | $H_e$ | $H_o$ | $F_{IS}$ | clonality index |
| --- | --- | --- | --- | --- | --- | --- | --- | --- |
| Arica | 138 | 4033 | 519<br>(12.9 %) | 3514<br>(87.1 %) | 0.0241 | 0.0285 | -0.0331 | 75 MLG* / 138<br>= 0.543 |
| Oyarbide | 146 | 3721 | 423<br>(11.4 %) | 3298<br>(88.6 %) | 0.0554 | 0.0699 | -0.0508 | 71 MLG* / 146<br>= 0.486 |
| Caldera | 348 | 3672 | 479<br>(13.0 %) | 3193<br>(87.0 %) | 0.0258 | 0.0302 | -0.1182 | 130 MLG* /<br>348 = 0.374 |

Genetic assignment tests and visualization of spatial occurrence of genetic clusters indicates strong spatial structure (Figs. 3, 4). Irrespective of the calculation of genetic assignment using the total dataset (Fig. 3) or re-running GBS datasets individually for all study sites (Fig. 4), the spatial structure does not change indicating overall quality of genetic data. Genetic diversity is structured in all cases following an orthogonal scheme and is reproducing the linear arrangement pattern of the vegetation on rough scale. The optimal number of genetic clusters chosen using CV error examination was *K* = 7 across the entire dataset. For split data sets *K* = 3 was the optimal value for Arica and Oyarbide, and *K* = 4 for Caldera (Supplementary material S4).

**Figure 3.**
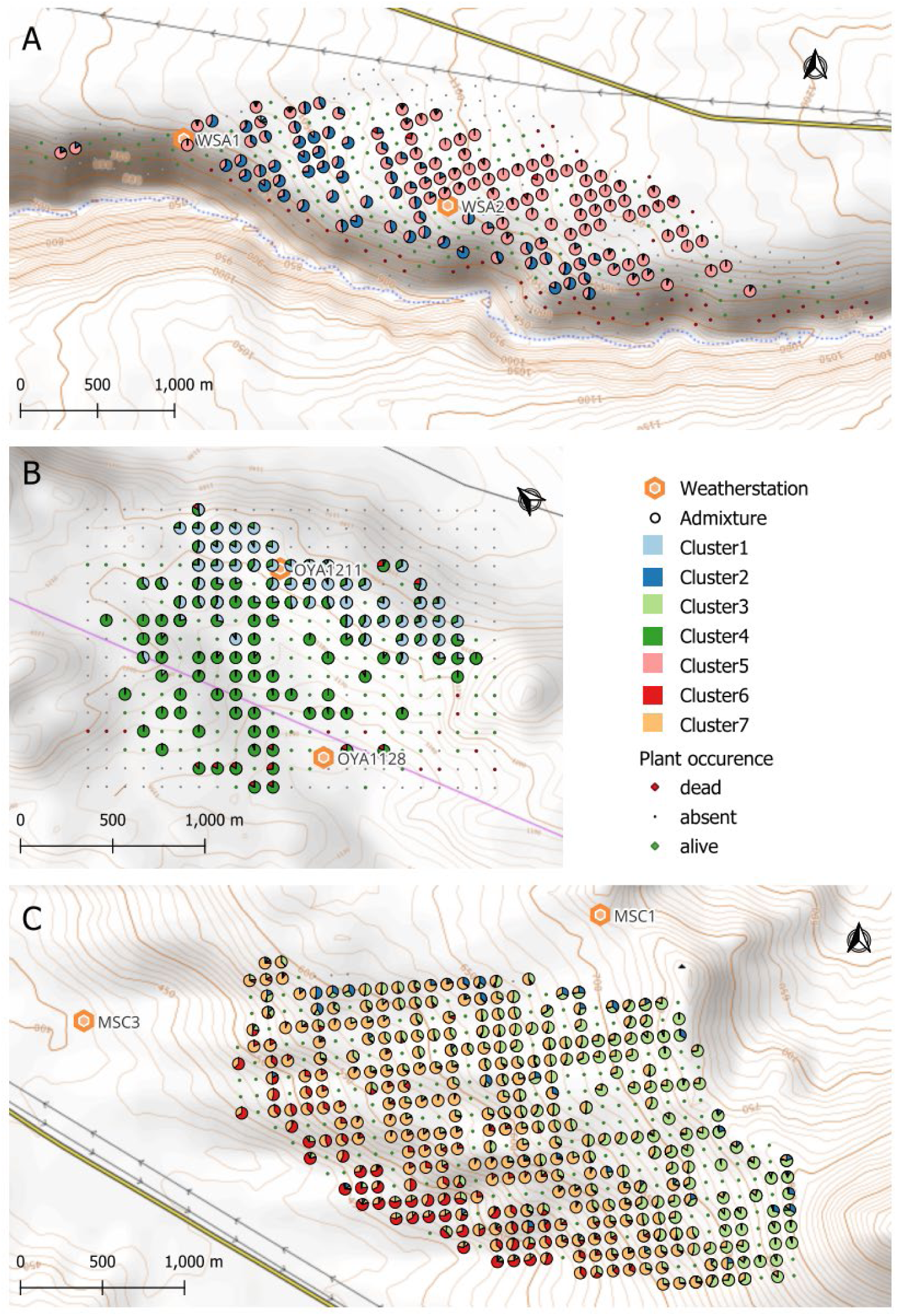
Sampling grids for substrate samples and indication of dead, alive or absent *Tillandsia*s. Assignment of genetic clusters to individuals is shown using optimal *K* = 7 for Arica (**a**), Oyarbide (**b**) and Caldera (**c**) study sites on grid scale (100 x 100m). The positions of weather stations are indicated.

**Figure 4.**
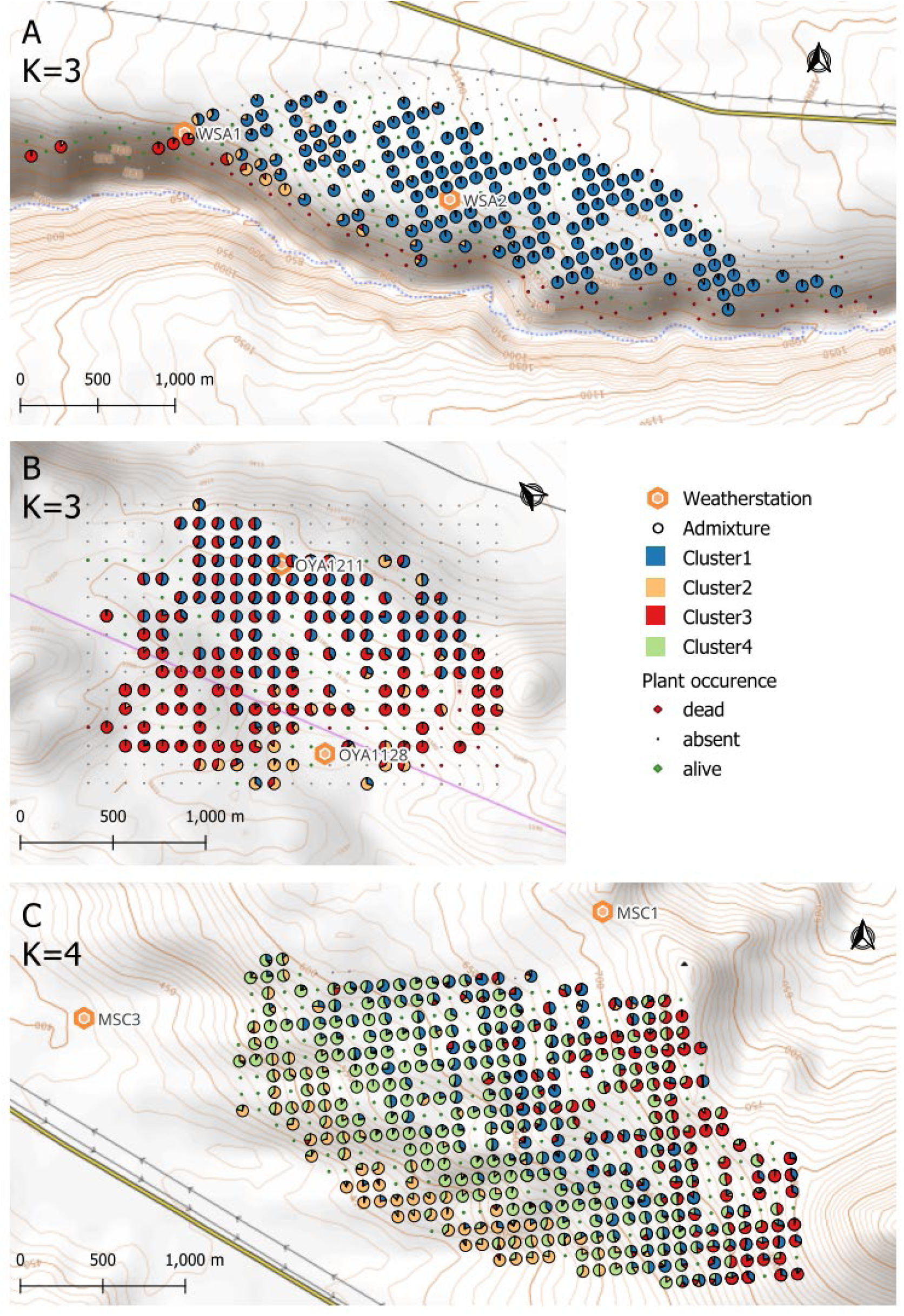
Sampling grids for substrate samples and indication of dead, alive or absent *Tillandsia*s. Assignment of genetic clusters to individuals is shown using optimal *K* = 3/4 for Arica (**a**), Oyarbide (**b**) and Caldera (**c**) study sites on grid scale (100 x 100m). The positions of weather stations are indicated.

Accordingly, we have (1) very high diversity within individuals (largely attributed to high heterozygosity), which is (2) fixed within and among populations via a preferentially vegetative reproduction mode, and (3) this genetic diversity is spatially structured as exemplified by genetic assignment tests as a result of vegetative propagation and positive selection on the respective geno- (and pheno-) types.

### 3.3 Windsystem and DTM characteristics

#### 3.3.1 Windsystem and sand drift potential

In particular, study fields at Arica and Caldera share some common features. At both sites, mean wind speed is very similar (Table 2). Wind direction as measured 2 m above ground shows also shared characteristics at Arica and Caldera. Mean wind direction is very similar among weather stations at the lower and upper range of the investigated areas, and wind systems can be classified as highly uni-modal coastal wind during daytimes. At Oyarbide we have not only stronger anabatic daytime winds, but also pronounced katabatic valley-directed winds (in particular from NNE) during nighttimes (Fig. 5, Table 2).

**Figure 5.**
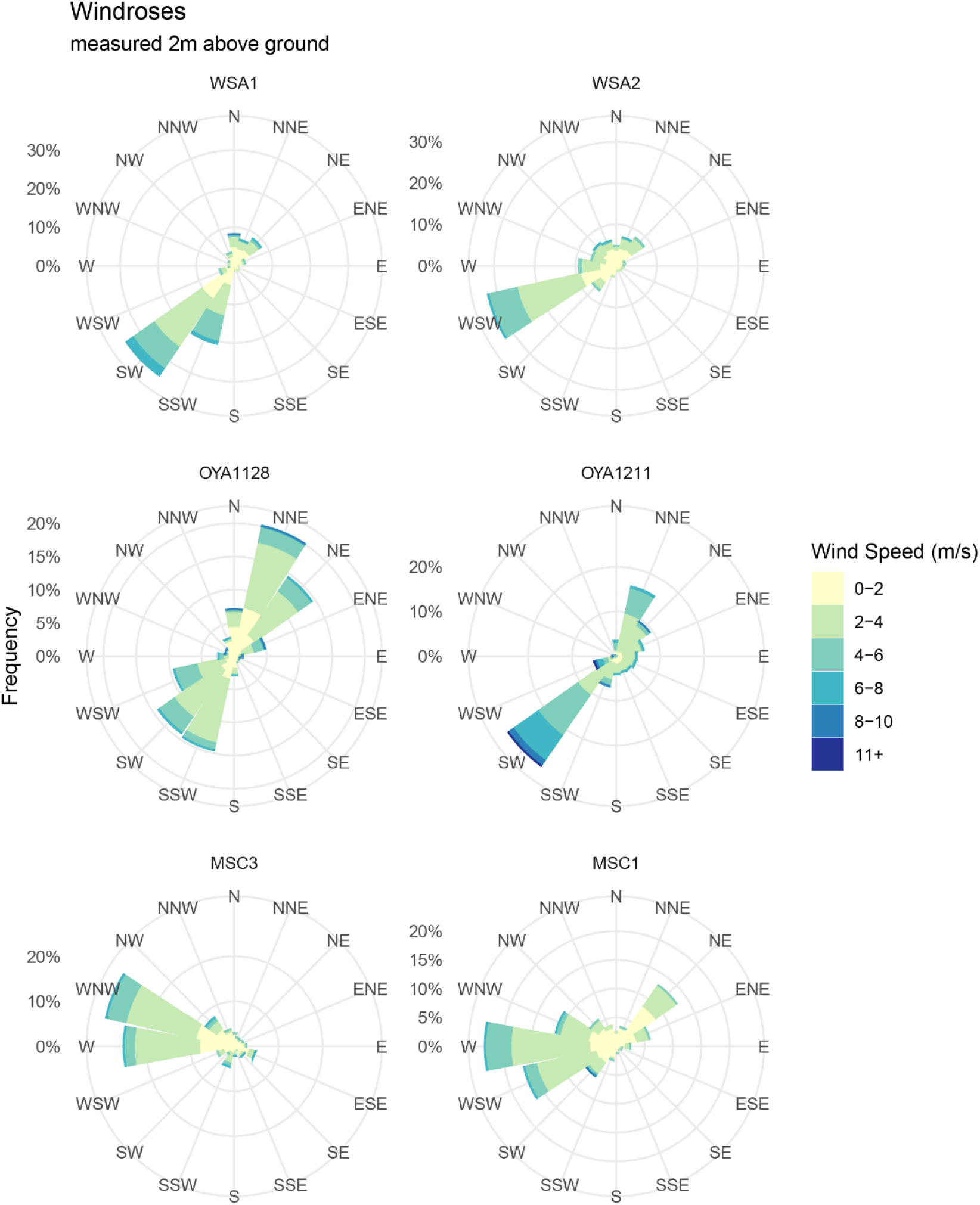
Wind speed and direction roses for real data (2 m above ground) for all six weather stations (see Figs. 3 and 4) for Arica (WSA 1 and 2), Oyarbide (OYA1128 and OYA1211) and Caldera (MSC3 and 1).

**Table 2.** Mean windspeed and temperatures aggregated by day- (7:00-19:00 local time) and nighttime (19:00-7:00 local time) per study region.

| | Mean Windspeed (Daytime, Nighttime) [$\text{ms}^{-1}$] | Mean Temperature (Daytime, Nighttime) [$^{\circ}\text{C}$] |
| --- | --- | --- |
| **Arica** | **1.4 (2.1, 0.7)** | **16.5 (18.5, 14.5)** |
| WSA1 (982 m a.s.l.) | 1.6 (2.4, 0.8) | 15.8 (17.8, 13.8) |
| WSA2 (1098 m a.s.l.) | 1.3 (2.0, 0.7) | 16.7 (18.8, 14.6) |
| **Oyarbide** | **3.1 (3.5, 2.7)** | **16.8 (19.2, 14.4)** |
| OYA1128 (1128 m a.s.l.) | 2.6 (2.9, 2.3) | 16.8 (19.2, 14.4) |
| OYA1211 (1211 m a.s.l.) | 3.6 (4.2, 3.0) | No data |
| **Caldera** | **1.5 (2.2, 0.8)** | **14.2 (16.4, 12.1)** |
| MSC3 (405 m a.s.l.) | 1.6 (2.3, 1.0) | 14.7 (17.1, 12.2) |
| MSC1 (732 m a.s.l.) | 1.4 (2.1, 0.7) | 13.8 (15.6, 11.9) |

The latter may be a result of the relatively uniform slope at Oyarbide with strong nocturnal radiation and resulting winds, whereas such winds in the two other sites are channelled in the here existing valley, where no wind data records exist. In comparison to 2 m above ground the extrapolation of wind speed at 10 m above ground resulted in accordingly higher values (Fig. 6), which have been used to calculate the various sand drift parameters (Fig. 7). We evaluated temperature information from weather stations across the three study sites and found comparable mean yearly temperatures between Arica (total average 16.5 °C, day 18.5 °C, night 14.5 °C) and Oyarbide (total average 16.8 °C, day 19.2 °C, night 14.4 °C), and lower temperatures in Caldera (total average 14.2 °C, day 16.4 °C, night 12.1 °C) (Table 2). On average, our study site temperatures are comparable to those (day 18.5 °C and night 14 °C at about 700–1350 m a.s.l.) reported earlier for the arid/hyperarid Atacama Desert (e.g. Schween et al., 2020).

**Figure 6.**
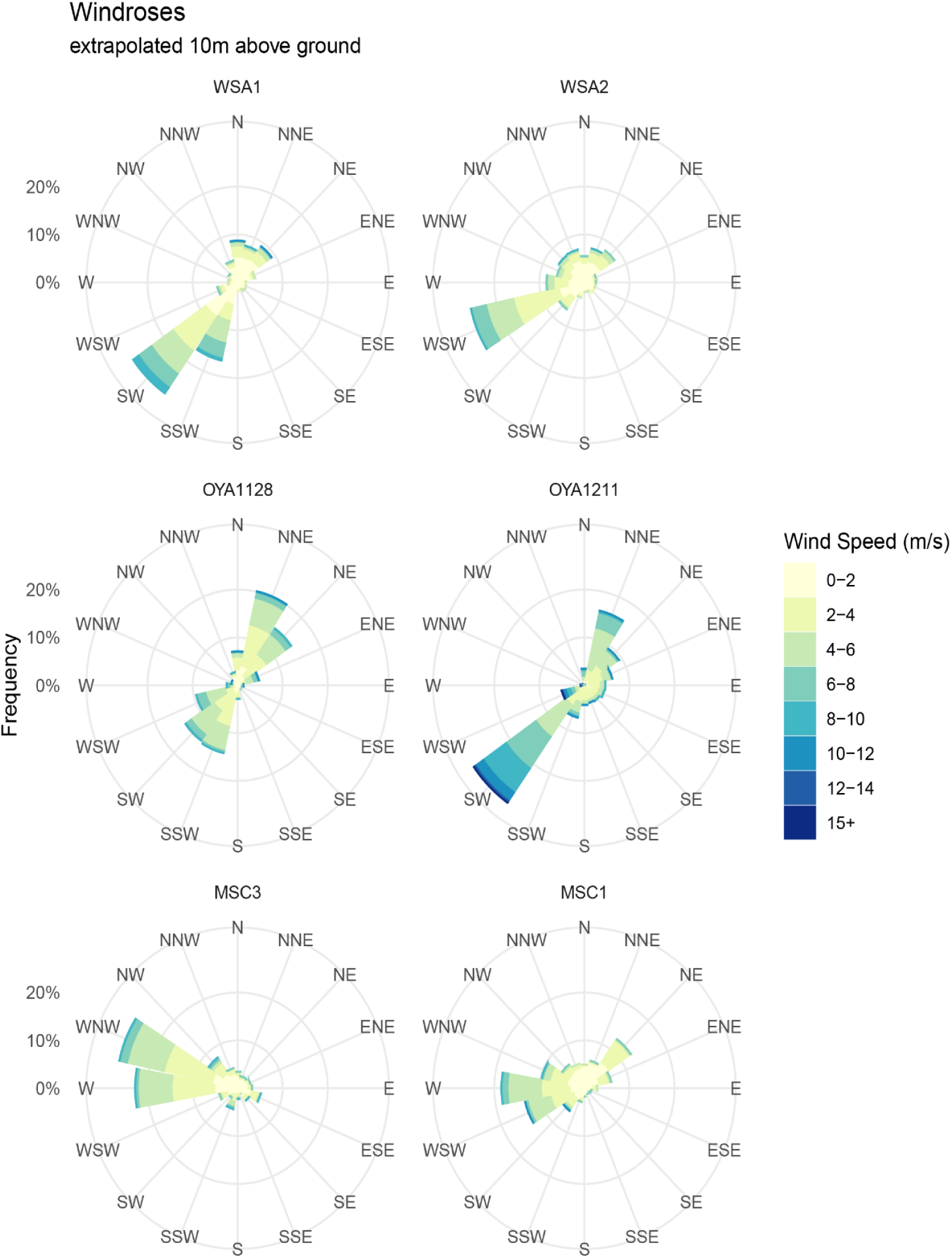
Wind speed and direction roses extrapolated to 10 m above ground for all six weather stations (see Figs. 3 and 4) for Arcia (WSA 1 and 2), Oyarbide (OYA1128 and OYA1211) and Caldeara (MSC3 and 1).

**Figure 7.**
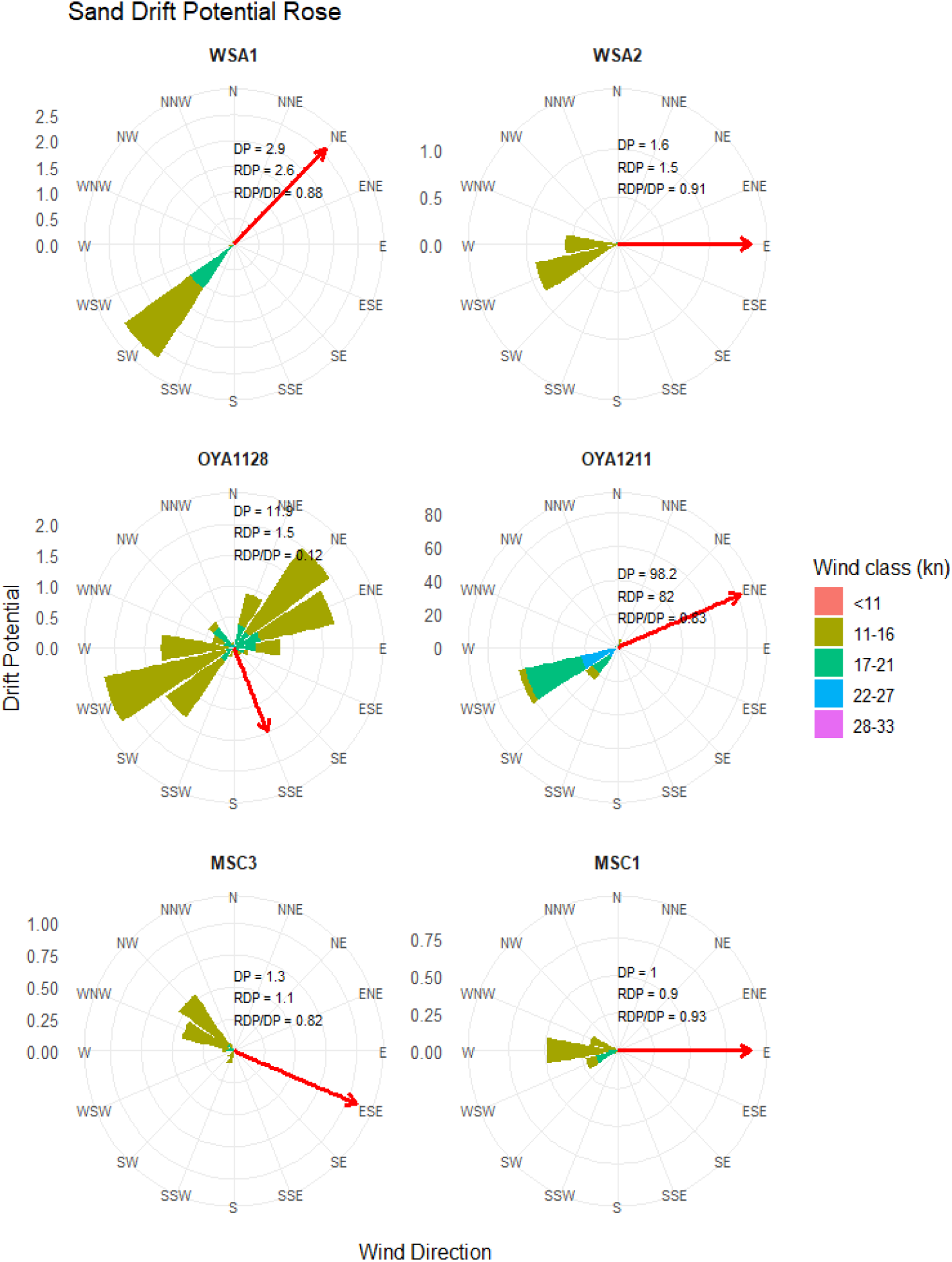
Sand drift roses extrapolated to 10 m above ground. For further comparisons with the literature wind speed is re-calculated and classified as knot speed. For all three sites DP (Drift potential), RDP (Resulting drift potential) and RDP/DB is provided as well as the resulting sand drift vector (red).

With regard to the calculated sand drift potential (Fig. 7), the three sites can be classified as low-energy systems (DP ≤ 200 VU; Fryberger and Dean, 1979). However, the existing stronger winds are able to transport sand in northeastern to eastern (Arica), south-southeastern to east-northeastern (Oyarbide), and east-southeastern to eastern direction. Only station OYA 1128 shows a low directional consistency, bidirectional wind environment (RDP/DP < 0.3). All other stations have a high directional consistency, strong unidirectional wind regime (RDP/DP ≥ 0.8).

#### 3.3.2 DTM characteristics and association with other abiotic variables

The analysis of exposition of sampling points is generally congruent and is facing mean incoming wind directions (Fig. 8). However, in particular at Caldera mean exposition is facing towards the SW and is not matching perfectly mean incoming wind direction from the W. The exposition roses also provide some good small-scaled characteristics of the terrain: Part of the dunes, which are at the wind-averted side (lee) and thereby facing inland towards ENE, are lacking *Tillandsia*s at Oyarbide (Fig. 8). This illustrates well the respective coppice dune systems and marks another difference between Arica, Caldera versus Oyarbide. Further correlations among latitudinal coordinate, altitudinal coordinate, aspect, sand grain size, sand sorting index and relative contribution of clay, sand and silt, respectively, are provided with Supplementary material S5. Across all study fields we have high and significant correlations (high r and high *p*) among all five “sand characteristics parameters” (grain size, sorting index, clay, sand, silt), which is quite typical in areas dominated by aeolian sand.

**Figure 8.**
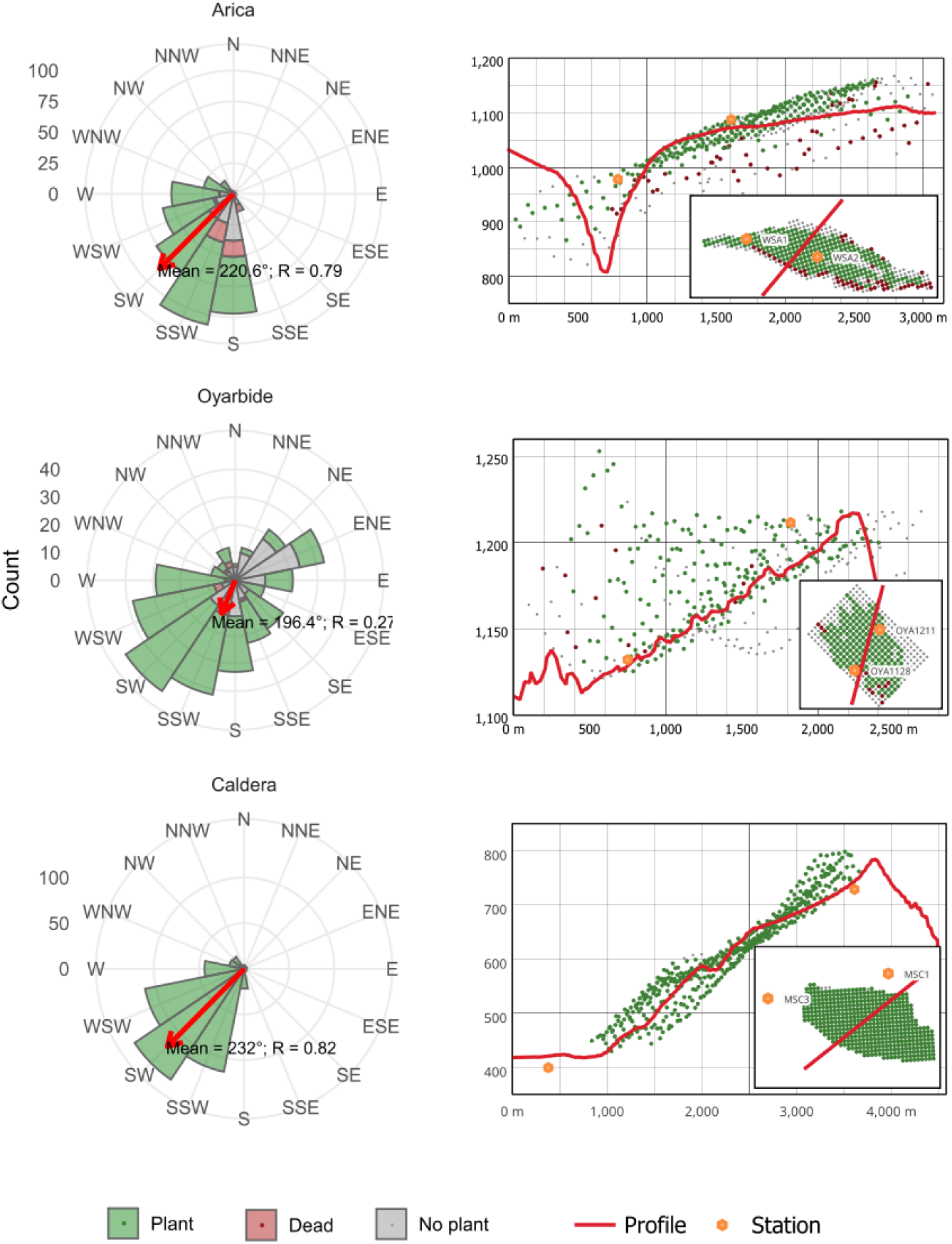
Exposition roses and profiles of the terrain. Hill exposition at each near-centroid sampling site within the 100 x 100 m grid was binned into 16 compass directions using 1 arcsecond (∼30m) resolution DSM data for the study fields at Arica, Oyarbide and Caldera. Grid-exposition characteristics described by *Tillandsia* plant occurrence are indicated (plants present, plants present but dead, no plants). Terrain profiles are given along the axis of mean exposition through the field, and orthogonal position relative to profile line (red) of weather stations (orange) and sampling points (green = live plants, red = dead remnants, grey = no plants) are indicated. Terrain profile and additional information refer also to Figs. 4 and 5.

Vice versa, high correlation is also found among the four terrain parameters (coordinates X and Y, aspect and altitude). Correlations among X, Y, elevation and aspects highlight, that sites are all facing the ocean like an inclined plane. This is best exemplified with Arica; at Oyarbide deeper coppice dune valleys are described by absence of elevation x aspect correlation, and at Caldera the inclined plane is little rotated.

However, significance and correlation value for comparisons of parameters between the two groups is often not significant, and in all cases display low correlation values only (Supplementary material S5). Accordingly, we can consider wind and sand dynamic parameters largely independent from landscape characteristics on the considered spatial scale (km²-scale with 100 x 100 m grid dimension).

## 4. Discussion

The *Tillandsia landbeckii* system in the hyperarid core of the Atacama Desert arouses curiosity but at first glance seems confusing and not very logical. *Tillandsia* individuals grow slowly with a few centimeters per year (Jabbusch and Koch, 2025; Koch et al., 2020) and reach individual ages of more than 50 years based on size calculations (Koch et al., 2020) and more than a hundred years as revealed from ^14^C dating (Jaeschke et al., 2019). Accordingly, any individual does not only face extreme conditions over time, but also has a significant risk of being confronted with rare catastrophic events resulting in extinction. That also applies to the entire vegetation, which develops into an integral, self–contained ecosystem with its geometric structures. These structures also persist over time spans of hundreds of years, as we know from ^14^C datings of fossil *Tillandsia* material (Jaeschke et al., 2024). And this long-term persistence is also true for time spans of thousands of years as indicated by a series of ^14^C dating of fossil dune remnants in present *Tillandsia* fields revealing past occurrences over time spans of more than 3.500 years (LaTorre et al., 2011). Results from Jaeschke et al. (2019) indicate that *Tillandsia* growth occurred in continuous intervals over the past 2500 yrs in the North while it ceased at ca. 1000 yrs BP in the South, in agreement with persistent dry conditions during the Medieval Climate Anomaly (Rein et al., 2004).

Accordingly, we can assume that the biotic system developed some mechanisms to reduce vulnerability to cope with generally extreme environmental conditions such as high UV radiation or limiting water availability at a critical threshold but also to compensate for stochastic fluctuations with occasional but severe impact on viability and survival such as El Niño-Southern Oscillations (ENSO) on fog and cloud formation (Del Río et al., 2021). In epiarenic *Tillandsia landbeckii* the long-lived perennial life cycle is a mechanism to minimize critical seedling phases. Accordingly, individuals can be released constantly as ramets via vegetative (clonal) propagation and are distributed across sites driven by terrain features (slope) and wind dynamics (Koch et al., 2019; Koch et al., 2020). Over time this should result in a population structure in which clonal replicates dominate by far (Jabbusch and Koch, 2025). With our results we can demonstrate this as a general principle for monospecific *Tillandsia landbeckii* vegetation. The distribution pattern of ramets is not random (Figs. 3 and 4) and reflects (i) distance, (ii) movement vectors, and (iii) available space for colonization. It has been shown earlier that on fine-scale the distribution of genetic variation at Oyarbide follows the linear growth pattern (Koch et al., 1999). With the same population Jabbusch and Koch (2025) demonstrated that ramet colonization frequency is higher at the lower part of the study field with less dense vegetation compared to the upper part. A fundamental genetic drawback of clonal propagation is a lack of gene flow among individuals, which is key to assure high genetic diversity on individual as well as on population level. Herein presented data highlights that across all investigated study fields there is a significant excess of heterozygosity (Table 1). The putative biological meaning of heterozygosity and associated increased fitness on individual (Búci et al., 2025) and population/vegetation level (Leimu et al., 2006) in *Tillandsia landbeckii* is exemplified with higher flowering frequency (Koch et al., 2019) and increased individual phenotypic variation (Jabbusch and Koch, 2025). Pollination and geneflow, of course, must occur, but its frequency is too low to be detectable in contemporary populations, e.g. while looking for seedling. However, over longer periods of time (e.g. hundreds to thousands of years) these rare events can be realized and, importantly, any successful offspring can be propagated clonally afterwards and will again contribute to overall diversity. This is best exemplified by *Tillandsia* vegetation with sympatrically occurring epiarenic *Tillandsia* species. Interspecies and adaptive gene flow between *T. landbeckii* and *T. capillaris* in Peru resulted in new geno- and phenotypes at non-random spatial distribution and enlarged population range contributing eventually to increased vegetation robustness at its margins (Stein et al., 2025). Furthermore, past Pleistocene range fluctuations forced populations into secondary contact and may have also fostered periodic episodes of increased outbreeding (Koch et al., 2022).

Wind and sand dynamics are assumed to play a key role in occurrence and persistence of *Tillandsia landbeckii* ecosystems. They provide the substrate for anchoring the rootless plants (Koch et al., 2019), and they transport the fog from shore distances of in average 15 km (Wolf et al., 2016). However, wind speed and high kinetic energy also causes some severe stress in respect to evaporation (water loss) and mechanical damage (deflation/abrasion, and/or sand burial).

We have evidence that fog occurrence or the amount of available fog within a *Tillandsia* field have a pronounced effect on *Tillandsia* growth phenotypes, and accordingly fine-scaled correlating patterns have been observed along clinal gradients (e.g. Jabbusch and Koch, 2025). We do not see this spatial effect in such a pronounced way for sand characteristics, grain size and sortation. All three areas studied show average grain sizes in the fraction preferentially moved by wind (< 250 µm). Furthermore, the overall poor sorting grades indicate both a comparatively short transport distance for the sediments and a low sorting influence of the *Tillandsia* populations. There is some correlation of topographic variables (coordinates X and Y, aspect, and elevation) with grain size and sortation in all of the three fields (Supplementary material S5). However, the responses are not congruent and overall they are study site specific and reflect best differing topology. On the contrary, we observe a largely homogeneous distribution of sand grain sizes and sortation associated with present and living *Tillandsia* plants. However, these grain sizes (and its mean) differ significantly among study sites (Fig. 2c), which simply reflects the different sand sources, and we can conclude that any sand grain size in a range from 117–260 µm is suitable for *Tillandsia* growth performance. Two single push cores from Arica and Caldera with 100 cm depth each also highlight quite stable sorting indices of around 1.8–2.0 over long timescales (Rennemüller, 2023). These push cores can be considered to span c. 500–1000 years back in time. However, the profiles examined also show sections where Tillandsia populations were initially eroded by sand winds and then covered by sterile sand. This demonstrates the sensitivity of the systems to climatic changes (Cerceda et al., 2008). Along this temporal transect sand grain size shows higher variation at around 200 µm at Arica (in agreement with our mean of 189 µm) and 280 µm at Caldera (in agreement with our mean of 260 µm). This illustrates well a long-term persistent sand source and stable wind system. In summary, our findings highlight the tremendous importance of sand as an abiotic factor and evolutionary parameter leading to the convergent/parallel evolution of nine different epiarenic Tillandsia species found only in the Chilean-Peruvian coastal desert system (Koch et al., 2026).

Do *Tillandsia*s shape and structure their own environment? We may answer the question with yes, because we often observe the regular growth banding patterns orthogonally arranged to incoming wind and fog and following contour lines mirroring aspect. This pattern formation has been analysed in detail in the past from a mechanistic and topographic perspective, and mostly did not consider *Tillandsia* biotic feedback loops and presence and absence of plants (Gandhi et al., 2018, Kästner et al., 2025). A recent study considered non-reciprocal feedback loops acting on a smaller scale using mathematical modeling (Hidalgo-Ogalde et al., 2024), that may explain varying banding patterns for example. Accordingly, it is important to conclude from our study that *Tillandsia* have not only significant effect on the process of trapping sand, but also sand sortation is affected and differs among sites with and without *Tillandsia*s. In addition to this kind of biotic-abiotic feedback loop, there are also biotic-biotic interactions between *Tillandsia* and diversity of microbiota also with severe coincidence with banding patterns and presence/absence of *Tillandsia* plants (Alfaro et al., 2021; Hakobyan et al., 2023; Jaeschke et al., 2024); and recently, specific nitrogen-fixing bacterial communities in the phyllosphere of *Tillandsia* potential source of nitrogen acquisition have been described (Lyu et al., 2024), which may have a direct effect on growth, which has to be fine-tuned to balance sand dynamics and fog perception.

Generally, we can conclude that *Tillandsia landbeckii* found its ecological niche in a “low wind energy system” in conditions where mean wind at 10 m height is below about 2 m/s and where the kinetic energy flux in the wind is very small. However, temporarily kinetic energy is still sufficient for aeolian sand transport over regional distances, and wind direction fosters fog transport from the coastal line and guarantees sufficient fog events as water supply. Other and more general topographic characteristics of the three study fields (Arica, Oyarbde, Caldera) stretching the entire distribution of *Tillandsia landbeckii* in Chile are in congruence with earlier and detailed calculations from satellite images and covering a large area in the Iquique region (Wolf et al., 2016; 95 % range): elevation ranges c. 1000–1120 m a.s.l. (our mean 970 m a.s.l.), aspect c. 160–270° (our mean 216°), distance from coast ranges from 12–21 km (our mean c. 19 km), and slope ranges from 6–13°. Alongside above-described parameters also temperature at day and in particular at night plays an important role to assure physiological integrity. *Tillandsia landbeckii* exhibit a special type of photosynthesis (CAM: Crassulaceae acid metabolism) separating CO_2_ fixation and accumulation of C_4_-carbonic acids at night and light reaction and carbon reduction (sugar production) during the day. CAM allows it to cope efficiently with limited water availability. CAM photosynthesis responds differently to temperature compared to C_3_ and C_4_ photosynthesis and it is operating optimal at lower temperatures (Heyduk, 2022; Lin et al., 2006; Yamori et al., 2014) with lower night temperatures also preventing C_4_-acid leakage from the vacuole (Friemert et al., 1988; Lüttge, 2004). At all three study sites temperature regimes are ideal with moderate temperature at day and mean night temperatures at about 12–14.5 °C (Table 2).

Over recent decades, fog-dependent ecosystems along the Chilean-Peruvian coast have exhibited increasing signs of decline, potentially associated with abrupt mesoscale climatic changes since the mid-1970s (Schulz et al., 2011), suggesting pronounced negative population trends. Yet, both the magnitude of this decline and the ecological–atmospheric mechanisms driving it remain insufficiently understood (Latorre et al., 2011). The few existing reports are alarming and describe high mortality of *Tillandsia* in the coastal lomas (Cereceda et al., 2008; Rundel et al., 1997); however, the spatial extent and specific characteristics of this die-back, as well as the role of altered fog regimes in loma degradation, are still unclear (Cereceda et al., 2008).

Past historical dieback has been reported for multiple phases of dune growth with alternating plant colonization, dieback, and sand accumulation during the past ∼1300 years and spanning the Medieval Climate Anomaly (Jaeschke et al., 2024). *Tillandsia* dieback of the past 50 years is likely the result of a long-lasting and major global change affecting Chilean coastal climate in general (Cai et al., 2014; Del Rio et al., 2018). A survey of sites from the central cluster of *T. landbeckii* in Chile around Iquique from 1955 to 1997 revealed a total loss of living *T. landbeckii* lomas of about 36 % (Osses et al., 2007). A general vegetation dieback in northern coastal Chile is associated with a total decrease of low cloud cover at Arica in the second half of the twentieth century and similar trends are shown for Iquique and Copiapó (closer to Caldera) throughout the 20th century (Schulz et al., 2011). Rundel et al. (1997) stated for the lquique region that a decline in abundance is taking place, albeit gradually. For *Tillandsia* loma vegetation it is in particular the orographic fog accounting for only 22 % of fog occurrence (Keim-Vera et al., 2024).

At our study sites it is in particular the Arica population at the lower elevation margin (Fig. 8) at steeper slopes towards the valley (Quebrada del Diablo), which is facing some severe dieback. It may indicate that the fog-system underwent changes during the past decades. Maybe this is fostered by topological constraints and ridge effects on wind speed dynamics. In Oyarbide there is some dieback at the lower margin (Fig. 8). Here vegetation is sparse and plants are smaller with significantly lower growth rates (Jabbusch and Koch, 2025; Koch et al., 2020). Accordingly, we may consider these lower margins as a typical abrupt ecocline zone towards unfavourable conditions, where only few genetically and phenotypically specialized individuals persist.

## 5. Conclusion

*Tillandsia landbeckii* forms a remarkably long–lived, clonally structured fog–dependent ecosystem that persists for millennia by combining rare but crucial genetic recombination with continuous vegetative spread, finely tuned to low–energy wind, stable sand supply, and narrow atmospheric and fog regimes – yet this finely balanced system is now increasingly destabilized by climate–driven changes in fog dynamics, triggering pronounced dieback at its distribution margins.

## Code availability

Codes and scripts for any analyses are compiled with Supplementary material File 4. Which can be accessed in this DOI link: https://doi.org/10.5281/zenodo.18635525 (Koch et al., 2026).

## Data availability

All accession and sample details, and all results can be accessed in this DOI link: https://doi.org/10.5281/zenodo.18635525 (Koch et al., 2026).

## Sample availability

Samples are stored at Heidelberg Herbarium (HEID) and can be accessed if needed, please reach the author for more information.

## Supplement

Supplementary material S1 to S5 related to this article is available online at https://doi.org/xxxxxxxxxxxxxxxxxxxxxxxx.

## Author contributions

Marcus A. Koch: Conceptualization, Data curation, Formal analysis, Funding acquisition, Investigation, Methodology, Project administration, Resources, Supervision, Validation, Visualization, Writing – original draft, Writing – review and editing. Robin Schweikert: Data curation, Formal analysis, Investigation, Methodology, Software, Writing – review and editing. Ron Eric Stein: Data curation, Formal analysis, Investigation, Methodology, Software, Writing – review and editing. Neil Bogs: Methodology. Olaf Bubenzer: Methodology, Writing – original draft, Writing – review & editing. Camilo del Río: Methodology, Writing – review & editing. Dörte Harpke: Investigation. Simon Matthias May: Data curation, Methodology. Alexander Siegmund: Data sharing, Writing – review & editing. Alexandra Stoll: Writing – review & editing. Dietmar Quandt: Funding acquisition, Writing – review & editing.

## Disclaimer

Publisher’s note: Copernicus Publications remains neutral with regard to jurisdictional claims made in the text, published maps, institutional affiliations, or any other geographical representation in this paper. The authors bear the ultimate responsibility for providing appropriate place names. Views expressed in the text are those of the authors and do not necessarily reflect the views of the publisher.

## Acknowledgements

We would like to thank our colleagues in the Collaborative Research Center (CRC) 1211 for ensuring the project was executable in such comfortable conditions and Heidelberg University of Education and Pontificia Universidad Católica de Chile for providing the necessary climate data for Oyarbide from their long-term comprehensive fog ecosystem monitoring program there. We thank Max Engel, Luana Anders and Alina Peters for technical support in the sand analytical laboratory at Heidelberg university, Peter Sack (Heidelberg Botanical Garden and Herbarium collections) for sample curation, Christiane Kiefer and Sarina Jabbusch (Heidelberg) for GBS data analyses support and Sarina Jabbusch, Julia Bechteler and Paloma Morales for assisting sampling in the field.

## References

Alexander, D.H., Novembre, J., and Lange, K.: Fast model-based estimation of ancestry in unrelated individuals, Genome Research, 19(9), 1655–1664, 10.1101/gr.094052.109 2009.

Alfaro, F., Manzano, M., Almiray, C., Garcia, J.L., Osses, P., Del Rio, C., Vargas, C., Latorre, C., Koch, M.A., Siegmund, A., and Abades, S.: Soil bacterial community structure of fog-dependent Tillandsia landbeckii dunes in the Atacama Desert, Plant Systematics and Evolution, 307, e56, 10.1007/s00606-021-01781-0, 2021.

Anderegg, W.R.L., Klein, T., Bartlett, M., Sack, L., Pellegrini, A.F.A., Choat, B., and Jansen, S.: Meta-analysis reveals that hydraulic traits explain cross-species patterns of drought-induced tree mortality, Proceedings of the National Academy of Sciences of the United States of America, 113(18), 5024–5029, 10.1073/pnas.1525678113 2016.

Arens, F. L., Airo, A., Feige, J., Sager, C., Wiechert, U., and Schulze-Makuch, D.: Geochemical proxies for water-soil interactions in the hyperarid Atacama Desert, Chile, Catena, 206, 105531, 10.1016/j.catena.2021.105531 2021.

Bagnold, R.A.: The physics of blown sand and desert dunes, Methuen & Co., London, UK, 265 pp., ISBN 9780486439310, 1954.

Blott, S.J., and Pye, K.: GRADISTAT: A grain size distribution and statistics package for the analysis of unconsolidated sediments, Earth Surface Processes and Landforms, 26(11), 1237–1248, 10.1002/esp.261 2001.

Búci, M., Jarčuška, B., Klinga, P., Ružinská, R., Berggren, A., and Kaňuch, P.: Individual heterozygosity and fitness in bottlenecked populations during early colonisation, Biological Invasions 27, 111, 10.1007/s10530-025-03571-y, 2025.

Cai, W., Borlace, S., Lengaigne, M., van Rensch, P., Collins, M., Vecchi, G., Timmermann, A., Santoso, A., McPhaden, M. J., Wu, L., England, M. H., Wang, G., Guilyardi, E., and Jin, F.-F.: Increasing frequency of extreme El Niño events due to greenhouse warming, Nature Climate Change, 4, 111–116, 10.1038/nclimate2100, 2014.

Chiappero, M.F., Rossetti, M.R., Moreno, M.L., Pérez-Harguindeguy, N.: A global meta-analysis reveals a consistent reduction of soil fauna abundance and richness as a consequence of land use conversion, Science of the Total Environment, 946, 173822, 10.1016/j.scitotenv.2024.173822 2024.

Cubino, J.P., Chytrý, M., Divíšek, J., Jiménez-Alfaro, B.: Climatic filtering and temporal instability shape the phylogenetic diversity of European alpine floras, Ecography, 2022, e06316, 10.1111/ecog.06316 2022.

Danecek, P., Auton, A., Abecasis, G., Albers, C.A., Banks, E., DePristo, M.A., Handsaker, R.E., Lunter, G., Marth, G.T., Sherry, S.T., McVean, G., Durbin, R.: 2011. The variant call format and VCFtools, Bioinformatics, 27(15), 2156–2158, 10.1093/bioinformatics/btr330 2011.

Del Río, C., Carter, V., Lobos-Roco, F., Astaburuaga, J.P., Qüense, J., Rivera, D., Espinoza, V., Anabalón, G., Bonet, M., Osses, P., and Contreras, C.: Mapa de agua de niebla (18°–35°S): Modelo de disponibilidad y gobernanza de una fuente complementaria para territorios de escasez hídrica [interactive resource], Centro UC Desierto de Atacama, Pontificia Universidad Católica de Chile, 10.7764/CDAUC.BD.01, 2025.

Del Rio, C., Garcia, J. L., Osses, P., Zanetta, N., Lambert, F., Rivera, D., Lobos, F.: ENSO influence on coastal fog-water yield in the Atacama Desert, Chile, Aerosol Air Quality Research, 18, 127–144, 10.4209/aaqr.2017.01.0022, 2018.

Del Rio, C., Lobos-Roco, F., Latorre, C., Koch, M.A., Garcia, J.L., Osses, P., Lambert, F., Alfaro, F., Siegmund, A.: Spatial distribution and interannual variability of coastal fog and low-cloud cover in the hyperarid Atacama Desert and implications for past and present Tillandsia landbeckii ecosystems, Plant Systematics and Evolution, 307, e58, 10.1007/s00606-021-01782-z, 2021a.

Del Río, C., Lobos, F., Siegmund, A., Tejos, C., Osses, P., Huaman, Z., Meneses, J. P., García, J.-L.: GOFOS, Ground Optical Fog Observation System for monitoring the vertical stratocumulus-fog cloud distribution in the coast of the Atacama Desert, Chile, Journal of Hydrology, 97, 126190, 10.1016/j.jhydrol.2021.126190 2021b.

Folk, R.L., Ward, W.C.: Brazos River bar: A study in the significance of grain size parameters, Journal of Sedimentary Petrology, 27, 3–26, 10.1306/74D70646-2B21-11D7-8648000102C1865D, 1957.

Frenette-Dussault, C., Shipley, B., Hingrat, Y.: 2012. Functional structure of an arid steppe plant community reveals similarities with Grime’s C–S–R theory, Journal of Vegetation Science, 23(2), 208–222, 10.1111/j.1654-1103.2011.01350.x 2012.

Friemert, V., Heininger, D., Kluge, M., Ziegler, H.: Temperature effects on malic-acid efflux from the vacuoles and on the carboxylation pathways in crassulacean-acid-metabolism plants, Planta 174, 453–461, 10.1007/BF00634473, 1988.

Fryberger, S.G., Dean, G.: Dune forms and wind regime, in: A study of global sand seas, edited by: McKee, E.D., U.S., Geological Survey Professional Paper 1052, pp. 137–169, 1979.

Gandhi, P., Werner, L., Iams, S., Gowda, K., Silber, M.: A topographic mechanism for arcing of dryland vegetation bands, Journal of the Royal Society Interface, 15(147), 20180508, 10.1098/rsif.2018.0508 2018.

Garcia, J.L., Lobos-Roco, F., Schween, J.H., Del Rio, C., Osses, P., Vives, R., Pezoa, M., Siegmund, A., Latorre, C., Alfaro, F., Koch, M.A., Löhnert, U.: Climate and coastal low-cloud weather dynamics in the hyperarid Atacama fog desert and geographic distribution of Tillandsia landbeckii (Bromeliaceae) dune ecosystems, Plant Systematics and Evolution, 307, e57, 10.1007/s00606-021-01775-y, 2021.

Garrison, E., Marth, G.: Haplotype-based variant detection from short-read sequencing. arXiv [preprint], 10.48550/arXiv.1207.3907, 2012.

Garrison, E., Kronenberg, Z.N., Dawson, E.T., Pedersen, B.S., Prins, P.: A spectrum of free software tools for processing the VCF variant call format: vcflib, bio-vcf, cyvcf2, hts-nim and slivar, PLOS Computational Biology, 18, e1009123, 10.1371/journal.pcbi.1009123, 2022.

Goudet, J.,Jombart, T.: hierfstat: Estimation and tests of hierarchical F-statistics, R package version 0.5–11, github [code], https://github.com/jgx65/hierfstat, 2024.

Goudie, A.: Desert Landscapes of the World with Google Earth, Springer Cham, Switzerland, 271 pp., 10.1007/978-3-031-15179-8, 2022.

Grime, J.P.: Evidence for the existence of three primary strategies in plants and its relevance to ecological and evolutionary theory, The American Naturalist, 111(982), 1169–1194, 10.1086/283244 1977.

Hakobyan, A., Velte, S., Sickel, W., Quandt, D., Stoll, A., and Knief, C.: Tillandsia landbeckii phyllosphere and laimosphere as refugia for bacterial life in a hyperarid desert environment, Microbiome, 11, 246, 10.1186/s40168-023-01684-x, 2023.

Heyduk, K: Evolution of Crassulacean acid metabolism in response to the environment: past, present, and future, Plant Physiology, 190, 19–30, 10.1093/plphys/kiac303, 2022.

Hereher, M.E.: Geomorphology and drift potential of major aeolian sand deposits in Egypt, Geomorphology, 304, 113–120, 10.1016/j.geomorph.2017.12.041, 2018.

Hidalgo-Ogalde, B., Pinto-Ramos, D., Clerc, M.G., Tlidi, M.: Nonreciprocal feedback induces migrating oblique and horizontal banded vegetation patterns in hyperarid landscapes, Scientific Reports, 14, 14635, 10.1038/s41598-024-63820-3 2024.

Ihaddaden, A., Velázquez, E., Rey-Benayas, J.M., and Kadi-Hanifi, H.: Climate and vegetation structure determine plant diversity in Quercus ilex woodlands along an aridity and human-use gradient in Northern Algeria, Flora, 208(4), 268–284, 10.1016/j.flora.2013.03.009, 2013.

Jabbusch, S., and Koch, M.A.: *Tillandsia landbeckii* secures high phenotypic plasticity via clonal propagation at the dry limits of plant life in the Atacama Desert, Perspectives in Plant Ecology, Evolution and Systematics, 66, 125846, 10.1016/j.ppees.2025.125846 2025.

Jaeschke, A., Böhm, C., Merklinger, F. F., Reyers, M., Kusch, S., Rethemeyer, J., Bernasconi, S.: Variation in δ15N of fog-dependent Tillandsia ecosystems reflect water availability across climate gradients in the hyperarid Atacama Desert, Global and Planetary Change, 183, 1–12, 10.1016/j.gloplacha.2019.103029, 2019.

Jaeschke, A., May, S.M., Hakobyan, A., Mörchen, R., Bubenzer, O., Bernasconi, S.M., Schefuß, E., Hoffmeister, D., Latorre, C., Gwozdz, M., Rethemeyer, J., Knief, C.: Microbial hotspots in a relict fog-dependent *Tillandsia landbeckii* dune from the coastal Atacama Desert, Global and Planetary Change, 234, 104383, 10.1016/j.gloplacha.2024.104383 2024.

Jaime, X.A., Angerer, J.P., Fuhlendorf, S.D., Walker, J.W., Yang, C., Tolleson, D.R., and Wu, X.B.: Effects of prescribed fire on plant alpha- and beta-diversity and the regulating role of soil in a mesquite–oak savanna, Landscape Ecology, 40, 233, 10.1007/s10980-025-02259-x, 2025.

Jürgens, N., Schmiedel, U., and Finckh, M.: Fairy circles of the Namib Desert: Ecosystem engineering by subterranean social insects, Biodiversity and Ecology, 7, 1–376, 10.7809/b-e.vol_07, 2022.

Kamvar, Z.N., Tabima, J.F., and Grünwald, N.J.: Poppr: An R package for genetic analysis of populations with clonal, partially clonal, and/or sexual reproduction, PeerJ, 2, e281, 10.7717/peerj.281, 2014.

Kassambara, A.: ggpubr: “ggplot2” based publication ready plots, R package, CRAN [code], https://CRAN.R-project.org/package=ggpubr, 2020.

Kassambara, A.: rstatix: Pipe-friendly framework for basic statistical tests, R package, CRAN [code], https://CRAN.R-project.org/package=rstatix, 2023.

Kästner, K., Caviedes-Voullième, D., and Hinz, C.: Formation of spatial vegetation patterns in heterogeneous environments, PLOS ONE, 20(5), e0324181, 10.1371/journal.pone.0324181 2025.

Keim-Vera, V., Lobos-Roco, F., Aguirre, I., Merino, C., and del Río, C.: Fog types frequency and their collectable water potential in the Atacama Desert, Atmospheric Research, 312, 107747, 10.1016/j.atmosres.2024.107747 2024.

Koch, M.A., Kiefer, C., Möbus, J., Quandt, D., Merklinger, D., Harpke, D. and Villasante Benavides, F.: Range expansion and contraction of *Tillandsia landbeckii* lomas in the hyperarid Chilean Atacama Desert indicates ancient introgression and gene flow, Perspectives in Plant Ecology, Evolution and Systematics, 56, 125689, 10.1016/j.ppees.2022.125689 2022.

Koch, M.A., Kleinpeter, D., Auer, E., Siegmund, A., del Rio, C., Osses, P., Garcia, J.L., Marzol, M.V., Zizka, G., and Kiefer, C.: Living at the dry limits: Ecological genetics of *Tillandsia landbeckii* lomas in the Chilean Atacama Desert, Plant Systematics and Evolution, 305, 1041–1053, 10.1007/s00606-019-01623-0 2019.

Koch, M., Schweikert, R., and Stein, R. E.: Drowning in a sandy ocean: Epiarenic growth of Tillandsia in the hyperarid Atacama Desert, ZENODO [data set], 10.5281/zenodo.18635525, 2026.

Koch, M.A., Stock, C., Kleinpeter, D., Del Rio, C., Osses, P., Merklinger, F.F., Quandt, D., Siegmund, A.: Vegetation growth and landscape genetics of *Tillandsia* lomas at their dry limits in the Atacama Desert show fine-scale response to environmental parameters, Ecology and Evolution, 10, 13260–13274, 10.1002/ece3.6924 2020.

Kok, J.F., Parteli, E.J.R., Michaels, T.I., and Karam, D.B.: The physics of wind-blown sand and dust, Reports on Progress in Physics, 75(10). 106901, 10.1088/0034-4885/75/10/106901, 2012.

Latorre, C., González, A.L., Quade, J., Fariña, J.M., Pinto, R., and Marquet, P.A.: Establishment and formation of fog-dependent *Tillandsia landbeckii* dunes in the Atacama Desert: Evidence from radiocarbon and stable isotopes, Journal of Geophysical Research, 116, G03033, 10.1029/2010JG001521 2011.

Leimu, R., Mutikanen, P., Koricheva, J., and Fischer, M.: How general are positive relationships between plant population size, fitness and genetic variation?, Journal of Ecology, 94, 942–952, 10.1111/j.1365-2745.2006.01150.x, 2006.

Lenné, J., and Wood, D.: Monodominant natural vegetation provides models for nature-based cereal production, Outlook on Agriculture, 51(1), 11–21, 10.1177/00307270221078022, 2022.

Li, H.: Aligning sequence reads, clone sequences and assembly contigs with BWA-MEM, arXiv [preprint], arXiv:1303.3997, 10.48550/arXiv.1303.3997, 2013.

Li, J., and Prentice, I.C.: Global patterns of plant functional traits and their relationships to climate, Communications Biology, 7, 1136, 10.1038/s42003-024-06777-3 2024.

Lin, Q., Abe, S., Nose, A., Sunami, A., and Kawamitsu, Y.: Effects of High Night Temperature on Crassulacean Acid Metabolism (CAM) Photosynthesis of Kalanchoë pinnata and Ananas comosus, Plant Production Science, 9(1), 10–19. 10.1626/pps.9.10. 2016.

Liu, C.K.C., Kuchma, O., and Krutovsky, K.V.: Mixed-species versus monocultures in plantation forestry: Development, benefits, ecosystem services and perspectives for the future, Global Ecology and Conservation, 15, e00419. 10.1016/j.gecco.2018.e00419 2018.

Long, Q., Zhan, Z., Du, H., Peng, W., Su, L., Zhang, H., Zeng, Z., Zeng, F., Tan, W., Mo, Y., Deng, X., Xie, Y., and Wang, K.: Environmental filtering and dispersal limitation jointly shape the taxonomic, functional and phylogenetic diversity in a subtropical karst forest of China, Frontiers in Plant Science, 16, 1655071, 10.3389/fpls.2025.1655071 2025.

Lüttke, U.: Ecophysiology of Crassulacean Acid Metabolism (CAM), Annals of Botany, 93, 629–652, 10.1093/aob/mch087, 2004.

Lyu, X., Li, P., Jin, L., Yang, F., Pucker, B., Wang, C., Liu, L., Zhao, M., Shi, L., Zhang, Y., Yang, Q., Xu, K., Li, X., Hu., Z., Yang, J., Yu, J., and Zhang, M.: 2024. Tracing the evolutionary and genetic footprints of atmospheric tillandsioids transition from land to air, Nature Communication 15, 9599, 10.1038/s41467-024-53756-7 2024.

Marani, M., Da Lio, C., and D’Alpaos, A.: Vegetation engineers marsh morphology through multiple competing stable states, Proceedings of the National Academy of Sciences of the United States of America, 110(9), 3259–3263, 10.1073/pnas.1218327110 2013.

Mastretta-Yanes, A., Arrigo, N., Alvarez, N., Jorgensen, T.H., Piñero, D., and Emerson, B.C.: Restriction site-associated DNA sequencing, genotyping error estimation and de novo assembly optimization for population genetic inference, Molecular Ecology Resources, 15, 28–41, 10.1111/1755-0998.12291, 2015.

McKee, E.D.: 1979. A study of global sand seas, U.S. Government Printing Office, Washington, DC., UCA, 429 pp., 10.3133/pp1052, 1979.

Mendieta-Leiva, G., Buckley, H.L., Zotz, G.: Directional changes over time in the species composition of tropical vascular epiphyte assemblages, Journal of Ecology, 110, 553–568, 10.1111/1365-2745.13817, 2022.

Meron, E.: Vegetation pattern formation: The mechanisms behind the forms, Physics Today, 72, 30–36, 10.1063/PT.3.4340, 2019.

Merklinger, F.F., Zheng, Y., Luebert, F., Harpke, D., Böhnert, T., Stoll, A., Koch, M.A., Blattner, F.R., Wiehe, T., Quandt, D.: Population genomics of *Tillandsia landbeckii* reveals unbalanced genetic diversity and founder effects in the Atacama Desert, Global and Planetary Change, 184, 103076, 10.1016/j.gloplacha.2019.103076 2020.

Mikulane, S., Siegmund, A., Del Rio, C., Koch, M.A., Osses, P.: Remote sensing-based mapping of *Tillandsia* fields: A semi-automated detection approach in the hyperarid coastal Atacama Desert, northern Chile, Journal of Arid Environments, 205, 104821, 10.1016/j.jaridenv.2022.104821 2022.

Möbus, J., Kiefer, C., Barfuss, M., Quandt, D., Koch, M.A.: Setting the evolutionary timeline: Tillandsia landbeckii in the Chilean Atacama Desert, Plant Systematics and Evolution, 307, e39, 10.1007/s00606-021-01760-5, 2021.

Morris, W.F., Ehrlén, J., Dahlgren, J.P., Loomis, A.K., and Louthan, A.M.: Biotic and anthropogenic forces rival climatic and abiotic factors in determining global plant population growth and fitness, Proceedings of the National Academy of Sciences of the United States of America, 117(2), 1107–1112, 10.1073/pnas.1918363117 2020.

Moura, M.R., Villalobos, F., Costa, G.C. and Garcia, P.C.A.: Disentangling the role of climate, topography and vegetation in species richness gradients, PLOS ONE, 11(3), e0152468, 10.1371/journal.pone.0152468 2016.

Nathan, J., Osem, Y., Shachak, M. and Meron, E.:. Linking functional diversity to resource availability and disturbance: A mechanistic approach for water-limited plant communities, Journal of Ecology, 104, 419–429, 10.1111/1365-2745.12525, 2016.

Nei, M.: Molecular evolutionary genetics, Columbia University Press, New York, 514 pp., 10.7312/nei-92038, 1987.

Nuñez, M.A. and Paritsis, J.: How are monospecific stands of invasive trees formed? Spatio-temporal evidence from Douglas fir invasions, AoB Plants, 10(4), ply041, 10.1093/aobpla/ply041, 2018.

Osses, P., Cereceda, P., Larrain, H. and Astaburuagam, J.P.: Distribution and geographical factors of the Tillandsia fields in the coastal mountain range of the Tarapacá Region, Chile, in: Proceedings of Fourth International Conference on Fog, Fog Collection and Dew, edited by: Biggs, A., and Cereceda, C., La Serena, Chile, 93–96, 2007.

Paschalis, A., Katul, G.G., Fatichi, S., Manoli, G., and Molnar, P.: Matching ecohydrological processes and scales of banded vegetation patterns in semiarid catchments, Water Resources Research, 52, 2259–2278, 10.1002/2015WR017679 2016.

Pichler, H.: Dynamik der Atmosphäre, Bibliographisches Institut, Mannheim, Germany, 459 pp., ISBN 9783411031412, 1986.

Posit team: RStudio: Integrated development environment for R, Posit Software, PBC [code], http://www.posit.co/, 2025.

Purcell, S., Neale, B., Todd-Brown, K., Thomas, L., Ferreira, M.A., Bender, D., Maller, J., Sklar, P., de Bakker, P.I., Daly, M.J., and Sham, P.C.: PLINK: A tool set for whole-genome association and population-based linkage analyses, American Journal of Human Genetics, 81(3), 559–575. 10.1086/519795. 2007.

Puritz, J.B., Hollenbeck, C.M., and Gold, J.R.: dDocent: A RADseq, variant-calling pipeline designed for population genomics of non-model organisms, PeerJ, 2, e431. 10.7717/peerj.431. 2014.

QGIS.org: QGIS geographic information system, QGIS Association, QGIS [software], http://www.qgis.org, 2025.

Qian, H., Zhang, J., Jin, Y., and Deng, T.: Effects of evolutionary history on assembly of flowering plants in regions across Africa, Ecography, 2023, e06775, 10.1111/ecog.06775 2023.

R Core Team: R: A language and environment for statistical computing, R Foundation for Statistical Computing, Vienna, Austria, R Core Team [code], https://www.R-project.org/, 2021.

Raux, P.S., Gravelle, S., and Dumais, J.: Design of a unidirectional water valve in Tillandsia, Nature Communications, 11, 396. 10.1038/s41467-019-14236-5. 2020.

Rein, B., Lückge, A., Sirocko, F.: A major Holocene ENSO anomaly during the Medieval period, Geophys. Res. Lett., 31, L17211, 10.1029/2004GL020161 2004.

Rennemüller, J.: Sedimentological analysis of two stratified push cores at Tillandsia lomas sites of the northern and southern Chilean Atacama, Bachelor thesis, Faculty of Chemistry and Earth Sciences, Heidelberg University, Heidelberg, Germany, 53 pp., 2023.

Rietkerk, M., Dekker, S.C., de Ruiter, P.C., and van de Koppel, J.: Self-organized patchiness and catastrophic shifts in ecosystems, Science, 305, 1926–1929, 10.1126/science.1101867 2004.

Rundel, P.W., Palma, B., Dillon, M.O., Sharifi, M.R., Nilsen, E.T., and Boonpragob, K.: Tillandsia landbeckii in the coastal Atacama Desert of northern Chile, Revista Chilena de Historia Natural, 70, 341–349, https://rchn.biologiachile.cl/pdfs/1997/3/Rundel_et_al_1997.pdf, 1997.

Rutllant, J. A., Fuenzalida, H. and Aceituno, P.: Climate dynamics along the arid northern coast of Chile: The 1997–1998 Dinámica del Clima de la Región de Antofagasta (DICLIMA) experiment, J. Geophys. Res., 108(D17), 4538, 10.1029/2002JD003357 2003.

Schemenauer, R.S. and Cerceda, P.: A proposed standard fog collector for use in high elevation regions, Journal of Applied Meteorology, 33, 1313–1322, 1994.

Schulz, N., Aceituno, P., and Richter, M.: Phytogeographic divisions, climate change and plant dieback along the coastal desert of northern Chile, Erdkunde 65, 169–187, https://www.jstor.org/stable/23030664. 2011.

Schween, J.H., Hoffmeister, D. and Löhnert, U.: Filling the observational gap in the Atacama Desert with a new network of climate stations, Global and Planetary Change, 184, 103034. 10.1016/j.gloplacha.2019.103034. 2020.

Schween, J.: Average diurnal means from climate stations 2017–2024, CRC1211 Database [data set], https://www.crc1211db.uni-koeln.de/search/view.php?dataID=1040, 2025.

SERNAGEOMIN: Mapa Geológico de Chile: versión digital, Servicio Nacional de Geología y Minería, Publicación Geológica Digital, No. 4 (CD-ROM, versión 1.0, 2003), Santiago, Chile, 2003.

Smith, L.B., Downs, R.J.: Tillandsioideae (Bromeliaceae), Flora Neotropica, 14, 888, 1997.

Stein, E., Jäger, D., Quandt, D., and Koch, M.A.: High resolution digital terrain models (DTMs) of Tillandsiales in the Chilean Atacama Desert: The Arica study field, CRC1211 Database (CRC1211DB) [data set], 10.5880/CRC1211DB.70, 2024a.

Stein, E., Jäger, D., Quandt, D., and Koch, M.A.: High resolution digital terrain models (DTMs) of Tillandsiales in the Chilean Atacama Desert: The Caldera study field, CRC1211 Database (CRC1211DB) [data set], 10.5880/CRC1211DB.69, 2024b.

Stein, R.E., Luque-Fernández, C.R., Kiefer, C., Möbus, J., Pauca-Tanco, G.A., Jabbusch, S., Harpke, D., Bechteler, J., Quandt, D., Villasante, F., and Koch, M.A.: Climate-driven past and present interspecies gene flow may have contributed to shape microscale adaptation capacity in *Tillandsia* lomas in hyperarid South American desert systems, Global and Planetary Change, 230, 104258, 10.1016/j.gloplacha.2023.104258 2023.

Takahashi, K., and Battisti, D. S.: Processes controlling the mean tropical Pacific precipitation pattern. Part I: The Andes and the eastern Pacific ITCZ, Journal of Climate, 20, 3434–3451, 10.1175/JCLI4198.1 2007.

Tel-zur, N., Abbo, S., Myslabodski, D., and Tel-Zur, N.: Modified CTAB procedure for DNA isolation from epiphytic cacti of the genera *Hylocereus* and *Selenicereus* (Cactaceae), Plant Molecular Biology Reporter, 17, 249–254. 10.1023/A:1007656315275, 1999.

Till, W.: Die Untergattung Diaphoranthema von Tillandsia, 4. Teil: Das Tillandsia recurvata Aggregat, Die Bromelie, 1, 15–20, 1992.

Urban, M.C., Bocedi, G., Hendry, A.P., Mihoub, J.B., Peer, G., Singer, A., Bridle, J.R., Crozier, L.G., De Mester, L., Godsoe, W., Gonzalez, A., Hellmann, J.J., Holt, R.D., Huth, A., Jojhst, K., Krug, C.B., Leadley, P.W., Palmer, S.C.F., Pantel, J.H., Schmitz, A., Zollner, P.A., and Travis, J.M.J.: Improving the forecast for biodiversity under climate change, Science, 353, aad8466, 10.1126/science.aad8466, 2016.

Voigt, C., Klipsch, S., Herwartz, D., Chong, G., and Staubwasser, M.: The spatial distribution of soluble salts in the surface soil of the Atacama Desert and their relationship to hyperaridity, Global and Planetary Change, 184, 103077. 10.1016/j.gloplacha.2019.103077 2020.

Wang, L., Du, X., Liu, J., Zhang, J., Lv, S.: Effects of grazing on plant functional groups across spatial scales in *Stipa breviflora* desert steppe, Frontiers in Plant Science, 16, 1643655, 10.3389/fpls.2025.1643655 2025.

Wang, T.-Y., Wang, P., Wang, Z.-L., Niu, G.-Y., Yu, J.-J., Ma, N., Wu, Z.-N., Pozdniakov, S.P., and Yan, D.-H.: Drought adaptability of phreatophytes: Insight from vertical root distribution in drylands of China, Journal of Plant Ecology, 14(6), 1128–1142, 10.1093/jpe/rtab059 2021.

Wang, X., and Zhang, G.: The influence of infiltration feedback on the characteristic of banded vegetation pattern on hillsides of semiarid area. PLOS ONE, 14(1), e0205715. 10.1371/journal.pone.0205715 2019.

Wei, T., and Simko, V.: R package ‘corrplot’: Visualization of a correlation matrix (version 0.95), Githup [code], https://github.com/taiyun/corrplot, 2024.

Wendler, N., Mascher, M., Nöh, C., Himmelbach, A., Scholz, U., Ruge-Wehling, B., Stein, N.: Unlocking the secondary gene pool of barley with next-generation sequencing, Plant Biotechnology Journal, 12, 1122–1131.

10.1111/pbi.12219, 2014.

Wickham, H.: ggplot2: Elegant graphics for data analysis. Springer-Verlag, New York, US, 260 pp., 10.1007/978-3-319-24277-4, 2016.

Wickham, H., Averick, M., Bryan, J., Chang, W., McGowan, L.D., François, R., Grolemund, G., Hayes, A., Henry, L., Hester, J., Kuhn, M., Pedersen, T.L., Miller, E., Bache, S.M., Müller, K., Ooms, J., Robinson, D., Seidel, D.P., Spinu, V., and Yutani, H.: Welcome to the tidyverse, Journal of Open Source Software, 4(43), 1686. 10.21105/joss.01686 2019.

Wieringa, J.: Representative roughness parameters for homogeneous terrain, Boundary-Layer Meteorology, 63, 323–363, 10.1007/BF00705357, 1993.

Willi, Y., Lucek, K., Bachmann, O., and Walden, N.: Recent speciation associated with range expansion and a shift to self-fertilization in North American *Arabidopsis*, Nature Communications, 13, 7564, 10.1038/s41467-022-35368-1 2022.

Yamori, W., Hikosaka, K., and Way, D.A.: Temperature response of photosynthesis in C3, C4, and CAM plants: temperature acclimation and temperature adaptation, Photosynthesis Research 119, 101–117, 10.1007/s11120-013-9874-6, 2014.

Zhang, H., Fechete, L., Himmelbach, A., Poehlein, A., Lohwasser, U., Börner, A., Maalouf, F., Kumar, S., Khazaei, H., Stein, N., and Jayakodi, M.: Optimization of genotyping-by-sequencing (GBS) for germplasm fingerprinting and trait mapping in faba bean, Legume Science, 6, e254. 10.1002/leg3.254 2024.

