## Supplementary File 2 for "Drowning in a sandy ocean: Epiarenic growth of *Tillandsia* in the hyperarid Atacama Desert"

Code


### Code

#### 2026-02-05

- A) GBS
  Processing
  - dDocent pipeline (BASH)
    - 1) Trim raw
      reads using trimmomatic (adapter trimming)
    - 2) dDocent: denovo
      reference assembly (rainbow)
    - 3) dDocent: bwa-mem mapping
    - 4) dDocent: FreeBayes SNP
      calling
    - 5) SNP
      filtering
  - Filtering
    Summary
  - Quality
    Control (R)
    - Determine error rate
    - Compute
      site-level discordance (likely paralogs etc.)
    - use plink for ADMIXTURE
      input (BASH)
    - run
      Admixture
  - Global Analysis of VCF (R)
    - PCA results
    - Heatmap of genetic
      distances
    - ADMIXTURE
      results
    - Genetic Diversity Scores
    - Identify
      clones
  - Local
    analyses
- B)
  Sand Data Processing
  - Boxplot1 - Field -
    Grainsize
  - Boxplot2 - Plant
    Occurence - Grainsize
  - Boxplot3 - Field ID -
    Sorting Index
  - Boxplot4 - Plant
    Occurence- Sorting Index
  - Statistical Analysis
- C) Weather and Terrain
  Data Processing
  - Wind speeds and directions
  - Yearly
    summaries
    - by Station
    - by Region
  - Plots of mean of daily
    mean/min/max
  - Drift
    Potential
    - Drift Potential Sandrose
    - Net
      Sand Drift vector
  - Mean exposition vector
  - Correlation Matrix and
    Table

### A) GBS Processing

```
# Load libraries
library(tidyverse)
library(ggplot2)
library(dplyr)
library(tidyr)
library(readr)
library(wesanderson)
library(ggsci)
library(viridis)
```

#### dDocent pipeline (BASH)

##### 1) Trim raw reads using trimmomatic (adapter trimming)

```
#!/bin/bash

# Define paths
INPUT_DIR="/rawfiles"
OUTPUT_DIR="/trimmed"
MAP_FILE="trimming_map.tsv"
ADAPTERS="TruSeq3-SE.fa"  # Update this path
LOG_DIR="${OUTPUT_DIR}/logs"
STATS_FILE="${OUTPUT_DIR}/trimming_summary.tsv"

# Create necessary directories
mkdir -p "$OUTPUT_DIR" "$LOG_DIR"

# Initialize stats file
echo -e "rawfile\ttrimmedfile\tinput_reads\toutput_reads\tdate" > "$STATS_FILE"

# Process each sample
tail -n +2 "$MAP_FILE" | while IFS=$'\t' read -r rawfile trimmedfile; do
    INFILE="$INPUT_DIR/$rawfile"
    OUTFILE="$OUTPUT_DIR/$trimmedfile"
    LOGFILE="$LOG_DIR/${trimmedfile%.fq.gz}.log"

    echo "Trimming $rawfile → $trimmedfile"

    # Run Trimmomatic with logging
    Trimmomatic SE -phred33 -threads 20 \
        "$INFILE" "$OUTFILE" \
        ILLUMINACLIP:"$ADAPTERS":2:30:10 \
        LEADING:20 TRAILING:20 SLIDINGWINDOW:4:25 MINLEN:50 \
        &> "$LOGFILE"

    # Extract stats
    input_reads=$(zcat "$INFILE" | echo $((`wc -l`/4)))
    output_reads=$(zcat "$OUTFILE" | echo $((`wc -l`/4)))
    timestamp=$(date +"%Y-%m-%d %H:%M:%S")

    # Append to summary
    echo -e "$rawfile\t$trimmedfile\t$input_reads\t$output_reads\t$timestamp" >> "$STATS_FILE"
done
```

##### 2) dDocent: denovo reference assembly (rainbow)

- 20 individuals (10 Caldera, 10 Iquique)
- Filter for reads starting with Pst1 restriction site
- cluster similarity >0.99, depth >10 in >6
  individuals

##### 3) dDocent: bwa-mem mapping

- Parameters -A 2 -B 5 -O 12

##### 4) dDocent: FreeBayes SNP calling

##### 5) SNP filtering

```
#!/bin/bash

echo "Welcome to the ddRAD filtering pipeline with checkpoints and site tallies."

# Logging setup
log_file="pipeline.log"
exec > >(tee -i "$log_file")
exec 2>&1
echo "Logging enabled. All outputs will be saved to $log_file."

# Input file selection
read -p "Enter the path to your initial VCF file: " initial_vcf

# Customizable parameters
read -p "Enter max missing threshold (default: 0.95): " max_missing
max_missing=${max_missing:-0.95}

read -p "Enter min allele frequency threshold (MAF, default: 0.01): " maf
maf=${maf:-0.01}

read -p "Enter min mean depth threshold (default: 5): " min_mean_dp
min_mean_dp=${min_mean_dp:-5}

read -p "Enter max mean depth threshold (default: 120): " max_mean_dp
max_mean_dp=${max_mean_dp:-120}

##############################################
# Step 1: Remove indels
##############################################
if [ ! -f 1_filtered_no_indels.recode.vcf ]; then
    echo "Step 1: Removing indels"
    vcftools --vcf "$initial_vcf" --remove-indels --recode --recode-INFO-all --out 1_filtered_no_indels
fi
echo "After Step 1: $(grep -vc '^#' 1_filtered_no_indels.recode.vcf) sites"

##############################################
# Step 2: Pre-filter by missingness (0.5), MAC, Q30
##############################################
if [ ! -f 2_filtered_missing_mac_quality.recode.vcf ]; then
    echo "Step 2: Pre-filtering by max missing 0.5, MAC 3, Q30"
    vcftools --vcf 1_filtered_no_indels.recode.vcf --max-missing 0.5 --mac 3 --minQ 30 --recode --recode-INFO-all --out 2_filtered_missing_mac_quality
fi
echo "After Step 2: $(grep -vc '^#' 2_filtered_missing_mac_quality.recode.vcf) sites"

##############################################
# Step 3: Minimum depth filter
##############################################
if [ ! -f 3_filtered_min_depth.recode.vcf ]; then
    echo "Step 3: Filtering by minimum depth (DP3)"
    vcftools --vcf 2_filtered_missing_mac_quality.recode.vcf --minDP 3 --recode --recode-INFO-all --out 3_filtered_min_depth
fi
echo "After Step 3: $(grep -vc '^#' 3_filtered_min_depth.recode.vcf) sites"

##############################################
# Step 4: Missing data per individual
##############################################
if [ ! -f 6_filtered_removed_low_dp_individuals.recode.vcf ]; then
    echo "Step 4: Assessing missing data per individual"
    vcftools --vcf 3_filtered_min_depth.recode.vcf --missing-indv
    mawk '!/IN/' out.imiss | cut -f5 > 4_missing_data_percentages.txt

    gnuplot << \EOF
set terminal dumb size 120, 30
set autoscale
unset label
set title "Histogram of % missing data per individual"
set ylabel "Number of Occurrences"
set xlabel "% of missing data"
binwidth=0.01
bin(x,width)=width*floor(x/width) + binwidth/2.0
plot '4_missing_data_percentages.txt' using (bin($1,binwidth)):(1.0) smooth freq with boxes
pause -1
EOF

    read -p "Enter the cutoff for % missing data per individual (e.g., 0.6): " missing_cutoff
    awk -v cutoff=$missing_cutoff '$5 > cutoff' out.imiss | cut -f1 > 5_individuals_to_remove.txt
    echo "The following individuals will be removed:"
    cat 5_individuals_to_remove.txt

    vcftools --vcf 3_filtered_min_depth.recode.vcf --remove 5_individuals_to_remove.txt --recode --recode-INFO-all --out 6_filtered_removed_low_dp_individuals
fi
echo "After Step 4: $(grep -vc '^#' 6_filtered_removed_low_dp_individuals.recode.vcf) sites"

##############################################
# Step 5: Apply stringent missingness, MAF, mean depth
##############################################
if [ ! -f 7_filtered_final_maf_depth.recode.vcf ]; then
    echo "Step 5: Applying max missing ($max_missing), MAF ($maf), min mean depth ($min_mean_dp)"
    vcftools --vcf 6_filtered_removed_low_dp_individuals.recode.vcf --max-missing $max_missing --maf $maf --min-meanDP $min_mean_dp --recode --recode-INFO-all --out 7_filtered_final_maf_depth
fi
echo "After Step 5: $(grep -vc '^#' 7_filtered_final_maf_depth.recode.vcf) sites"

##############################################
# Step 6: Site-level quality filters 
##############################################
if [ ! -f 11_filtered_quality_depth.vcf ]; then
    echo "Step 6: Applying AB, MQ ratio, and QUAL filters"

    vcffilter -s -f "AB > 0.25 & AB < 0.75 | AB < 0.01" 7_filtered_final_maf_depth.recode.vcf > 8_filtered_allele_balance.vcf
    echo "After AB filter: $(grep -vc '^#' 8_filtered_allele_balance.vcf) sites"

    vcffilter -f "MQM / MQMR > 0.85 & MQM / MQMR < 1.10" 8_filtered_allele_balance.vcf > 9_filtered_mapping_quality.vcf
    echo "After MQ ratio filter: $(grep -vc '^#' 9_filtered_mapping_quality.vcf) sites"

    vcffilter -f "QUAL > 30" 9_filtered_mapping_quality.vcf > 10_filtered_qual.vcf
    echo "After QUAL filter: $(grep -vc '^#' 10_filtered_qual.vcf) sites"

    cp 10_filtered_qual.vcf 11_filtered_quality_depth.vcf
fi
echo "After Step 6: $(grep -vc '^#' 11_filtered_quality_depth.vcf) sites"

##############################################
# Step 7: Depth distribution and filtering
##############################################
if [ ! -f 12_filtered_final_depth.recode.vcf ]; then
    echo "Step 7: Calculating mean depth and filtering sites with extreme depth..."
    vcftools --vcf 11_filtered_quality_depth.vcf --site-depth --out 11_filtered_quality_depth
    cut -f3 11_filtered_quality_depth.ldepth > 11_filtered_quality_depth.site.depth

    num_samples=$(grep -m 1 "^#CHROM" 11_filtered_quality_depth.vcf | awk '{print NF-9}')
    mawk '!/D/' 11_filtered_quality_depth.site.depth | mawk -v x=$num_samples '{print $1/x}' > meandepthpersite

    gnuplot << \EOF
set terminal dumb size 120, 30
set autoscale
set xrange [10:300]
unset label
set title "Histogram of mean depth per site"
set ylabel "Number of Occurrences"
set xlabel "Mean Depth"
binwidth=1
bin(x,width)=width*floor(x/width) + binwidth/2.0
set xtics 5
plot 'meandepthpersite' using (bin($1,binwidth)):(1.0) smooth freq with boxes
pause -1
EOF

    read -p "Enter the cutoff for mean depth per site (default: $max_mean_dp): " mean_depth_cutoff
    mean_depth_cutoff=${mean_depth_cutoff:-$max_mean_dp}

# overwrite the global so summary uses the chosen value
max_mean_dp=$mean_depth_cutoff

    vcftools --vcf 11_filtered_quality_depth.vcf --max-meanDP $mean_depth_cutoff --recode --recode-INFO-all --out 12_filtered_final_depth
fi
echo "After Step 7: $(grep -vc '^#' 12_filtered_final_depth.recode.vcf) sites"

##############################################
# Final output
##############################################
if [ ! -f FinalFiltered.vcf ]; then
    read -p "Enter the desired name for the final filtered VCF file (e.g., FinalFiltered.vcf): " final_vcf
    mv 12_filtered_final_depth.recode.vcf "$final_vcf"
    echo "Filtering complete. Final VCF file: $final_vcf"
else
    echo "Final VCF already exists, skipping."
fi

##############################################
# Summary report
##############################################
echo ""
echo "========== FILTERING SUMMARY =========="

# Initial counts
init_sites=$(grep -vc '^#' "$initial_vcf")
init_indiv=$(grep -m1 "^#CHROM" "$initial_vcf" | awk '{print NF-9}')
echo "Initial dataset:            $init_sites sites, $init_indiv individuals"

# Step 1
sites1=$(grep -vc '^#' 1_filtered_no_indels.recode.vcf)
echo "Step 1 (No indels):         $sites1 sites ($(awk -v a=$sites1 -v b=$init_sites 'BEGIN{printf "%.1f", (a/b*100)}')% of initial)"

# Step 2
sites2=$(grep -vc '^#' 2_filtered_missing_mac_quality.recode.vcf)
echo "Step 2 (Pre-filter):        $sites2 sites ($(awk -v a=$sites2 -v b=$init_sites 'BEGIN{printf "%.1f", (a/b*100)}')%)"

# Step 3
sites3=$(grep -vc '^#' 3_filtered_min_depth.recode.vcf)
echo "Step 3 (Min depth):         $sites3 sites ($(awk -v a=$sites3 -v b=$init_sites 'BEGIN{printf "%.1f", (a/b*100)}')%)"

# Step 4
sites4=$(grep -vc '^#' 6_filtered_removed_low_dp_individuals.recode.vcf)
indiv4=$(grep -m1 "^#CHROM" 6_filtered_removed_low_dp_individuals.recode.vcf | awk '{print NF-9}')
echo "Step 4 (Indiv removal):     $sites4 sites ($(awk -v a=$sites4 -v b=$init_sites 'BEGIN{printf "%.1f", (a/b*100)}')%), $indiv4 individuals ($(awk -v a=$indiv4 -v b=$init_indiv 'BEGIN{printf "%.1f", (a/b*100)}')% of initial)"

# Step 5
sites5=$(grep -vc '^#' 7_filtered_final_maf_depth.recode.vcf)
echo "Step 5 (Max-miss/MAF/DP):   $sites5 sites ($(awk -v a=$sites5 -v b=$init_sites 'BEGIN{printf "%.1f", (a/b*100)}')%)"

# Step 6
sites6=$(grep -vc '^#' 11_filtered_quality_depth.vcf)
echo "Step 6 (Quality filters):   $sites6 sites ($(awk -v a=$sites6 -v b=$init_sites 'BEGIN{printf "%.1f", (a/b*100)}')%)"

# Step 7
if [ -f "$final_vcf" ]; then
    sites7=$(grep -vc '^#' "$final_vcf")
    indiv7=$(grep -m1 "^#CHROM" "$final_vcf" | awk '{print NF-9}')
    echo "Step 7 (Depth cutoff):      $sites7 sites ($(awk -v a=$sites7 -v b=$init_sites 'BEGIN{printf "%.1f", (a/b*100)}')%), $indiv7 individuals"
fi

# Final
if [ -f "$final_vcf" ]; then
    final_sites=$(grep -vc '^#' "$final_vcf")
    final_indiv=$(grep -m1 "^#CHROM" "$final_vcf" | awk '{print NF-9}')
    echo "Final filtered dataset:     $final_sites sites ($(awk -v a=$final_sites -v b=$init_sites 'BEGIN{printf "%.1f", (a/b*100)}')%), $final_indiv individuals ($(awk -v a=$final_indiv -v b=$init_indiv 'BEGIN{printf "%.1f", (a/b*100)}')% of initial)"
fi

echo ""
echo "========== PARAMETERS USED =========="
echo "Missing cutoff (Step 4):    $missing_cutoof"
echo "Max missing (Step 5):       $max_missing"
echo "MAF (Step 5):               $maf"
echo "Min mean depth (Step 5):    $min_mean_dp"
echo "Max mean depth (Step 7):    $max_mean_dp"
echo "Allele balance (Step 6):    AB > 0.25 & AB < 0.75 | AB < 0.01"
echo "MQM/MQMR ratio (Step 6):    0.85 – 1.10"
echo "QUAL filter (Step 6):       QUAL > 30"
echo "======================================="
```

- refilter any individuals >0.3 missingness

```
vcftools --vcf FINAL_allipk_pst1ref.vcf --missing-indv
awk '$5 > 0.3 {print $1}' out.imiss > remove_indv.txt
vcftools --vcf FINAL_allipk_pst1ref.vcf --remove remove_indv.txt --recode --out FINAL_allipk_pst1ref_postfilter
```

#### Filtering Summary

- Initial dataset: 778893 sites, 979 individuals
- Step 1 (No indels): 667605 sites (85.7% of initial)
- Step 2 (Pre-filter): 82482 sites (10.6%)
- Step 3 (Min depth): 82482 sites (10.6%)
- Step 4 (Indiv removal): 82482 sites (10.6%), 872 individuals
  (89.1% of initial)
- Step 5 (Max-miss/MAF/DP): 4920 sites (0.6%)
- Step 6 (Quality filters): 2754 sites (0.4%)
- Final filtered dataset: 2726 sites (0.3%), 872 individuals (89.1%
  of initial)

========== PARAMETERS USED ==========

- Missing cutoff (Step 4): 0.66
- Max missing (Step 5): 0.8
- MAF (Step 5): 0.01
- Min mean depth (Step 5): 5
- Max mean depth (Step 7): 120
- Allele balance (Step 6): AB > 0.25 & AB < 0.75 | AB
  < 0.01
- MQM/MQMR ratio (Step 6): 0.85 – 1.10
- QUAL filter (Step 6): QUAL > 30

#### Quality Control (R)

```
library(vcfR)
library(dplyr)

# Read your filtered VCF
vcf <- read.vcfR("FINAL_allipk_pst1ref_postfilter.recode.vcf")
```

```
## Scanning file to determine attributes.
## File attributes:
##   meta lines: 63
##   header_line: 64
##   variant count: 2726
##   column count: 727
## Meta line 63 read in.
## All meta lines processed.
## gt matrix initialized.
## Character matrix gt created.
##   Character matrix gt rows: 2726
##   Character matrix gt cols: 727
##   skip: 0
##   nrows: 2726
##   row_num: 0
## Processed variant 1000Processed variant 2000Processed variant: 2726
## All variants processed
```

```
# read in metadata
meta <- read.csv("metadata.csv", sep = ";")

# Convert to genlight (adegenet) or a genotype matrix
library(adegenet)
gl <- vcfR2genlight(vcf)
geno_matrix <- as.matrix(gl)
rownames(geno_matrix) <- indNames(gl)
```

##### Determine error rate

```
dup_labels <- c("rep")

# --- 2. Build replicate pairs robustly ---
replicate_pairs_df <- meta %>%
  filter(sample %in% indNames(gl)) %>%
  group_by(plantID, category) %>%
  # keep only groups with at least one duplicate AND one original
  filter(any(rep %in% dup_labels) & any(rep == "")) %>%
  # extract original sample safely
  mutate(original = sample[rep == ""][1]) %>%   # take first if multiple originals
  # keep only duplicates
  filter(rep %in% dup_labels) %>%
  select(original, duplicate = sample, dup_category = category) %>%
  ungroup()

# --- 3. Extract genotype matrix ---
geno_matrix <- as.matrix(gl)
rownames(geno_matrix) <- indNames(gl)

# --- 4. Compute error rates per replicate pair ---
error_rates <- apply(replicate_pairs_df, 1, function(row) {
  orig <- row["original"]
  dup  <- row["duplicate"]
  
  if (!(orig %in% rownames(geno_matrix)) || !(dup %in% rownames(geno_matrix))) {
    return(NA_real_)
  }
  
  g1 <- geno_matrix[orig, ]
  g2 <- geno_matrix[dup, ]
  
  valid_idx <- !(is.na(g1) | is.na(g2))
  total_compared <- sum(valid_idx)
  
  if (total_compared == 0) return(NA_real_)
  
  mismatches <- sum(g1[valid_idx] != g2[valid_idx])
  mismatches / total_compared
})

# --- 5. Create dataframe with duplicate info ---
error_df <- replicate_pairs_df %>%
  mutate(error_rate = error_rates)

print(error_df)
```

```
## # A tibble: 72 × 5
##    plantID  original                       duplicate     dup_category error_rate
##    <chr>    <chr>                          <chr>         <chr>             <dbl>
##  1 C-U03-a  CALDERA_3093924-C-U03.trimmed  CALDERA_3093… fieldsample      0.134 
##  2 A-AF07-a ARICA_3093922-A-AF07.trimmed   ARICA_320531… fieldsample      0.141 
##  3 C-AD08-a CALDERA_3243963-C-AD08.trimmed CALDERA_3243… fieldsample      0.482 
##  4 C-Q18-a  CALDERA_3074268-C-Q18.trimmed  CALDERA_3244… fieldsample      0.179 
##  5 C-A15-a  CALDERA_3074197-C-A15.trimmed  CALDERA_3285… fieldsample      0.159 
##  6 C-A16-a  CALDERA_3074198-C-A16.trimmed  CALDERA_3285… fieldsample      0.189 
##  7 C-C15-a  CALDERA_3074203-C-C15.trimmed  CALDERA_3285… fieldsample      0.0965
##  8 C-C17-a  CALDERA_3074204-C-C17.trimmed  CALDERA_3285… fieldsample      0.125 
##  9 C-D18-a  CALDERA_3074206-C-D18.trimmed  CALDERA_3285… fieldsample      0.0998
## 10 C-D22-a  CALDERA_3074209-C-D22.trimmed  CALDERA_3285… fieldsample      0.0945
## # ℹ 62 more rows
```

```
# --- 6. Visualize distribution ---
ggplot(error_df, aes(x = error_rate)) +
  geom_histogram(binwidth = 0.02, fill = "steelblue", color = "white") +
  theme_minimal() +
  labs(title = "Distribution of replicate error rates",
       x = "Error rate",
       y = "Number of samples")
```

```
# --- 7. Mean error rate per category ---
error_df %>%
  group_by(dup_category) %>%
  summarise(mean_error_rate = mean(error_rate, na.rm = TRUE))
```

##### Compute site-level discordance (likely paralogs etc.)

```
# Build all pairs index
pairs <- error_df %>% select(original, duplicate)

# replicate_pairs_df: original, duplicate columns
geno_matrix <- as.matrix(gl)
rownames(geno_matrix) <- indNames(gl)

# Compute mismatch rate per SNP across all replicate pairs
site_discord <- apply(geno_matrix, 2, function(snp) {
  mism <- 0; comp <- 0
  for (i in seq_len(nrow(pairs))) {
    g1 <- snp[pairs$original[i]]
    g2 <- snp[pairs$duplicate[i]]
    if (!is.na(g1) && !is.na(g2)) {
      comp <- comp + 1
      mism <- mism + as.integer(g1 != g2)
    }
  }
  if (comp == 0) return(NA_real_)
  mism / comp
})

plot(site_discord)
```

##### use plink for ADMIXTURE input (BASH)

```
# 1) Convert VCF to PLINK binary, treat half-calls as missing
plink --vcf FINAL_allipk_pst1ref_postfilter.recode.vcf \
  --double-id \
  --allow-extra-chr \
  --snps-only just-acgt \
  --vcf-half-call m \
  --make-bed --out admix_00_raw

# 2) Assign unique IDs to variants (contig:position)
plink --bfile admix_00_raw \
  --allow-extra-chr \
  --set-missing-var-ids @:# \
  --make-bed --out admix_01_withIDs

# 3) allow up to 20% missing per SNP
plink --bfile admix_01_withIDs \
  --allow-extra-chr \
  --geno 0.20 \
  --make-bed --out admix_02_filtered


# 4) LD pruning (window 50 SNPs, step 5, r^2 > 0.2)
plink --bfile admix_02_filtered \
  --allow-extra-chr \
  --indep-pairwise 50 5 0.2 \
  --out admix_ld

# 5) Extract pruned SNPs
plink --bfile admix_02_filtered \
  --allow-extra-chr \
  --extract admix_ld.prune.in \
  --make-bed --out admix_03_pruned

# 6) Generate PCA file  
plink --bfile admix_03_pruned \
  --allow-extra-chr \
  --recode A \
  --out pca_input
```

> Total genotyping rate is 0.897029. 1127 variants and 718 people pass
> filters and QC.

##### run Admixture

```
for K in 1 2 3 4 5 6 7 8 9 10 11 12 13 14 15; do
  admixture --cv=10 admix_03_pruned.bed $K | tee log${K}.out
done
```

#### Global Analysis of VCF (R)

```
library(vcfR)
library(dplyr)

# Read your filtered VCF
vcf <- read.vcfR("FINAL_allipk_pst1ref_postfilter.recode.vcf")
```

```
## Scanning file to determine attributes.
## File attributes:
##   meta lines: 63
##   header_line: 64
##   variant count: 2726
##   column count: 727
## Meta line 63 read in.
## All meta lines processed.
## gt matrix initialized.
## Character matrix gt created.
##   Character matrix gt rows: 2726
##   Character matrix gt cols: 727
##   skip: 0
##   nrows: 2726
##   row_num: 0
## Processed variant 1000Processed variant 2000Processed variant: 2726
## All variants processed
```

```
gl <- vcfR2genlight(vcf)
geno_matrix <- as.matrix(gl)
rownames(geno_matrix) <- indNames(gl)

# attach pop info
pop(gl) <- meta$population[match(indNames(gl), meta$sample)]
```

##### PCA results

```
library(tidyverse)

# Read genotype matrix
geno <- read_delim("pca_input.raw", delim = " ")

# Drop non-genotype columns
geno_clean <- geno %>%
  select(-FID, -PAT, -MAT, -SEX, -PHENOTYPE) %>%
  column_to_rownames("IID")

# Remove SNPs with too much missing data
geno_clean <- geno_clean %>% select(where(~ mean(is.na(.)) < 0.2))

# Neutrally impute missing values
geno_imputed <- geno_clean %>%
  mutate(across(everything(), ~ ifelse(is.na(.), mean(., na.rm = TRUE), .)))
```

```
pca <- prcomp(geno_imputed, scale. = TRUE)

summary(pca)$importance[2, 1:5]  # PC1–PC5
```

```
##     PC1     PC2     PC3     PC4     PC5 
## 0.25055 0.16773 0.05133 0.02965 0.02723
```

```
# merge with meta data
scores <- as_tibble(pca$x) %>%
  mutate(sample = rownames(pca$x)) %>%
  left_join(meta, by = "sample")
```

```
ggplot(scores, aes(x = PC1, y = PC2, color = population, shape = category)) +
  geom_point(size = 3, alpha = 0.8) +
  theme_minimal() +
  labs(title = "PCA of ddRAD SNP matrix", x = "PC1", y = "PC2")
```

###### Removal of outliers

Three samples are obviously misplaced within PCA and are likely a
result of label switching errors, they will be removed for downstream
analyses:

```
outliers <- c("CALDERA_3243989-C-AH09.trimmed","CALDERA_3243963-C-AD08.trimmed","CALDERA_3243983-C-AF11.trimmed")

# Remove outlier individuals from the genlight object
gl <- gl[!indNames(gl) %in% outliers, ]
```

##### Heatmap of genetic distances

```
library(pheatmap)
library(poppr)

# Compute SNP-based pairwise distances
d <- bitwise.dist(gl)   # returns a dist object
d_mat <- as.matrix(d)   # convert to matrix
```

```
annotation <- data.frame(population = pop(gl))
rownames(annotation) <- indNames(gl)

pheatmap(d_mat,
         annotation_row = annotation,
         annotation_col = annotation,
         show_colnames = F,
         show_rownames = F,
         clustering_distance_rows = "euclidean",
         clustering_distance_cols = "euclidean",
         clustering_method = "average",
         color = colorRampPalette(c("blue", "white", "red"))(100),
         main = "Genetic distances")
```

##### ADMIXTURE results

###### CV errors

```
library(tidyverse)

# 1) Read all log files into a tibble
logs <- list.files(pattern = "log[0-9]+\\.out")

cv_df <- map_dfr(logs, function(f) {
  # read lines
  lines <- readLines(f)
  # find the line with CV error
  cv_line <- grep("CV error", lines, value = TRUE)
  # extract K and CV value using regex
  k <- as.integer(str_match(cv_line, "CV error \\(K=(\\d+)\\)")[,2])
  cv <- as.numeric(str_match(cv_line, ": ([0-9\\.]+)")[,2])
  tibble(K = k, CV = cv)
})

# 2) Plot CV error vs K
ggplot(cv_df, aes(x = K, y = CV)) +
  geom_line(color = "steelblue") +
  geom_point(size = 2, color = "darkred") +
  geom_point(data = cv_df %>% filter(CV == min(CV)),
             aes(x = K, y = CV), color = "green", size = 3) +
  labs(title = "ADMIXTURE CV Error by K",
       x = "Number of clusters (K)",
       y = "Cross-validation error")
```

###### Plot all K’s

```
# Path to your ADMIXTURE outputs
prefix <- "admix_03_pruned"   # change to your dataset prefix
Kvals <- 2:15                       # range of K values you ran

# Read FAM file for sample IDs
fam <- read_table2(paste0(prefix, ".fam"), col_names = FALSE)
sample_ids <- fam$X2
```

```
read_Q <- function(K) {
  qfile <- paste0(prefix, ".", K, ".Q")
  Q <- read_table2(qfile, col_names = FALSE)
  colnames(Q) <- paste0("Cluster", 1:K)
  Qdf <- cbind(sample = sample_ids, Q)
  Qdf$K <- K
  Qdf
}

Qall <- map_df(Kvals, read_Q)

# Merge with metadata and sort by elevation
Qall <- left_join(Qall, meta, by = "sample", unmatched = "drop") %>% arrange(population, elevation)

# remove outliers
outliers <- c("CALDERA_3243989-C-AH09.trimmed","CALDERA_3243963-C-AD08.trimmed","CALDERA_3243983-C-AF11.trimmed")
Qall <- Qall %>% filter(!sample %in% outliers)

# Reshape for plotting
Qlong <- Qall %>%
  pivot_longer(starts_with("Cluster"),
               names_to = "Cluster",
               values_to = "Ancestry")

# Force Cluster levels to be ordered numerically
Qlong$Cluster <- factor(Qlong$Cluster,
                        levels = paste0("Cluster", 1:max(Kvals)))

# preserve order in ggplot
Qlong$sample <- factor(Qlong$sample, levels = unique(Qlong$sample))

# Generate 15 distinct colors from viridis
cluster_colors <- viridis(15, option = "D")

# Name them Cluster1, Cluster2, ...
names(cluster_colors) <- paste0("Cluster", 1:15)
```

```
ggplot(Qlong, aes(x = sample, y = Ancestry, fill = Cluster)) +
  geom_bar(stat = "identity", width = 1) +
  facet_grid(K ~ population, scales = "free_x", space = "free_x") +
  theme_bw() +
  theme(axis.text.x = element_blank(),
        axis.ticks.x = element_blank(),
        panel.spacing = unit(0.1, "lines")) +
  labs(x = "Individuals", y = "Ancestry proportion",
       title = "ADMIXTURE results across K values")+
  scale_fill_manual(values = cluster_colors)
```

###### Plot K = 7

```
# Filter to K=9 only
Q7 <- Qlong %>% filter(K == 7)

# Drop unused factor levels (important for the legend)
Q7$Cluster <- factor(Q7$Cluster, levels = paste0("Cluster", 1:7))

# Ensure sample order is preserved
Q7 <- Q7 %>%
  arrange(population, longitude) %>%
  mutate(sample = factor(sample, levels = unique(sample)))

# Plot stacked barplot for K=9
ggplot(Q7, aes(x = sample, y = Ancestry, fill = Cluster)) +
  geom_bar(stat = "identity", width = 1) +
  facet_wrap(~population, scales = "free_x", nrow = 1) +
  theme_bw() +
  theme(axis.text.x = element_blank(),
        axis.ticks.x = element_blank(),
        panel.spacing = unit(0.1, "lines")) +
  labs(x = "Individuals (ordered by elevation)",
       y = "Ancestry proportion",
       title = "ADMIXTURE results at K=7")
```

```
#  scale_fill_manual(values = cluster_colors)
```

```
Q7 %>% 
  filter(has_rep == TRUE) %>%
  ggplot(aes(x = sample, y = Ancestry, fill = Cluster)) +
  geom_bar(stat = "identity", width = 1) +
  geom_point(aes(x=sample, y = 0.1, shape = category, color = category), 
             size = 3, stroke = 0.5, inherit.aes = FALSE) +
  scale_shape_manual(values = c("fieldsample" = 17, "growthexp" = 16)) +
  scale_color_manual(values = c("fieldsample" = "black", "growthexp" = "green")) +
  facet_wrap(~ plantID, scales = "free") +
  theme_bw() +
  theme(axis.text.x = element_blank(),
        axis.ticks.x = element_blank(),
        panel.spacing = unit(0.1, "lines"),
        axis.text.y = element_blank()) +
  labs(x = "Individuals", y = "Ancestry proportion",
       title = "ADMIXTURE results (K=7) of duplicate samples with category markers",
       color = "Sample Category", shape = "Sample Category", fill = "Ancestry Cluster")
```

###### Maps K = 7

```
library(tidyverse)
library(scatterpie)
library(ggthemes)
```

```
# ensure lat/long are numeric
Qall <- Qall %>%
  mutate(
    latitude = as.numeric(latitude),
    longitude = as.numeric(longitude)
  )


# Select relevant columns
pie_data <- Qall %>%
  filter(K == 7) %>%
  select(sample, latitude, longitude, population, starts_with("Cluster"))
```

```
pal7 <- cluster_colors[paste0("Cluster", 1:7)]

for (pop in unique(pie_data$population)) {
  pie_subset <- pie_data %>% filter(population == pop)

  p <- ggplot() +
    geom_scatterpie(
      aes(x = longitude, y = latitude),
      data = pie_subset,
      cols = paste0("Cluster", 1:7),
      pie_scale = 0.8
    ) +
    coord_equal() +
    theme_minimal() +
    labs(title = paste("ADMIXTURE pie map:", pop),
         x = "Longitude", y = "Latitude")

  print(p)
}
```

```
# export for QGIS
write.csv(pie_data, "../QGIS/piechart_data.csv")
```

###### Q-matrix Distance Replicates

This codes extracts the Q-vectors for original and replicate pairs
and calculates several distance metrics to quantify divergence in
ancestry estimates between duplicate samples.

> L1 distance (Manhattan distance) - Measures the total absolute
> difference in ancestry proportions between two samples. It adds up how
> much each cluster’s value differs.
>
> Range: 0 to 2
>
> 0 means the Q‑vectors are identical Higher values mean larger overall
> differences

\[ L1 = \sum\_{k=1}^{K} \left|
q\_{k,\text{orig}} - q\_{k,\text{dup}} \right| \]

> L1\_scaled - A rescaled version of L1 distance that ranges from 0 to
> 1. It divides L1 by 2 so the values are easier to interpret.
>
> 0 = identical ancestry profiles
>
> 1 = completely different ancestry profiles
>
> This is the most intuitive “error rate” for ADMIXTURE outputs. It
> directly corresponds to the fraction of ancestry that would need to be
> redistributed to make the two Q-vectors identical.

\[ L1\_{\text{scaled}} = \frac{1}{2}
\sum\_{k=1}^{K} \left| q\_{k,\text{orig}} - q\_{k,\text{dup}} \right|
\]

> Cosine similarity - Measures how similar the shape or pattern of
> ancestry proportions is, regardless of magnitude.
>
> Range: 0 to 1
>
> 1 = identical pattern Values near 0 indicate completely different
> ancestry patterns
>
> Helpful when one cluster dominates the ancestry profile

\[ \text{Cosine} = \frac{ \sum\_{k=1}^{K}
q\_{k,\text{orig}} \cdot q\_{k,\text{dup}} }{ \sqrt{\sum\_{k=1}^{K}
q\_{k,\text{orig}}^2} \;\; \sqrt{\sum\_{k=1}^{K} q\_{k,\text{dup}}^2} }
\]

```
# Choose your K
K_use <- 7

Qwide <- Qall %>%
  filter(K == K_use) %>%
  select(sample, starts_with("Cluster")) %>%
  select(sample, paste0("Cluster", 1:K_use)) %>%
  distinct()

Qmat <- Qwide %>%
  column_to_rownames("sample") %>%
  as.matrix()


# Function to compute distances between two Q vectors
q_metrics <- function(q1, q2) {
  l1  <- sum(abs(q1 - q2))
  l1_scaled <- l1 / 2
  cos <- sum(q1 * q2) / (sqrt(sum(q1^2)) * sqrt(sum(q2^2)))
  tibble(L1 = l1, L1_scaled = l1_scaled, Cosine = cos)
}

replicate_pairs_df <- replicate_pairs_df %>% filter(!original %in% outliers & !duplicate %in% outliers)

q_error_df <- replicate_pairs_df %>%
  rowwise() %>%
  do({
    orig <- .$original
    dup  <- .$duplicate
    
    if (!(orig %in% rownames(Qmat)) | !(dup %in% rownames(Qmat))) {
      return(tibble(L1 = NA, L1_scaled = NA, Cosine = NA))
    }
    
    q1 <- Qmat[orig, ]
    q2 <- Qmat[dup, ]
    
    q_metrics(q1, q2)
  }) %>%
  bind_cols(replicate_pairs_df, .)


q_error_df
```

```
mean_L1 <- mean(q_error_df$L1_scaled, na.rm = TRUE)

ggplot(q_error_df, aes(x = L1_scaled)) +
  geom_histogram(binwidth = 0.01, fill = "steelblue", color = "white") +
  geom_vline(xintercept = mean_L1, color = "red", linetype = "dashed", size = 1) +
  theme_minimal() +
  labs(
    title = "Q-matrix error distribution with mean",
    subtitle = paste("Mean L1_scaled =", round(mean_L1, 3)),
    x = "L1_scaled",
    y = "Count"
  )
```

```
ggplot(q_error_df, aes(x = Cosine)) +
  geom_histogram(binwidth = 0.01, fill = "darkorange", color = "white") +
  theme_minimal() +
  labs(
    title = "Distribution of Cosine similarity among duplicates",
    x = "Cosine similarity",
    y = "Number of duplicate pairs"
  )
```

```
# identify outliers
q_error_df %>%
  filter(L1_scaled > 0.3) %>%
  select(original, duplicate, L1_scaled, Cosine)
```

```
# summary statitics
q_error_df %>%
  summarise(
    mean_L1 = mean(L1_scaled, na.rm = TRUE),
    median_L1 = median(L1_scaled, na.rm = TRUE),
    sd_L1 = sd(L1_scaled, na.rm = TRUE),
    q05 = quantile(L1_scaled, 0.05, na.rm = TRUE),
    q95 = quantile(L1_scaled, 0.95, na.rm = TRUE)
  )
```

##### Genetic Diversity Scores

```
library(vcfR)
library(adegenet)
library(hierfstat)
library(poppr)
```

```
# Paths
vcf_file <- "FINAL_allipk_pst1ref_postfilter.recode.vcf"
vcf <- read.vcfR(vcf_file)
```

```
## Scanning file to determine attributes.
## File attributes:
##   meta lines: 63
##   header_line: 64
##   variant count: 2726
##   column count: 727
## Meta line 63 read in.
## All meta lines processed.
## gt matrix initialized.
## Character matrix gt created.
##   Character matrix gt rows: 2726
##   Character matrix gt cols: 727
##   skip: 0
##   nrows: 2726
##   row_num: 0
## Processed variant 1000Processed variant 2000Processed variant: 2726
## All variants processed
```

```
# remove outliers (as determined by PCA above)
outliers <- c("CALDERA_3243989-C-AH09.trimmed","CALDERA_3243963-C-AD08.trimmed","CALDERA_3243983-C-AF11.trimmed")
vcf <- vcf[, !colnames(vcf@gt) %in% outliers]

# filter for FIRST occurence of each unique plantID (i.e. remove duplicate samples)

unique_meta <- meta[!duplicated(meta$plantID), ]

vcf_samples <- colnames(vcf@gt)[-1]   # remove FORMAT column

keep <- intersect(unique_meta$sample, vcf_samples)

unique_meta <- unique_meta[match(keep, unique_meta$sample), ]

vcf_unique <- vcf[, c("FORMAT", keep)]

geno <- extract.gt(vcf_unique, element = "GT", as.numeric = TRUE)

# Build hierfstat data frame
hf <- data.frame(
  pop = unique_meta$population[match(colnames(geno), unique_meta$sample)],
  t(geno)  # transpose so individuals are rows
)
```

```
# count how many non-duplicate samples remained in vcf

table(unique_meta$population)
```

```
## 
##    Arica  Caldera Oyarbide 
##      138      348      146
```

###### Het-Stats

```
# Basic diversity stats per population
div_stats <- basic.stats(hf)

# Extract summaries
within_div <- data.frame(
  population = colnames(div_stats$Hs),
  He = colMeans(div_stats$Hs, na.rm = TRUE),
  Ho = colMeans(div_stats$Ho, na.rm = TRUE),
  FIS = colMeans(div_stats$Fis, na.rm = TRUE)
)

print(within_div)
```

```
##          population         He         Ho         FIS
## Arica         Arica 0.02413114 0.02849369 -0.03308523
## Caldera     Caldera 0.02583012 0.03015026 -0.05078721
## Oyarbide   Oyarbide 0.05541346 0.06986141 -0.11815136
```

He = Heterozygosity expected Ho = Heterozygosity observed FIS =
Inbreeding Coefficient in subpopulation

###### Fst-Stats

```
# Pairwise FST
fst_matrix <- pairwise.WCfst(hf)

# Convert to tidy table
between_fst <- as.data.frame(as.table(fst_matrix))
colnames(between_fst) <- c("Pop1", "Pop2", "FST")

print(between_fst)
```

```
##       Pop1     Pop2        FST
## 1    Arica    Arica         NA
## 2  Caldera    Arica 0.09915124
## 3 Oyarbide    Arica 0.14728921
## 4    Arica  Caldera 0.09915124
## 5  Caldera  Caldera         NA
## 6 Oyarbide  Caldera 0.22438391
## 7    Arica Oyarbide 0.14728921
## 8  Caldera Oyarbide 0.22438391
## 9 Oyarbide Oyarbide         NA
```

###### AMOVA

```
# Convert vcf to genlight object
gl <- vcfR2genlight(vcf_unique)

# Make sure population info is attached
pop(gl) <- unique_meta$population[match(indNames(gl), unique_meta$sample)]

# Define strata data frame
strata(gl) <- data.frame(
  population = pop(gl),
  Individual = indNames(gl)
)

amova_res <- poppr.amova(gl, ~population)

print(amova_res)
```

```
## $call
## ade4::amova(samples = xtab, distances = xdist, structures = xstruct)
## 
## $results
##                                     Df   Sum Sq    Mean Sq
## Between population                   2 105906.6 52953.3193
## Between samples Within population  629  96144.9   152.8536
## Within samples                     632 228019.5   360.7903
## Total                             1263 430071.0   340.5155
## 
## $componentsofcovariance
##                                                   Sigma         %
## Variations  Between population                 140.2332  35.31832
## Variations  Between samples Within population -103.9684 -26.18487
## Variations  Within samples                     360.7903  90.86655
## Total variations                               397.0552 100.00000
## 
## $statphi
##                                Phi
## Phi-samples-total       0.09133451
## Phi-samples-population -0.40482665
## Phi-population-total    0.35318319
```

###### Private and shared alleles per fields

```
# Suppose hf has genotypes coded as 0,1,2 (diploid SNPs)
# Convert to allele strings like "0/0", "0/1", "1/1"
geno_strings <- apply(hf[,-1], 2, function(x) {
  sapply(x, function(g) {
    if (is.na(g)) return(NA)
    switch(as.character(g),
           "0" = "A/A",
           "1" = "A/B",
           "2" = "B/B")
  })
})

geno_strings <- as.data.frame(geno_strings)

# generate gi from hf
gi <- df2genind(geno_strings, pop = hf$pop, ploidy = 2, type = "codom", sep = "/")
```

```
library(adegenet)

# Tabulate allele presence per individual
allele_tab <- tab(gi, freq = TRUE, NA.method = "zero")

# For each population, count how many individuals carry each allele
pop_levels <- levels(pop(gi))

allele_counts <- sapply(pop_levels, function(p) {
  idx <- which(pop(gi) == p)
  colSums(allele_tab[idx, , drop = FALSE] > 0)
})

# allele_counts is now a matrix: rows = alleles/loci, columns = populations
head(allele_counts)
```

```
##                         Oyarbide Caldera Arica
## dDocent_Contig_138_7.A       132     344   129
## dDocent_Contig_138_7.B         0       0     1
## dDocent_Contig_139_31.A      128     298   121
## dDocent_Contig_139_31.B        0       0     4
## dDocent_Contig_139_40.A      128     298   121
## dDocent_Contig_139_40.B        0       0     3
```

```
# Private if present in exactly one population
is_private <- rowSums(allele_counts > 0) == 1

# Shared if present in more than one population
is_shared <- rowSums(allele_counts > 0) > 1
```

```
private_totals <- colSums((allele_counts > 0)[is_private, ])
shared_totals  <- colSums((allele_counts > 0)[is_shared, ])
total_counts   <- colSums(allele_counts > 0)

allele_summary <- data.frame(
  Population = names(total_counts),
  Private = private_totals,
  Shared = shared_totals,
  Total = total_counts,
  Private_pct = round(private_totals / total_counts * 100, 2),
  Shared_pct = round(shared_totals / total_counts * 100, 2)
)

print(allele_summary)
```

```
##          Population Private Shared Total Private_pct Shared_pct
## Oyarbide   Oyarbide     423   3298  3721       11.37      88.63
## Caldera     Caldera     479   3193  3672       13.04      86.96
## Arica         Arica     519   3514  4033       12.87      87.13
```

##### Identify clones

```
# Suppose hf has genotypes coded as 0,1,2 (diploid SNPs)
# Convert to allele strings like "0/0", "0/1", "1/1"
geno_strings <- apply(hf[,-1], 2, function(x) {
  sapply(x, function(g) {
    if (is.na(g)) return(NA)
    switch(as.character(g),
           "0" = "A/A",
           "1" = "A/B",
           "2" = "B/B")
  })
})

geno_strings <- as.data.frame(geno_strings)

# generate gi from hf
gi <- df2genind(geno_strings, pop = hf$pop, ploidy = 2, type = "codom", sep = "/")

gc <- as.genclone(gi)     # convert genind → genclone
```

```
# 1) Compute SNP-wise Hamming distance (bitwise differences)
d <- bitwise.dist(gl)  # fast, suitable for SNPs

# 2) Inspect the distance distribution to choose a threshold
hist(d, breaks = 50, main = "Pairwise SNP distances")
```

###### Number of genets (multi-locus-genotypes) per similarity threshold

```
library(poppr)

# Pairwise SNP distance (already computed as 'd')
thresholds <- seq(0.01, 0.15, by = 0.01)

# Total MLG counts per threshold
mlg_counts <- sapply(thresholds, function(thr) {
  mlg_assignments <- mlg.filter(gl, distance = d, threshold = thr, stats = "MLG")
  length(unique(mlg_assignments))
})

plot(thresholds, mlg_counts, type = "b", pch = 19,
     xlab = "Distance threshold", ylab = "Number of MLGs",
     main = "MLG count vs. threshold")
```

```
# Returns a tidy data frame with counts per population and threshold
per_field <- do.call(rbind, lapply(thresholds, function(thr) {
  mlg_assignments <- mlg.filter(gl, distance = d, threshold = thr, stats = "MLG")
  data.frame(
    threshold = thr,
    population = pop(gl),
    mlg = mlg_assignments
  )
}))

# Summarize unique MLGs per field at each threshold
library(dplyr)
summary_per_field <- per_field %>%
  group_by(threshold, population) %>%
  summarise(unique_mlg = n_distinct(mlg), .groups = "drop")

print(summary_per_field)
```

```
## # A tibble: 45 × 3
##    threshold population unique_mlg
##        <dbl> <fct>           <int>
##  1      0.01 Arica             136
##  2      0.01 Caldera           347
##  3      0.01 Oyarbide          146
##  4      0.02 Arica             123
##  5      0.02 Caldera           324
##  6      0.02 Oyarbide          142
##  7      0.03 Arica             107
##  8      0.03 Caldera           275
##  9      0.03 Oyarbide          123
## 10      0.04 Arica              92
## # ℹ 35 more rows
```

#### Local analyses

same denovo reference, repeat bwa mapping and freebayes SNP calling
and filtering for each field separately

```
run_analysis <- function(field_dir, prefix, Kvals, chosenK) {
  library(vcfR)
  library(dplyr)
  library(tidyverse)
  library(pheatmap)
  library(poppr)
  library(scatterpie)
  library(ggthemes)

  # --- Load VCF ---

vcf_file <- list.files(field_dir, pattern = "\\.vcf$", full.names = TRUE)
if (length(vcf_file) == 0) {
  stop("No VCF file found in ", field_dir)
}
vcf <- read.vcfR(vcf_file[1])   # take the first match

gl  <- vcfR2genlight(vcf)

# Remove outlier individuals from the genlight object
outliers <- c("CALDERA_3243989-C-AH09.trimmed","CALDERA_3243963-C-AD08.trimmed","CALDERA_3243983-C-AF11.trimmed")
gl <- gl[!indNames(gl) %in% outliers, ]

ploidy(gl) <- 2

# convert to matrix
geno_matrix <- as.matrix(gl)

# add rownames
rownames(geno_matrix) <- indNames(gl)

# attach population info
pop(gl) <- meta$population[match(indNames(gl), meta$sample)]


  # --- PCA ---
  geno <- read_delim(file.path(field_dir, "pca_input.raw"), delim = " ")
# Drop non-genotype columns
geno_clean <- geno %>%
  select(-FID, -PAT, -MAT, -SEX, -PHENOTYPE) %>%
  column_to_rownames("IID")

# Remove SNPs with too much missing data
geno_clean <- geno_clean %>% select(where(~ mean(is.na(.)) < 0.2))

# Impute missing values with mean
geno_imputed <- geno_clean %>%
  mutate(across(everything(), ~ ifelse(is.na(.), mean(., na.rm = TRUE), .)))

# Remove constant (zero-variance) SNPs
geno_imputed <- geno_imputed %>% select(where(~ var(.) > 0))

# Run PCA
pca <- prcomp(geno_imputed, scale. = TRUE)
scores <- as_tibble(pca$x) %>%
    mutate(sample = rownames(pca$x)) %>%
    left_join(meta, by = "sample")

  # --- PCA plot ---
  p_pca <- ggplot(scores, aes(x = PC1, y = PC2, color = population, shape = category)) +
    geom_point(size = 3, alpha = 0.8) +
    theme_minimal() +
    labs(title = paste("PCA:", basename(field_dir)))
  print(p_pca)

  # --- Heatmap of genetic distances ---
  d <- bitwise.dist(gl)
  d_mat <- as.matrix(d)
  annotation <- data.frame(population = pop(gl))
  rownames(annotation) <- indNames(gl)
  p_heatmap <- pheatmap(d_mat,
           annotation_row = annotation,
           annotation_col = annotation,
           show_colnames = FALSE,
           show_rownames = FALSE,
           clustering_distance_rows = "euclidean",
           clustering_distance_cols = "euclidean",
           clustering_method = "average",
           color = colorRampPalette(c("blue", "white", "red"))(100),
           main = paste("Genetic distances:", basename(field_dir)))
  print(p_heatmap)

  # --- ADMIXTURE CV error plots ---
logs <- list.files(field_dir, pattern = "log[0-9]+\\.out", full.names = TRUE)

cv_df <- purrr::map_dfr(logs, function(f) {
  lines <- readLines(f)
  cv_line <- grep("CV error", lines, value = TRUE)
  k <- as.integer(stringr::str_match(cv_line, "CV error \\(K=(\\d+)\\)")[,2])
  cv <- as.numeric(stringr::str_match(cv_line, ": ([0-9\\.]+)")[,2])
  tibble(K = k, CV = cv)
})

p_cv <- ggplot(cv_df, aes(x = K, y = CV)) +
  geom_line(color = "steelblue") +
  geom_point(size = 2, color = "darkred") +
  geom_point(data = cv_df %>% filter(CV == min(CV)),
             aes(x = K, y = CV), color = "green", size = 3) +
  labs(title = paste("ADMIXTURE CV Error:", basename(field_dir)),
       x = "Number of clusters (K)",
       y = "Cross-validation error") +
  theme_minimal()

print(p_cv)

  
  # --- ADMIXTURE ---
  fam <- read_table2(file.path(field_dir, paste0(prefix, ".fam")), col_names = FALSE)
  sample_ids <- fam$X2

  read_Q <- function(K) {
    qfile <- file.path(field_dir, paste0(prefix, ".", K, ".Q"))
    Q <- read_table2(qfile, col_names = FALSE)
    colnames(Q) <- paste0("Cluster", 1:K)
    Qdf <- cbind(sample = sample_ids, Q)
    Qdf$K <- K
    Qdf
  }

  Qall <- map_df(Kvals, read_Q) %>%
    left_join(meta, by = "sample", unmatched = "drop") %>%
    arrange(population, elevation)
  
  outliers <- c("CALDERA_3243989-C-AH09.trimmed","CALDERA_3243963-C-AD08.trimmed","CALDERA_3243983-C-AF11.trimmed")
  Qall <- Qall %>% filter(!sample %in% outliers)

  Qlong <- Qall %>%
    pivot_longer(starts_with("Cluster"),
                 names_to = "Cluster",
                 values_to = "Ancestry") %>%
    mutate(Cluster = factor(Cluster, levels = paste0("Cluster", 1:max(Kvals))),
           sample = factor(sample, levels = unique(sample)))

  cluster_colors <- viridis(max(Kvals), option = "D")
  names(cluster_colors) <- paste0("Cluster", 1:max(Kvals))

  # --- Plot all Ks ---
  p_allK <- ggplot(Qlong, aes(x = sample, y = Ancestry, fill = Cluster)) +
    geom_bar(stat = "identity", width = 1) +
    facet_grid(K ~ population, scales = "free_x", space = "free_x") +
    theme_bw() +
    labs(title = paste("ADMIXTURE results:", basename(field_dir))) +
    scale_fill_manual(values = cluster_colors)
  print(p_allK)

  # --- Focused plot for chosen K ---
  Qchosen <- Qlong %>% filter(K == chosenK)
  p_chosenK <- ggplot(Qchosen, aes(x = sample, y = Ancestry, fill = Cluster)) +
    geom_bar(stat = "identity", width = 1) +
    facet_wrap(~population, scales = "free_x", nrow = 1) +
    theme_bw() +
    labs(title = paste("ADMIXTURE results at K =", chosenK, ":", basename(field_dir)))
  print(p_chosenK)

  # --- Pie maps ---
  pie_data <- Qall %>%
  filter(K == chosenK) %>%
  mutate(
    latitude  = as.numeric(latitude),
    longitude = as.numeric(longitude)
  ) %>%
  select(sample, latitude, longitude, population, starts_with("Cluster"))

  pal <- cluster_colors[paste0("Cluster", 1:chosenK)]
  for (pop in unique(pie_data$population)) {
    pie_subset <- pie_data %>% filter(population == pop)
    p <- ggplot() +
      geom_scatterpie(aes(x = longitude, y = latitude),
                      data = pie_subset,
                      cols = paste0("Cluster", 1:chosenK),
                      pie_scale = 0.8) +
      coord_equal() +
      theme_minimal() +
      labs(title = paste("ADMIXTURE pie map:", pop))
    print(p)
  }

  write.csv(pie_data, file.path(field_dir, paste0("piechart_data_K", chosenK, basename(field_dir),".csv")))
}
```

```
fields <- list(
  Arica   = list(dir = "onlyArica",   prefix = "admix_03_pruned",   Kvals = 2:10, chosenK = 3),
  Iquique = list(dir = "onlyIquique", prefix = "admix_03_pruned", Kvals = 2:10, chosenK = 3),
  Caldera = list(dir = "onlyCaldera", prefix = "admix_03_pruned", Kvals = 2:10, chosenK = 4)
)

for (f in names(fields)) {
  params <- fields[[f]]
  run_analysis(field_dir = params$dir,
               prefix    = params$prefix,
               Kvals     = params$Kvals,
               chosenK   = params$chosenK)
}
```

```
## Scanning file to determine attributes.
## File attributes:
##   meta lines: 63
##   header_line: 64
##   variant count: 3772
##   column count: 179
## Meta line 63 read in.
## All meta lines processed.
## gt matrix initialized.
## Character matrix gt created.
##   Character matrix gt rows: 3772
##   Character matrix gt cols: 179
##   skip: 0
##   nrows: 3772
##   row_num: 0
## Processed variant 1000Processed variant 2000Processed variant 3000Processed variant: 3772
## All variants processed
```

```
## Scanning file to determine attributes.
## File attributes:
##   meta lines: 63
##   header_line: 64
##   variant count: 8068
##   column count: 232
## Meta line 63 read in.
## All meta lines processed.
## gt matrix initialized.
## Character matrix gt created.
##   Character matrix gt rows: 8068
##   Character matrix gt cols: 232
##   skip: 0
##   nrows: 8068
##   row_num: 0
## Processed variant 1000Processed variant 2000Processed variant 3000Processed variant 4000Processed variant 5000Processed variant 6000Processed variant 7000Processed variant 8000Processed variant: 8068
## All variants processed
```

```
## Scanning file to determine attributes.
## File attributes:
##   meta lines: 63
##   header_line: 64
##   variant count: 417
##   column count: 487
## Meta line 63 read in.
## All meta lines processed.
## gt matrix initialized.
## Character matrix gt created.
##   Character matrix gt rows: 417
##   Character matrix gt cols: 487
##   skip: 0
##   nrows: 417
##   row_num: 0
## Processed variant: 417
## All variants processed
```

### B) Sand Data Processing

```
library(patchwork)
library(tidyverse)
library(ggpubr)
library(Ternary)
library(readxl)
library(ggtern)
library(scales)
library(dplyr)
library(purrr)
library(rstatix)
```

```
# import data
Sanddata <- read_excel("Sanddata.xlsx")
```

```
# data tidying
Sanddata <- Sanddata %>%
  mutate(
    sorting_category = factor(
      sorting_category,
      levels = c(
        "very well sorted",
        "well sorted",
        "moderately well sorted",
        "moderately sorted",
        "poorly sorted",
        "very poorly sorted"
      )
    )
  )
```

```
give.n <- function(x){
   return(c(y = -1, label = length(x)))
}

give.median <- function(x){
   return(c(y = median(x)+(median(x)*0.10), label = round(median(x),2)))
}

symnum.args <- list(cutpoints = c(0, 0.0001, 0.001, 0.01, 0.05, 1), symbols = c("****", "***", "**", "*", "ns"))
```

```
Sanddata <- Sanddata %>%
  mutate(grainsize_mean = as.numeric(as.character(grainsize_mean))) %>%
  filter(!is.na(grainsize_mean), !is.na(fieldID))
```

```
Sanddata <- Sanddata %>%
  mutate(sortingidx = as.numeric(as.character(sortingidx))) %>%
  filter(!is.na(sortingidx), !is.na(fieldID))
```

#### Boxplot1 - Field - Grainsize

```
Sanddata$plantstate <- factor(Sanddata$plantstate,
                              levels = c("PLANT", "DEAD", "NO_PLANT"))

Sanddata$fieldID <- factor(Sanddata$fieldID,
                           levels = c("Arica", "Oyarbide", "Caldera"))

p<-Sanddata %>% ggplot(aes(x=fieldID,y=grainsize_mean,fill=fieldID))+
  geom_boxplot(
  alpha = 0.2,
  outlier.shape = 16,
  outlier.size = 1.5,
  outlier.position = position_jitter(width = 0.1),
  coef = 1
)+
  stat_compare_means(na.rm=T, method = "wilcox.test", exact=TRUE, symnum.args = symnum.args, comparisons = list(c("Caldera", "Oyarbide"), c("Oyarbide", "Arica"), c("Arica", "Caldera")),
  step.increase = 0.07)+
  facet_grid(~plantstate)+
  stat_summary(fun.data = give.n, geom = "text")+
  geom_hline(yintercept = c(30, 63, 125, 250), color = "red", linetype = "dashed")+
    theme(strip.text = element_text(size = 10),
      strip.background = element_rect(color = NA),
      panel.spacing = unit(0.2, "lines"))
p
```

```
#ggsave("Sanddata_grainsize_boxplot1.pdf", plot = p, width = 8, height = 5, units = "in")
```

#### Boxplot2 - Plant Occurence - Grainsize

```
Sanddata$plantstate <- factor(Sanddata$plantstate,
                              levels = c("PLANT", "DEAD", "NO_PLANT"))
Sanddata$fieldID <- factor(Sanddata$fieldID,
                           levels = c("Arica", "Oyarbide", "Caldera"))

p<-Sanddata %>% ggplot(aes(x=plantstate,y=grainsize_mean,fill=plantstate))+
  geom_boxplot(
  alpha = 0.2,
  outlier.shape = 16,
  outlier.size = 1.5,
  outlier.position = position_jitter(width = 0.1),
  coef = 1)+
  ggpubr::stat_compare_means(na.rm=T, method = "wilcox.test", exact=TRUE, symnum.args = symnum.args, comparisons = list(c("DEAD", "PLANT"), c("PLANT", "NO_PLANT"), c("NO_PLANT", "DEAD")),
                     step.increase = 0.07)+
  facet_grid(~fieldID)+
  stat_summary(fun.data = give.n, geom = "text")+
  geom_hline(yintercept = c(30, 63, 125, 250), color = "red", linetype = "dashed")+
   theme(strip.text = element_text(size = 10),
      strip.background = element_rect(color = NA),
      panel.spacing = unit(0.2, "lines"))
p
```

```
#ggsave("Sanddata_grainsize_boxplot2.pdf", plot = p, width = 8, height = 5, units = "in")
```

#### Boxplot3 - Field ID - Sorting Index

```
Sanddata$plantstate <- factor(Sanddata$plantstate,
                              levels = c("PLANT", "DEAD", "NO_PLANT"))
Sanddata$fieldID <- factor(Sanddata$fieldID,
                           levels = c("Arica", "Oyarbide", "Caldera"))

p<-Sanddata %>% ggplot(aes(x=fieldID,y=sortingidx,fill=fieldID))+
  geom_boxplot(
  alpha = 0.2,
  outlier.shape = 16,
  outlier.size = 1.5,
  outlier.position = position_jitter(width = 0.1),
  coef = 1)+
  stat_compare_means(na.rm=T, method = "wilcox.test", exact=TRUE, symnum.args = symnum.args, comparisons = list(c("Caldera", "Oyarbide"), c("Oyarbide", "Arica"), c("Arica", "Caldera")),
                     step.increase = 0.07)+
  facet_grid(~plantstate)+
  stat_summary(fun.data = give.n, geom = "text")+
   geom_hline(yintercept = c(4, 2, 1.62, 1.41, 1.27, 0.0), color = "red", linetype = "dashed")+
   theme(strip.text = element_text(size = 10),
      strip.background = element_rect(color = NA),
      panel.spacing = unit(0.2, "lines"))
p
```

```
#ggsave("Sanddata_grainsize_boxplot3.pdf", plot = p, width = 8, height = 5, units = "in")
```

#### Boxplot4 - Plant Occurence- Sorting Index

```
Sanddata$plantstate <- factor(Sanddata$plantstate,
                              levels = c("PLANT", "DEAD", "NO_PLANT"))
Sanddata$fieldID <- factor(Sanddata$fieldID,
                           levels = c("Arica", "Oyarbide", "Caldera"))

p<-Sanddata %>% ggplot(aes(x=plantstate,y=sortingidx,fill=plantstate))+
  geom_boxplot(alpha = 0.2,
  outlier.shape = 16,
  outlier.size = 1.5,
  outlier.position = position_jitter(width = 0.1),
  coef = 1)+
  stat_compare_means(na.rm=T, method = "wilcox.test", exact=TRUE, symnum.args = symnum.args, comparisons = list(c("DEAD", "PLANT"), c("PLANT", "NO_PLANT"), c("NO_PLANT", "DEAD")),
                     step.increase = 0.07)+
  facet_grid(~fieldID)+
  stat_summary(fun.data = give.n, geom = "text")+
  geom_hline(yintercept = c(4, 2, 1.62, 1.41, 1.27, 0.0), color = "red", linetype = "dashed")+
     theme(strip.text = element_text(size = 10),
      strip.background = element_rect(color = NA),
      panel.spacing = unit(0.2, "lines"))
p
```

```
#ggsave("Sanddata_grainsize_boxplot4.pdf", plot = p, width = 8, height = 5, units = "in")
```

#### Statistical Analysis

Note for all statistical analysis: wilcox.test (package: ggpubr)
function will not write r - values for comparisions with low sample
sizes in the dataset. Therefore, r and p values where additionally
calculated with the same formular by hand. Values based on low sample
size were not used for further statistical interpretation.

```
# Grainsize: Wilcoxon Test ( calculation of p and r values ) with Bonferroni Holm adjustion of p values
# p values Grainsize - Field

library(ggpubr)
library(dplyr)
library(rstatix)
library(purrr)

my_comparisons <- list(
  c("Caldera", "Oyarbide"),
  c("Oyarbide", "Arica"),
  c("Arica", "Caldera")
)

# compute Wilcoxon tests for each plantstate (facet)
p_values_table <- compare_means(
  grainsize_mean ~ fieldID,
  group.by = "plantstate",
  data = Sanddata,
  comparisons = my_comparisons,
  method = "wilcox.test",
  exact = FALSE,
  na.rm = TRUE,
  p.adjust.method = "holm"
)

# optional: round p-values
p_values_table$p <- signif(p_values_table$p, 3)

p_values_table
```

```
write.csv(p_values_table, "p_values_table_gs", row.names = FALSE)
```

```
# additional r and Z values Grainsize - Field
# filte the single comparisions
valid_comps <- function(df, comps) {
  present <- unique(df$fieldID)
  Filter(function(x) all(x %in% present), comps)
}

# raw Wilcoxon-Tests
wilcox_raw <- Sanddata %>%
  group_split(plantstate) %>%
  set_names(unique(Sanddata$plantstate)) %>%
  map(function(df) {
    comps <- valid_comps(df, my_comparisons)
    if (length(comps) == 0) return(NULL)

    df %>%
      wilcox_test(
        grainsize_mean ~ fieldID,
        comparisons = comps,
        exact = FALSE,
        detailed = TRUE   # wichtig!
      )
  })

wilcox_raw
```

```
## $PLANT
## # A tibble: 3 × 14
##   estimate .y.            group1 group2    n1    n2 statistic         p conf.low
## *    <dbl> <chr>          <chr>  <chr>  <int> <int>     <dbl>     <dbl>    <dbl>
## 1   -131.  grainsize_mean Oyarb… Calde…   219   488        20 2.06e-100   -138. 
## 2     81.2 grainsize_mean Arica  Oyarb…   278   219     52348 3.28e- 43     71.0
## 3    -53.4 grainsize_mean Arica  Calde…   278   488     33000 2.78e- 32    -62.0
## # ℹ 5 more variables: conf.high <dbl>, method <chr>, alternative <chr>,
## #   p.adj <dbl>, p.adj.signif <chr>
## 
## $NO_PLANT
## # A tibble: 1 × 12
##   estimate .y.      group1 group2    n1    n2 statistic     p conf.low conf.high
## *    <dbl> <chr>    <chr>  <chr>  <int> <int>     <dbl> <dbl>    <dbl>     <dbl>
## 1     12.6 grainsi… Arica  Oyarb…    32    12       247 0.151    -8.13      41.0
## # ℹ 2 more variables: method <chr>, alternative <chr>
## 
## $DEAD
## # A tibble: 3 × 14
##   estimate .y.    group1 group2    n1    n2 statistic       p conf.low conf.high
## *    <dbl> <chr>  <chr>  <chr>  <int> <int>     <dbl>   <dbl>    <dbl>     <dbl>
## 1   -156.  grain… Oyarb… Calde…   134     4         0 6.87e-4   -186.     -134. 
## 2     46.9 grain… Arica  Oyarb…    79   134      7452 6.77e-7     28.3      61.5
## 3   -114.  grain… Arica  Calde…    79     4        10 2   e-3   -175.      -55.3
## # ℹ 4 more variables: method <chr>, alternative <chr>, p.adj <dbl>,
## #   p.adj.signif <chr>
```

```
#calculate r and Z values by hand
wilcox_with_r <- wilcox_raw %>%
  map_df(~ .x %>%
           rowwise() %>%
           mutate(
             mu_W = n1 * n2 / 2,
             sigma_W = sqrt(n1 * n2 * (n1 + n2 + 1) / 12),
             Z = (statistic - mu_W) / sigma_W,
             r = Z / sqrt(n1 + n2),
             r = signif(r, 3)
           ) %>%
           ungroup() %>%
          select(group1, group2, n1, n2, statistic, r, p)
         )
wilcox_with_r
```

```
write.csv(wilcox_with_r, "r_values_gs", row.names = FALSE)
```

```
# p values: Sorting Index - Field
my_comparisons <- list(
  c("Caldera", "Oyarbide"),
  c("Oyarbide", "Arica"),
  c("Arica", "Caldera")
)

# compute Wilcoxon tests for each plantstate (facet)
p_values_table_sort <- compare_means(
  sortingidx ~ fieldID,
  group.by = "plantstate",
  data = Sanddata,
  comparisons = my_comparisons,
  method = "wilcox.test",
  exact = FALSE,
  na.rm = TRUE,
  p.adjust.method = "holm"
)

#optional: round p-values
p_values_table_sort$p <- signif(p_values_table$p, 3)

p_values_table_sort
```

```
write.csv(p_values_table_sort, "p_values_table_sort", row.names = FALSE)
```

```
# additional r and Z values Sorting - Field
# filter the single comparisions
valid_comps <- function(df, comps) {
  present <- unique(df$fieldID)
  Filter(function(x) all(x %in% present), comps)
}

# only raw wilcoxon test
wilcox_raw <- Sanddata %>%
  group_split(plantstate) %>%
  set_names(unique(Sanddata$plantstate)) %>%
  map(function(df) {
    comps <- valid_comps(df, my_comparisons)
    if (length(comps) == 0) return(NULL)

    df %>%
      wilcox_test(
        sortingidx ~ fieldID,
        comparisons = comps,
        exact = FALSE,
        detailed = TRUE
      )
  })

wilcox_raw
```

```
## $PLANT
## # A tibble: 3 × 14
##   estimate .y.   group1 group2    n1    n2 statistic        p conf.low conf.high
## *    <dbl> <chr> <chr>  <chr>  <int> <int>     <dbl>    <dbl>    <dbl>     <dbl>
## 1    0.229 sort… Oyarb… Calde…   219   488     69901 5.50e-11    0.130     0.301
## 2   -0.275 sort… Arica  Oyarb…   278   219     25205 9.89e- 4   -0.414    -0.123
## 3    0.145 sort… Arica  Calde…   278   488     89454 2.10e-13    0.110     0.184
## # ℹ 4 more variables: method <chr>, alternative <chr>, p.adj <dbl>,
## #   p.adj.signif <chr>
## 
## $NO_PLANT
## # A tibble: 1 × 12
##   estimate .y.    group1 group2    n1    n2 statistic       p conf.low conf.high
## *    <dbl> <chr>  <chr>  <chr>  <int> <int>     <dbl>   <dbl>    <dbl>     <dbl>
## 1    0.520 sorti… Arica  Oyarb…    32    12       295 0.00691    0.144     0.711
## # ℹ 2 more variables: method <chr>, alternative <chr>
## 
## $DEAD
## # A tibble: 3 × 14
##   estimate .y.    group1 group2    n1    n2 statistic       p conf.low conf.high
## *    <dbl> <chr>  <chr>  <chr>  <int> <int>     <dbl>   <dbl>    <dbl>     <dbl>
## 1    0.385 sorti… Oyarb… Calde…   134     4       510 2   e-3   0.0987     1.66 
## 2    0.457 sorti… Arica  Oyarb…    79   134      7138 2.18e-5   0.241      0.739
## 3    1.25  sorti… Arica  Calde…    79     4       292 5   e-3   0.362      2.48 
## # ℹ 4 more variables: method <chr>, alternative <chr>, p.adj <dbl>,
## #   p.adj.signif <chr>
```

```
wilcox_with_r <- wilcox_raw %>%
  map_df(~ .x %>%
           rowwise() %>%
           mutate(
             mu_W = n1 * n2 / 2,
             sigma_W = sqrt(n1 * n2 * (n1 + n2 + 1) / 12),
             Z = (statistic - mu_W) / sigma_W,
             r = Z / sqrt(n1 + n2),
             r = signif(r, 3)
           ) %>%
           ungroup() %>%
          select(group1, group2, n1, n2, statistic, r, p)
         )
wilcox_with_r
```

```
write.csv(wilcox_with_r, "r_values_sort", row.names = FALSE)
```

```
# p values: Grainsize - Plant Occurence
my_comparisons <- list(
  c("DEAD", "PLANT"),
  c("PLANT", "NO_PLANT"),
  c("NO_PLANT", "DEAD")
)

# compute Wilcoxon tests for each fieldID (facet)
p_values_fieldID <- compare_means(
  grainsize_mean ~ plantstate,
  group.by = "fieldID",
  data = Sanddata,
  comparisons = my_comparisons,
  method = "wilcox.test",
  exact = FALSE,
  na.rm = TRUE,
  p.adjust.method = "holm"
)

# optional: round p-values
p_values_fieldID$p <- signif(p_values_table$p, 3)

p_values_fieldID
```

```
write.csv(p_values_fieldID, "p_values_fieldID.csv", row.names = FALSE)
```

```
# Additional r and Z values Grainsize - Plant Occurence

# --- 1. Plantstate-Paare ---
my_comparisons <- list(
  c("DEAD", "PLANT"),
  c("PLANT", "NO_PLANT"),
  c("NO_PLANT", "DEAD")
)

# --- 2. Wilcoxon-Test pro FieldID (mit grainsize_mean) ---
wilcox_raw_grain <- Sanddata %>%
  group_split(fieldID) %>%
  set_names(unique(Sanddata$fieldID)) %>%
  map(function(df) {
    #only if plantstates are available in the coomparisons
    df <- df %>%
      filter(plantstate %in% unlist(my_comparisons)) %>%
      droplevels()
    
    # observations are not 0
    valid <- Filter(function(x) all(sapply(x, function(g) sum(df$plantstate == g) >= 2)),
                    my_comparisons)
    if(length(valid) == 0) return(NULL)
    
    #Wilcoxon-Test (raw)
    res <- df %>%
      wilcox_test(grainsize_mean ~ plantstate,
                  comparisons = valid,
                  exact = FALSE,
                  detailed = TRUE) %>%
      mutate(fieldID = unique(df$fieldID))
    
    res
  })

# calculate Z and r values
wilcox_with_r_fieldID_grain <- wilcox_raw_grain %>%
  compact() %>%
  map_df(~ .x %>%
           rowwise() %>%
           mutate(
             mu_W = n1 * n2 / 2,
             sigma_W = sqrt(n1 * n2 * (n1 + n2 + 1) / 12),
             Z = (statistic - mu_W) / sigma_W,
             r = Z / sqrt(n1 + n2),
             r = signif(r, 3)
           ) %>%
           ungroup() %>%
           select(fieldID, group1, group2, n1, n2, statistic, r, p)
  )

wilcox_with_r_fieldID_grain
```

```
write.csv(wilcox_with_r_fieldID_grain,
          "wilcox_with_r_fieldID_grain.csv",
          row.names = FALSE)
```

```
# p-values Sorting Index - Plant Occurence
my_comparisons <- list(
  c("DEAD", "PLANT"),
  c("PLANT", "NO_PLANT"),
  c("NO_PLANT", "DEAD")
)

# compute Wilcoxon tests for each fieldID (facet)
p_values_fieldID_sorting <- compare_means(
  sortingidx ~ plantstate,
  group.by = "fieldID",
  data = Sanddata,
  comparisons = my_comparisons,
  method = "wilcox.test",
  exact = FALSE,
  na.rm = TRUE,
  p.adjust.method = "holm"
)

# optional: round p-values
p_values_fieldID_sorting$p <- signif(p_values_table$p, 3)

p_values_fieldID_sorting
```

```
write.csv(p_values_fieldID_sorting, "p_values_fieldID_sorting.csv", row.names = FALSE)
```

```
# Additional r and Z values Sortin Index - Plant Occurence
# Plantstate-Paare
my_comparisons <- list(
  c("DEAD", "PLANT"),
  c("PLANT", "NO_PLANT"),
  c("NO_PLANT", "DEAD")
)

# Wilcoxon-Test
wilcox_raw <- Sanddata %>%
  group_split(fieldID) %>%
  set_names(unique(Sanddata$fieldID)) %>%
  map(function(df) {
    #only if plantstates are available in the coomparisons
    df <- df %>%
      filter(plantstate %in% unlist(my_comparisons)) %>%
      droplevels()
    
    #not 0 observations
    valid <- Filter(function(x) all(sapply(x, function(g) sum(df$plantstate == g) >= 2)),
                    my_comparisons)
    if(length(valid) == 0) return(NULL)
    
    # Wilcoxon-Test (raw)
    res <- df %>%
      wilcox_test(sortingidx ~ plantstate,
                  comparisons = valid,
                  exact = FALSE,
                  detailed = TRUE) %>%
      mutate(fieldID = unique(df$fieldID))
    
    res
  })

# calculate addiotinal Z- und r- values 
wilcox_with_r_fieldID_sorting <- wilcox_raw %>%
  compact() %>%
  map_df(~ .x %>%
           rowwise() %>%
           mutate(
             mu_W = n1 * n2 / 2,
             sigma_W = sqrt(n1 * n2 * (n1 + n2 + 1) / 12),
             Z = (statistic - mu_W) / sigma_W,
             r = Z / sqrt(n1 + n2),
             r = signif(r, 3)
           ) %>%
           ungroup() %>%
           select(fieldID, group1, group2, n1, n2, statistic, r, p)
  )

wilcox_with_r_fieldID_sorting
```

```
write.csv(wilcox_with_r_fieldID_sorting,
          "wilcox_with_r_fieldID_sorting.csv",
          row.names = FALSE)
```

```
# Mean and Variance calculation for all data

mean_data <- Sanddata %>%
  group_by(fieldID) %>%
  summarise(
    mean_grain_size = mean(grainsize_mean, na.rm = TRUE),
    mean_sorting_idx = mean(sortingidx, na.rm = TRUE)
  )
```

```
#Kruskal test (comparison of the mean grain size between all three fields (Arica, Oyarbide, Caldera)
Sanddata %>%
  kruskal_test(grainsize_mean ~ fieldID)
```

```
Sanddata %>%
  kruskal_effsize(grainsize_mean ~ fieldID)
```

### C) Weather and Terrain Data Processing

```
library(tidyverse)
library(readxl)
library(GGally)
library(ggpubr)
library(gridExtra)
library(chron)
library(lubridate)
library(hms)
library(readr)
library(fmsb)
library(scales)
library(purrr)
library(stringr)
library(RColorBrewer)
library(patchwork)
library(dplyr)
library(grid)
library(corrplot)
```

```
weatherdata <- read.csv("WindTempData.csv", sep=";")
```

```
# Convert to Chilean time
## assign class to DateTime

weatherdata$DateTimeGMT0 <- as.POSIXct(weatherdata$DateTimeGMT0, format = "%d.%m.%Y %H:%M")
weatherdata$DateTimeCL <- with_tz(weatherdata$DateTimeGMT0, tzone = "America/Santiago")
```

```
# Create new columns
weatherdata$ymd      <- ymd(format(weatherdata$DateTimeCL, "%Y-%m-%d"))   # yyyy-mm-dd
weatherdata$ym        <- format(weatherdata$DateTimeCL, "%Y-%m")      # yyyy-mm
weatherdata$month <- month(weatherdata$DateTimeCL, label = TRUE, abbr = TRUE)         # mm
weatherdata$hms       <- format(weatherdata$DateTimeCL, "%H:%M:%S")  # hh:mm:ss
weatherdata$year <- year(weatherdata$ymd)

head(weatherdata)
```

```
## FILTER DATA
weatherdata <- weatherdata %>% filter(is.na(DateTimeCL) == FALSE)
```

```
# mask defect sensor values (negative values)

weatherdata <- weatherdata %>%
  mutate(across(where(is.numeric), ~ ifelse(. < 0, NA, .)))
```

```
# mean per month
monthmeans <- weatherdata %>% group_by(Region,Station,year,ym,month) %>% summarise(across(where(is.numeric),mean,na.rm=T))
head(monthmeans)
```

```
# mean per day
daymeans <- weatherdata %>% group_by(Region,Station,year,ymd) %>% summarise(across(where(is.numeric),mean,na.rm=T))
head(daymeans)
```

```
weatherdata_long <- weatherdata %>%
  pivot_longer(
    cols = -c(Region,Station,`DateTimeGMT0`,DateTimeCL,ymd,ym,month,hms,year),
    names_to = "parameter",
    values_to = "value"
  )
```

#### Wind speeds and directions

```
wind <- weatherdata_long %>% filter(parameter=="Winddir" | parameter=="Windspeed") %>% pivot_wider(names_from=parameter,values_from = value) %>% filter(Winddir>=0,Windspeed>=0.01)

# assign order
wind <- wind %>%
  mutate(Station = factor(Station,
                          levels = c("WSA1", "WSA2", "OYA1128","OYA1211","MSC3","MSC1")))
```

```
make_windrose <- function(wind,
                          spd_col = "Windspeed",
                          dir_col = "Winddir",
                          group_vars = "Station",
                          spdres = 2,
                          dirres = 22.5,
                          palette = "YlGnBu") {
  
  # dynamic binning based on max observed speed
  max_speed <- ceiling(max(wind[[spd_col]], na.rm = TRUE))
  spd.breaks <- c(seq(0, max_speed, by = spdres), Inf)
  spd.labels <- c(paste0(seq(0, max_speed - spdres, by = spdres), "-",
                         seq(spdres, max_speed, by = spdres)),
                  paste0(max_speed, "+"))
  
  # direction bins
  dir.breaks <- c(seq(0, 360, by = dirres))
  dir.labels <- c("N","NNE","NE","ENE","E","ESE","SE","SSE",
                  "S","SSW","SW","WSW","W","WNW","NW","NNW")
  
  wind_binned <- wind %>%
    filter(!is.na(.data[[spd_col]]), !is.na(.data[[dir_col]])) %>%
    mutate(
      spd.binned = cut(.data[[spd_col]],
                       breaks = spd.breaks,
                       labels = spd.labels,
                       right = FALSE,
                       include.lowest = TRUE),
      dir.binned = cut(.data[[dir_col]],
                       breaks = dir.breaks,
                       labels = dir.labels,
                       right = FALSE,
                       include.lowest = TRUE),
      spd.binned = factor(spd.binned, levels = spd.labels, ordered = TRUE),
      dir.binned = factor(dir.binned, levels = dir.labels, ordered = TRUE)
    ) %>%
    count(across(all_of(c(group_vars, "dir.binned", "spd.binned"))), name = "n") %>%
    group_by(across(all_of(group_vars))) %>%
    mutate(p = n / sum(n)) %>%
    ungroup()
  
  # palette
  n.colors <- length(spd.labels)
  spd.colors <- colorRampPalette(brewer.pal(min(9,n.colors), palette))(n.colors)
  spd.colors <- rev(spd.colors)
  
  # reverse factor levels so high speeds stack outward
  wind_binned <- wind_binned %>%
    mutate(spd.binned = factor(spd.binned, levels = rev(levels(spd.binned)), ordered = TRUE))
  
  ggplot(wind_binned, aes(x = dir.binned, y = p, fill = spd.binned, color = spd.binned)) +
    geom_col(position = "stack") +
    scale_y_continuous(labels = percent) +
    scale_x_discrete(drop = FALSE, labels = dir.labels) +
    coord_polar(start = -((dirres/2)/360) * 2 * pi) +
    scale_fill_manual(name = "Wind Speed (m/s)",
                      values = spd.colors,
                      drop = FALSE,
                      guide = guide_legend(reverse = TRUE)) +
    scale_color_manual(name = "Wind Speed (m/s)",
                      values = spd.colors,
                      drop = FALSE,
                      guide = guide_legend(reverse = TRUE))+
    labs(title = "Windspeed and direction", x = NULL, y = "Frequency")
}
```

```
# different color scaling

make_windrose <- function(wind,
                          spd_col = "Windspeed",
                          dir_col = "Winddir",
                          group_vars = "Station",
                          spdres = 2,
                          dirres = 22.5,
                          palette = "YlGnBu") {
  
  # dynamic binning based on max observed speed
  max_speed <- floor(max(wind[[spd_col]], na.rm = TRUE))
cutoff <- (max_speed - 7)
if (cutoff < 1) cutoff <- 1

spd.breaks <- c(0, cutoff, (cutoff+1):max_speed, Inf)
spd.labels <- c(paste0("0-", cutoff),
                as.character((cutoff+1):max_speed),
                paste0(max_speed, "+"))

n.colors <- length(spd.labels)
spd.colors <- colorRampPalette(brewer.pal(min(9, n.colors), palette))(n.colors)

spd.colors <- rev(spd.colors)

  
  # direction bins
  dir.breaks <- c(seq(0, 360, by = dirres))
  dir.labels <- c("N","NNE","NE","ENE","E","ESE","SE","SSE",
                  "S","SSW","SW","WSW","W","WNW","NW","NNW")
  
  wind_binned <- wind %>%
    filter(!is.na(.data[[spd_col]]), !is.na(.data[[dir_col]])) %>%
    mutate(
      spd.binned = cut(.data[[spd_col]],
                       breaks = spd.breaks,
                       labels = spd.labels,
                       right = FALSE,
                       include.lowest = TRUE),
      dir.binned = cut(.data[[dir_col]],
                       breaks = dir.breaks,
                       labels = dir.labels,
                       right = FALSE,
                       include.lowest = TRUE),
      spd.binned = factor(spd.binned, levels = spd.labels, ordered = TRUE),
      dir.binned = factor(dir.binned, levels = dir.labels, ordered = TRUE)
    ) %>%
    count(across(all_of(c(group_vars, "dir.binned", "spd.binned"))), name = "n") %>%
    group_by(across(all_of(group_vars))) %>%
    mutate(p = n / sum(n)) %>%
    ungroup()
  
  # reverse factor levels so high speeds stack outward
  wind_binned <- wind_binned %>%
    mutate(spd.binned = factor(spd.binned, levels = rev(levels(spd.binned)), ordered = TRUE))
  
  ggplot(wind_binned, aes(x = dir.binned, y = p, fill = spd.binned,color=spd.binned)) +
    geom_col(position = "stack") +
    scale_y_continuous(labels = percent) +
    scale_x_discrete(drop = FALSE, labels = dir.labels) +
    coord_polar(start = -((dirres/2)/360) * 2 * pi) +
    scale_fill_manual(name = "Wind Speed (m/s)",
                      values = spd.colors,
                      drop = FALSE,
                      guide = guide_legend(reverse = TRUE)) +
    scale_color_manual(name = "Wind Speed (m/s)",
                      values = spd.colors,
                      drop = FALSE,
                      guide = guide_legend(reverse = TRUE))+
    labs(title = "Windspeed and direction", x = NULL, y = "Frequency")
}
```

```
#pdf("windrose2m.pdf",width=8,height=9)
p <- make_windrose(wind, group_vars = "Station",spdres = 2) +
  facet_wrap(~ Station,ncol=2,scales="free")+
  theme_minimal()+
  labs(title="Windroses",subtitle = "measured 2m above ground")
p
```

```
#dev.off()
```

```
wind <- wind %>%
  mutate(hour = hour(DateTimeCL),
         daynight = ifelse(hour >= 7 & hour < 19, "Day", "Night"),
         daynight = factor(daynight, levels = c("Day", "Night")))


#pdf("nightwinds.pdf",height=9,width=12)
# Create the day plot
p_day <- make_windrose(filter(wind, daynight == "Day", .preserve = TRUE)) +
  facet_wrap(~ Station,ncol=2) +
  theme_minimal() +
  labs(title = "Daytime Winds (7:00-19:00)")+
  theme(legend.position = "none")

# Create the night plot with dark theme
p_night <- make_windrose(filter(wind, daynight == "Night", .preserve = TRUE)) +
  facet_wrap(~ Station,ncol=2) +
  theme_minimal(base_family = "sans") +
  theme(
    panel.background = element_rect(fill = "black"),
    plot.background  = element_rect(fill = "black"),
    axis.text        = element_text(color = "white"),
    axis.title       = element_text(color = "white"),
    legend.background= element_rect(fill = "black"),
    legend.text      = element_text(color = "white"),
    legend.title     = element_text(color = "white"),
    strip.background = element_rect(fill = "black"),
    strip.text       = element_text(color = "white"),
    plot.title       = element_text(color = "white")
  ) +
  labs(title = "Nighttime Winds (19:00-7:00)",y="")

# Arrange them side by side
p_day + p_night
```

```
#dev.off()
```

#### Yearly summaries

```
###### AGGREGATED BY STATION


# 1. Add day/night classification
weatherdata_long_DN <- weatherdata_long %>%
  mutate(period = ifelse(hour(DateTimeCL) >= 7 & hour(DateTimeCL) < 19, "Day", "Night"))

# 2a. Monthly summaries with parameter-specific aggregation (Day/Night)
monthly_means <- weatherdata_long_DN %>%
  mutate(year = year(DateTimeCL),
         month = month(DateTimeCL)) %>%
  group_by(Region, Station, year, month, parameter, period) %>%
  summarise(
    mean_val = mean(value, na.rm = TRUE),
    max_val = max(value, na.rm = TRUE),
    min_val = min(value, na.rm = TRUE),
    sum_val = case_when(
      parameter %in% c("Nebel","Nebelmenge","Regen") ~ sum(value, na.rm = TRUE),
      TRUE ~ NA_real_
    ),
    n_days = n_distinct(as.Date(DateTimeCL)),
    .groups = "drop"
  ) %>%
  filter(n_days >= 25)%>%
  distinct(Region, Station, year, month, parameter, period, .keep_all = TRUE)

# 2b. Add Total for continuous parameters (combine Day+Night)
monthly_totals <- weatherdata_long_DN %>%
  mutate(year = year(DateTimeCL),
         month = month(DateTimeCL)) %>%
  group_by(Region, Station, year, month, parameter) %>%
  summarise(
    mean_val = mean(value, na.rm = TRUE),
    max_val  = max(value, na.rm = TRUE),
    min_val  = min(value, na.rm = TRUE),
    sum_val  = case_when(
      parameter %in% c("Nebel","Nebelmenge","Regen") ~ sum(value, na.rm = TRUE),
      TRUE ~ NA_real_),
    n_days   = n_distinct(as.Date(DateTimeCL)),
    .groups = "drop"
  ) %>%
  filter(n_days >= 25) %>%
  mutate(period = "Total")%>%
  distinct(Region, Station, year, month, parameter, period, .keep_all = TRUE)

# Combine both
monthly_means <- bind_rows(monthly_means, monthly_totals)

monthly_means <- monthly_means %>%
  distinct(Region, Station, year, month, parameter, period, .keep_all = TRUE)


# 3. Month-balanced yearly averages
monthly_avg_over_years <- monthly_means %>%
  group_by(Region, Station, month, parameter, period) %>%
  summarise(
    mean_month = mean(mean_val, na.rm = TRUE),
    max_month  = mean(max_val, na.rm = TRUE),
    min_month  = mean(min_val, na.rm = TRUE),
    sum_month  = mean(sum_val, na.rm = TRUE),   # average monthly sums across years
    n_years    = n(),
    .groups = "drop"
  )

yearly_balanced <- monthly_avg_over_years %>%
  group_by(Region, Station, parameter, period) %>%
  summarise(
    mean_year = mean(mean_month, na.rm = TRUE),
    max_year  = mean(max_month, na.rm = TRUE),
    min_year  = mean(min_month, na.rm = TRUE),
    sum_year  = sum(sum_month, na.rm = TRUE),
    n_months_used      = n(),
    .groups = "drop"
  )

# 4. Seasonal averages (DJF, MAM, JJA, SON)
seasonal_means <- monthly_means %>%
  mutate(season = case_when(
    month %in% c(12,1,2)   ~ "SummerDJF",
    month %in% c(3,4,5)    ~ "AutumnMAM",
    month %in% c(6,7,8)    ~ "WinterJJA",
    month %in% c(9,10,11)  ~ "SpringSON"
  )) %>%
  group_by(Region, Station, year, season, parameter, period) %>%
  summarise(
    # continuous stats: averaged across the 3 months
    mean_season = mean(mean_val, na.rm = TRUE),
    max_season  = mean(max_val,  na.rm = TRUE),
    min_season  = mean(min_val,  na.rm = TRUE),
    sum_season  = sum(sum_val,   na.rm = TRUE),
    n_months    = n_distinct(month),
    .groups = "drop"
  ) %>%
  filter(n_months >= 2)  # require complete seasons

seasonal_avg_over_years <- seasonal_means %>%
  group_by(Region, Station, season, parameter, period) %>%
  summarise(
    mean_season = mean(mean_season, na.rm = TRUE),
    max_season  = mean(max_season,  na.rm = TRUE),
    min_season  = mean(min_season,  na.rm = TRUE),
    sum_season  = mean(sum_season,  na.rm = TRUE),  # average seasonal totals across years
    n_seasons_used      = n(),
    .groups = "drop"
  )

# 5. Pivot to wide format (yearly)
yearly_wide <- yearly_balanced %>%
  pivot_wider(
    names_from = c(parameter, period),
    values_from = c(mean_year, max_year, min_year, sum_year),
    names_glue = "{parameter}_{period}_{.value}"
  ) %>%
  mutate(method = "yearly")

# Seasonal wide via long -> wide (robust to NA combos)
seasonal_wide <- seasonal_avg_over_years %>%
  pivot_longer(
    cols = c(mean_season, max_season, min_season, sum_season),
    names_to = "stat", values_to = "value"
  ) %>%
  pivot_wider(
    names_from = c(parameter, period, stat),
    values_from = value,
    names_glue = "{parameter}_{period}_{stat}"
  ) %>%
  mutate(method = "seasonal")
```

```
######## AGGREGATED BY REGION

# 1. Add day/night classification
weatherdata_long_DN <- weatherdata_long %>%
  mutate(period = ifelse(hour(DateTimeCL) >= 7 & hour(DateTimeCL) < 19, "Day", "Night"))

# 2a. Monthly summaries with parameter-specific aggregation (Day/Night)
monthly_means_region <- weatherdata_long_DN %>%
  mutate(year = year(DateTimeCL),
         month = month(DateTimeCL)) %>%
  group_by(Region,year, month, parameter, period) %>%
  summarise(
    mean_val = mean(value, na.rm = TRUE),
    max_val = max(value, na.rm = TRUE),
    min_val = min(value, na.rm = TRUE),
    sum_val = case_when(
      parameter %in% c("Nebel","Nebelmenge","Regen") ~ sum(value, na.rm = TRUE),
      TRUE ~ NA_real_
    ),
    n_days = n_distinct(as.Date(DateTimeCL)),
    .groups = "drop"
  ) %>%
  filter(n_days >= 25)%>%
  distinct(Region, year, month, parameter, period, .keep_all = TRUE)

# 2b. Add Total for continuous parameters (combine Day+Night)
monthly_totals_region <- weatherdata_long_DN %>%
  mutate(year = year(DateTimeCL),
         month = month(DateTimeCL)) %>%
  group_by(Region, year, month, parameter) %>%
  summarise(
    mean_val = mean(value, na.rm = TRUE),
    max_val  = max(value, na.rm = TRUE),
    min_val  = min(value, na.rm = TRUE),
    sum_val  = case_when(
      parameter %in% c("Nebel","Nebelmenge","Regen") ~ sum(value, na.rm = TRUE),
      TRUE ~ NA_real_),
    n_days   = n_distinct(as.Date(DateTimeCL)),
    .groups = "drop"
  ) %>%
  filter(n_days >= 25) %>%
  mutate(period = "Total")%>%
  distinct(Region, year, month, parameter, period, .keep_all = TRUE)

# Combine both
monthly_means_region <- bind_rows(monthly_means_region, monthly_totals_region)

monthly_means_region <- monthly_means_region %>%
  distinct(Region, year, month, parameter, period, .keep_all = TRUE)


# 3. Month-balanced yearly averages
monthly_avg_over_years_region <- monthly_means_region %>%
  group_by(Region, month, parameter, period) %>%
  summarise(
    mean_month = mean(mean_val, na.rm = TRUE),
    max_month  = mean(max_val, na.rm = TRUE),
    min_month  = mean(min_val, na.rm = TRUE),
    sum_month  = mean(sum_val, na.rm = TRUE),   # average monthly sums across years
    n_years    = n(),
    .groups = "drop"
  )

yearly_balanced_region <- monthly_avg_over_years_region %>%
  group_by(Region, parameter, period) %>%
  summarise(
    mean_year = mean(mean_month, na.rm = TRUE),
    max_year  = mean(max_month, na.rm = TRUE),
    min_year  = mean(min_month, na.rm = TRUE),
    sum_year  = sum(sum_month, na.rm = TRUE),
    n_months_used      = n(),
    .groups = "drop"
  )

# 4. Seasonal averages (DJF, MAM, JJA, SON)
seasonal_means_region <- monthly_means_region %>%
  mutate(season = case_when(
    month %in% c(12,1,2)   ~ "SummerDJF",
    month %in% c(3,4,5)    ~ "AutumnMAM",
    month %in% c(6,7,8)    ~ "WinterJJA",
    month %in% c(9,10,11)  ~ "SpringSON"
  )) %>%
  group_by(Region, year, season, parameter, period) %>%
  summarise(
    # continuous stats: averaged across the 3 months
    mean_season = mean(mean_val, na.rm = TRUE),
    max_season  = mean(max_val,  na.rm = TRUE),
    min_season  = mean(min_val,  na.rm = TRUE),
    sum_season  = sum(sum_val,   na.rm = TRUE),
    n_months    = n_distinct(month),
    .groups = "drop"
  ) %>%
  filter(n_months >= 2)  # require complete seasons

seasonal_avg_over_years_region <- seasonal_means_region %>%
  group_by(Region, season, parameter, period) %>%
  summarise(
    mean_season = mean(mean_season, na.rm = TRUE),
    max_season  = mean(max_season,  na.rm = TRUE),
    min_season  = mean(min_season,  na.rm = TRUE),
    sum_season  = mean(sum_season,  na.rm = TRUE),  # average seasonal totals across years
    n_seasons_used      = n(),
    .groups = "drop"
  )

# 5. Pivot to wide format (yearly)
yearly_wide_region <- yearly_balanced_region %>%
  pivot_wider(
    names_from = c(parameter, period),
    values_from = c(mean_year, max_year, min_year, sum_year),
    names_glue = "{parameter}_{period}_{.value}"
  ) %>%
  mutate(method = "yearly")

# Seasonal wide via long -> wide (robust to NA combos)
seasonal_wide_region <- seasonal_avg_over_years_region %>%
  pivot_longer(
    cols = c(mean_season, max_season, min_season, sum_season),
    names_to = "stat", values_to = "value"
  ) %>%
  pivot_wider(
    names_from = c(parameter, period, stat),
    values_from = value,
    names_glue = "{parameter}_{period}_{stat}"
  ) %>%
  mutate(method = "seasonal")
```

```
yearly_wide
```

```
# Define units
param_units <- c(
  Temp = "°C",
  Windspeed = "m/s",
  Winddir = "°deg"
)

# Build nested compact summary from yearly_balanced
# by Station
compact_summary <- yearly_balanced %>%
  mutate(
    unit = param_units[parameter],
    # Create a nested string for each row depending on parameter type
    stats = case_when(
      parameter %in% c("Nebel","Nebelmenge","Regen") ~ paste0("sum=", round(sum_year, 1)),
      TRUE ~ paste0(round(mean_year, 1),
                    " (", round(min_year, 1),
                    "-", round(max_year, 1),")")
    )
  ) %>%
  select(Region, Station, n_months_used, parameter, unit, period, stats) %>%
  pivot_wider(names_from = period, values_from = stats)

# by Region
compact_summary_region <- yearly_balanced_region %>%
  mutate(
    unit = param_units[parameter],
    # Create a nested string for each row depending on parameter type
    stats = case_when(
      parameter %in% c("Nebel","Nebelmenge","Regen") ~ paste0("sum=", round(sum_year, 1)),
      TRUE ~ paste0(round(mean_year, 1),
                    " (", round(min_year, 1),
                    "-", round(max_year, 1),")")
    )
  ) %>%
  select(Region, n_months_used, parameter, unit, period, stats) %>%
  pivot_wider(names_from = period, values_from = stats)


# Seasonal compact summary (nested Day/Night/Total)
# by Station
seasonal_compact_summary <- seasonal_avg_over_years %>%
  mutate(
    unit = param_units[parameter],
    stats = case_when(
      parameter %in% c("Nebel","Nebelmenge","Regen") ~ paste0("sum=", round(sum_season, 1)),
      TRUE ~ paste0(
        round(mean_season, 1), " (",
        round(min_season, 1), "-",
        round(max_season, 1), ")"
      )
    )
  ) %>%
  select(Region, Station, n_seasons_used, season, parameter, unit, period, stats) %>%
  pivot_wider(names_from = period, values_from = stats)

# by Region
seasonal_compact_summary_region <- seasonal_avg_over_years_region %>%
  mutate(
    unit = param_units[parameter],
    stats = case_when(
      parameter %in% c("Nebel","Nebelmenge","Regen") ~ paste0("sum=", round(sum_season, 1)),
      TRUE ~ paste0(
        round(mean_season, 1), " (",
        round(min_season, 1), "-",
        round(max_season, 1), ")"
      )
    )
  ) %>%
  select(Region, n_seasons_used, season, parameter, unit, period, stats) %>%
  pivot_wider(names_from = period, values_from = stats)
```

##### by Station

```
# selected parameters
compact_summary %>% filter(parameter %in% c("Temp","Windspeed"))
```

##### by Region

```
compact_summary_region %>% filter(parameter %in% c("Temp","Windspeed"))
```

#### Plots of mean of daily mean/min/max

```
# Define units
param_units <- c(
  Nebelmenge = "ml", Regen = "mm", Temp = "°C", TempTill = "°C",
  Nässe = "%", Windspeed = "m/s"
)

selected_params <- names(param_units)

# Aggregate to one row per Region × Station × Season × Parameter
plot_data <- seasonal_avg_over_years %>%
  filter(parameter %in% selected_params) %>%
  group_by(Region, Station, season, parameter) %>%
  summarise(
    mean = mean(mean_season, na.rm = TRUE),
    min = mean(min_season, na.rm = TRUE),
    max = mean(max_season, na.rm = TRUE),
    sum = mean(sum_season, na.rm = TRUE),
    .groups = "drop"
  ) %>%
  mutate(
    season = factor(season, levels = c("SummerDJF","AutumnMAM","WinterJJA","SpringSON")),
    unit = param_units[parameter],
    facet_label = paste0(parameter, " [", unit, "]")
  )

plot_data$facet_label[plot_data$facet_label == "Windspeed [m/s]"] <- "Wind [m/s]"

ggplot() +
  # Continuous: mean dot + min/max ticks
  geom_point(data = filter(plot_data, !parameter %in% c("Nebelmenge","Regen")),
             aes(x = season, y = mean, color = Station), size = 2,position = position_dodge(width=0.2)) +
  geom_errorbar(data = filter(plot_data, !parameter %in% c("Nebelmenge","Regen")),
                aes(x = season, ymin = min, ymax = max, color = Station),
                width = 0.1, linewidth = 0.4, position = position_dodge(width=0.2)) +
# Mean labels (centered)
geom_text(data = filter(plot_data, !parameter %in% c("Nebelmenge","Regen")),
          aes(x = season, y = mean, label = round(mean, 1), color = Station),
          vjust = 0, size = 3, position = position_dodge(width = 0.9)) +

# Min labels (below)
geom_text(data = filter(plot_data, !parameter %in% c("Nebelmenge","Regen")),
          aes(x = season, y = min, label = round(min, 1), color = Station),
          vjust = 0, size = 2.8, position = position_dodge(width = 0.9)) +

# Max labels (above)
geom_text(data = filter(plot_data, !parameter %in% c("Nebelmenge","Regen")),
          aes(x = season, y = max, label = round(max, 1), color = Station),
          vjust = 0, size = 2.8, position = position_dodge(width = 0.9))+
  
  # Event bar labels (sum values)
geom_text(data = filter(plot_data, parameter %in% c("Nebelmenge","Regen")),
          aes(x = season, y = sum, label = round(sum, 1), group = Station),
          position = position_dodge(width = 0.9),
          vjust = -0.5, size = 3)+


  # Events: seasonal sums as bars
  geom_col(data = filter(plot_data, parameter %in% c("Nebelmenge","Regen")),
           aes(x = season, y = sum, fill = Station), position = "dodge") +

  facet_grid(facet_label ~ Region, scales = "free_y") +
  labs(title = "Seasonal min–mean–max (continuous) and sums (events)",
       y = "Value", x = "Season") +
  theme_bw() +
  theme(axis.text.x = element_text(angle = 45, hjust = 1))+
  scale_y_continuous(expand = expansion(mult = c(0.05, 0.2)))+
  labs(title = "Seasonal mean daily min–mean–max values and sums (fog/rain)",
       subtitle = "● mean, | min/max range, ▮ seasonal sum",
       y = "Value")
```

```
# Define parameters and units
param_units <- c(
  Nebelmenge = "ml", Regen = "mm", Temp = "°C", TempTill = "°C",
  Nässe = "%", Windspeed = "m/s"
)

selected_params <- names(param_units)

# Prepare yearly data
plot_yearly <- yearly_balanced %>%
  filter(parameter %in% selected_params) %>%
  group_by(Region, Station, parameter) %>%
  summarise(
    mean = mean(mean_year, na.rm = TRUE),
    min = mean(min_year, na.rm = TRUE),
    max = mean(max_year, na.rm = TRUE),
    sum = mean(sum_year, na.rm = TRUE),
    .groups = "drop"
  ) %>%
  mutate(
    unit = param_units[parameter],
    facet_label = paste0(parameter, " [", unit, "]")
  ) 

plot_yearly$facet_label[plot_yearly$facet_label == "Windspeed [m/s]"] <- "Wind [m/s]"

ggplot() +
  # Continuous: mean dot + min/max ticks
  geom_point(data = filter(plot_yearly, !parameter %in% c("Nebelmenge","Regen")),
             aes(x = Region, y = mean, color = Station), size = 2, position = position_dodge(width=0.1)) +
  geom_errorbar(data = filter(plot_yearly, !parameter %in% c("Nebelmenge","Regen")),
                aes(x = Region, ymin = min, ymax = max, color = Station),
                width = 0.1, linewidth = 0.4, position = position_dodge(width=0.1)) +

  # Mean labels
  geom_text(data = filter(plot_yearly, !parameter %in% c("Nebelmenge","Regen")),
            aes(x = Region, y = mean, label = round(mean, 1), color = Station),
            hjust = 0, size = 3, position = position_dodge(width = 0.7)) +

  # Min labels
  geom_text(data = filter(plot_yearly, !parameter %in% c("Nebelmenge","Regen")),
            aes(x = Region, y = min, label = round(min, 1), color = Station),
            hjust = 0, size = 2.8, position = position_dodge(width = 0.7)) +

  # Max labels
  geom_text(data = filter(plot_yearly, !parameter %in% c("Nebelmenge","Regen")),
            aes(x = Region, y = max, label = round(max, 1), color = Station),
            hjust = 0, size = 2.8, position = position_dodge(width = 0.7)) +

  # Events: yearly sums as bars
  geom_col(data = filter(plot_yearly, parameter %in% c("Nebelmenge","Regen")),
           aes(x = Region, y = sum, fill = Station), position = "dodge") +

  # Event bar labels
  geom_text(data = filter(plot_yearly, parameter %in% c("Nebelmenge","Regen")),
            aes(x = Region, y = sum, label = round(sum, 1), group = Station),
            position = position_dodge(width = 0.9),
            vjust = -0.5, size = 3) +

  facet_grid(facet_label ~ Region, scales = "free") +
  labs(
    title = "Yearly mean daily min–mean–max values and sums (fog/rain)",
    subtitle = "● mean, | min/max range, ▮ yearly sum",
    y = "Value", x = ""
  ) +
  theme_bw() +
  theme(axis.text.x = element_text(angle = 45, hjust = 1)) +
  scale_y_continuous(expand = expansion(mult = c(0.05, 0.2)))+
  scale_x_discrete(drop = TRUE)
```

#### Drift Potential

##### Drift Potential Sandrose

```
## Extrapolate wind speeds at 10m height from wind speeds measured at a lower point

windsensorheight <- 2.5 # height of wind measurements in m

## formula

#z0 <- 0.002 # smooth sand
z0 <- 0.05 # rough ground / vegetation / Heidekraut

wind_profile_10m <- function(v, h = height, d = 0.0, z0 = z0) {
  v * log((10 - d) / z0) / log((h - d) / z0)
}
```

```
## convert wind speed to 10m

weatherdata$Windspeed_10m <- wind_profile_10m(weatherdata$Windspeed, h = windsensorheight, z0 = z0)

weatherdata <- weatherdata %>% mutate(Station = factor(Station,
                                                       levels = c("WSA1", "WSA2","OYA1128","OYA1211","MSC3","MSC1")))
```

```
#pdf("windrose10m.pdf",height=9,width=8)
make_windrose(weatherdata,spd_col = "Windspeed_10m",group_vars = "Station")+
  facet_wrap(~Station,ncol=2)+
  theme_minimal()+
  labs(title="Windroses",subtitle="extrapolated 10m above ground")
```

```
#dev.off()
```

```
## Convert to knots; 1 m/s = 1.94384 knots

weatherdata$wind_knots <- weatherdata$Windspeed_10m * 1.94384 # use extrapolated wind data
#weatherdata$wind_knots <- weatherdata$Windspeed * 1.94384 # use raw wind data
```

```
# Define 16 compass sectors (22.5° wide)
breaks <- seq(-11.25, 371.25, by = 22.5)   # sector boundaries
labels <- c("N","NNE","NE","ENE","E","ESE","SE","SSE",
            "S","SSW","SW","WSW","W","WNW","NW","NNW","N")

# Cut wind direction into compass bins
weatherdata$Winddir_label <- cut(weatherdata$Winddir,
                             breaks = breaks,
                             labels = labels,
                             include.lowest = TRUE)
```

```
## assign wind classes as Fryberger & Dean (1979)

bins <- c(0, 11, 16.5, 21.5, 27.5, 33.5, 40, Inf)
labels <- c("<11", "11-16", "17-21", "22-27", "28-33", "34-40", "≥40")

weatherdata$wind_class <- cut(weatherdata$wind_knots, breaks = bins, labels = labels, right = FALSE)
```

Weighting factors (Fryberger & Dean):

- <11 = 0
- 11–16 = 2.7
- 17–21 = 25.3
- 22–27 = 75.0
- 28–33 = 172.1
- 34–40 = 342.3
- ≥40 = 0

```
weights <- c("<11"=0, "11-16"=2.7, "17-21"=25.3, "22-27"=75.0, "28-33"=172.1, "34-40"=342.3, "≥40"=0)

weatherdata$weightingfactor <- weights[as.character(weatherdata$wind_class)]

drift_data <- weatherdata %>%
  group_by(Station, Winddir_label, wind_class) %>%
  summarise(counts = n(), .groups="drop") %>%
  mutate(rel_freq = counts / sum(counts) * 100,
         drift_potential = rel_freq * weights[as.character(wind_class)])
```

```
# Filter NAs

drift_data <- drift_data %>% filter(!Winddir_label == "NA")

# assign order
drift_data <- drift_data %>%
  mutate(Station = factor(Station,
                          levels = c("WSA1", "WSA2", "OYA1128","OYA1211","MSC3","MSC1")))
```

```
library(dplyr)

# Assume df already has:
# Winddir (°), wind_knots, wind_class (factor from cut()), etc.

# Define weighting factors (Fryberger & Dean 1979)
weights <- c("<11"=0, "11-16"=2.7, "17-21"=25.3,
             "22-27"=75.0, "28-33"=172.1,
             "34-40"=342.3, "≥40"=0)

# Group by wind direction and class
grouped_table <- weatherdata %>%
  group_by(Station, Winddir_label, wind_class) %>%
  summarise(counts = n(), .groups="drop") %>%
  mutate(
    total_counts = sum(counts),                        # total per direction
    rel_freq = counts / total_counts * 100,            # relative frequency (%)
    weightingfactor = weights[as.character(wind_class)],
    drift_potential = rel_freq * weightingfactor       # Q = f × t
  )

print(grouped_table)
```

```
## # A tibble: 232 × 8
##    Station Winddir_label wind_class counts total_counts rel_freq weightingfactor
##    <fct>   <fct>         <fct>       <int>        <int>    <dbl>           <dbl>
##  1 WSA1    N             <11          2580       425989 0.606                0  
##  2 WSA1    N             11-16          39       425989 0.00916              2.7
##  3 WSA1    N             17-21           4       425989 0.000939            25.3
##  4 WSA1    NNE           <11          2587       425989 0.607                0  
##  5 WSA1    NNE           11-16          23       425989 0.00540              2.7
##  6 WSA1    NNE           17-21           3       425989 0.000704            25.3
##  7 WSA1    NE            <11          2389       425989 0.561                0  
##  8 WSA1    NE            11-16          50       425989 0.0117               2.7
##  9 WSA1    NE            17-21           4       425989 0.000939            25.3
## 10 WSA1    ENE           <11          2614       425989 0.614                0  
## # ℹ 222 more rows
## # ℹ 1 more variable: drift_potential <dbl>
```

##### Net Sand Drift vector

```
plot_sand_rose <- function(drift_data) {
  library(dplyr)
  library(ggplot2)
  library(grid)

  # Define compass angles (English 16-point)
  compass_angles <- c("N"=0, "NNE"=22.5, "NE"=45, "ENE"=67.5,
                      "E"=90, "ESE"=112.5, "SE"=135, "SSE"=157.5,
                      "S"=180, "SSW"=202.5, "SW"=225, "WSW"=247.5,
                      "W"=270, "WNW"=292.5, "NW"=315, "NNW"=337.5)

  # Ensure Winddir_label is a factor with correct order
  drift_data$Winddir_label <- factor(drift_data$Winddir_label,
                                          levels = names(compass_angles))

  # Add angle and vector components
  drift_data <- drift_data %>%
    mutate(angle_rad = compass_angles[as.character(Winddir_label)] * pi / 180,
           dp_x = drift_potential * sin(angle_rad),
           dp_y = drift_potential * cos(angle_rad))

  # Compute vector summary per station
  vector_summary <- drift_data %>%
    group_by(Station) %>%
    summarise(
      DP = sum(drift_potential, na.rm = TRUE),
      RDP_x = sum(dp_x, na.rm = TRUE),
      RDP_y = sum(dp_y, na.rm = TRUE),
      RDP = sqrt(RDP_x^2 + RDP_y^2),
      RDP_angle = atan2(RDP_x, RDP_y) * 180 / pi,
      directionality = ifelse(DP > 0, RDP / DP, NA_real_),
      .groups = "drop"
    )
  
  # Flip the vector components
vector_summary <- vector_summary %>%
  mutate(
    RDP_x = -RDP_x,
    RDP_y = -RDP_y,
    RDP_angle = atan2(RDP_x, RDP_y) * 180 / pi
  )


  # Map RDP angle to nearest compass sector for plotting
  nearest_sector <- function(angle_deg) {
    angle_deg <- angle_deg %% 360
    diffs <- abs(unlist(compass_angles) - angle_deg)
    names(compass_angles)[which.min(diffs)]
  }

  vector_summary$RDP_sector <- sapply(vector_summary$RDP_angle, nearest_sector)

  # Compute max drift per station for scaling
  max_dp <- drift_data %>%
    group_by(Station) %>%
    summarise(max_dp = max(drift_potential, na.rm = TRUE), .groups = "drop")

  vector_summary <- vector_summary %>%
    left_join(max_dp, by = "Station") %>%
    mutate(scaled_RDP = RDP / max_dp * max_dp)  # scale to match each station's plot

  # Plot
  ggplot(drift_data, aes(x = Winddir_label, y = drift_potential, fill = wind_class)) +
    geom_bar(stat = "identity") +
    coord_polar(start = -pi/16) +
    facet_wrap(~Station, ncol = 2, scales = "free") +
    geom_segment(
      data = vector_summary,
      aes(x = RDP_sector, xend = RDP_sector, y = 0, yend = scaled_RDP),
      inherit.aes = FALSE, color = "red", size = 1.1,
      arrow = arrow(length = unit(0.25, "cm"))
    ) +
    geom_text(
      data = vector_summary,
      aes(x = "N", y = max_dp * 0.9,
          label = paste0("DP = ", round(DP, 1),
                         "\nRDP = ", round(RDP, 1),
                         "\nRDP/DP = ", round(directionality, 2))),
      inherit.aes = FALSE, size = 3, hjust = 0
    ) +
    labs(title = "Sand Drift Potential Rose",
         x = "Wind Direction", y = "Drift Potential") +
    theme_minimal(base_size = 12) +
    theme(strip.text = element_text(face = "bold"),
          axis.text.x = element_text(size = 8))+
    scale_fill_discrete(name="Wind class (kn)")
}
```

```
#pdf("driftrose_10m_z0_5cm.pdf",width=10,height=10)
plot_sand_rose(drift_data)
```

```
#dev.off()
```

#### Mean exposition vector

```
Sanddata <- read_excel("Sanddata.xlsx")
Sanddata$aspect <- as.numeric(Sanddata$aspect)


# Define 16 compass sectors (22.5° wide)
breaks <- seq(-11.25, 371.25, by = 22.5)   # sector boundaries
labels <- c("N","NNE","NE","ENE","E","ESE","SE","SSE",
            "S","SSW","SW","WSW","W","WNW","NW","NNW","N")

# Cut exposition direction into compass bins
Sanddata$aspect_sector <- cut(Sanddata$aspect,
                             breaks = breaks,
                             labels = labels,
                             include.lowest = TRUE)

aspectdata <- Sanddata %>% filter(!aspect_sector == "NA")
```

```
# adjust order 
aspectdata <- aspectdata %>%
  mutate(fieldID = factor(fieldID,
                          levels = c("Arica", "Oyarbide", "Caldera")))


plot_aspect_rose <- function(df,
                             aspect_col = "aspect",
                             sector_col = "aspect_sector",
                             group_col = "fieldID",
                             state_col = "plantstate") {
  
  df[[state_col]] <- factor(
  df[[state_col]],
  levels = c("PLANT", "DEAD", "NO_PLANT")
)

  state_colors <- c(
  "PLANT" = "forestgreen",
  "DEAD" = "firebrick",
  "NO_PLANT" = "grey60"
)

  state_labels <- c(
  "PLANT" = "Plant",
  "DEAD" = "Dead",
  "NO_PLANT" = "No plant"
)

  
  # 16 compass labels
  compass_labels <- c("N","NNE","NE","ENE","E","ESE","SE","SSE",
                      "S","SSW","SW","WSW","W","WNW","NW","NNW")
  dirres <- 22.5
  
  
  # FAN SHAPE DATA (absolute counts)

  binned_fans <- df %>%
    filter(!is.na(.data[[sector_col]])) %>%
    group_by(.data[[group_col]], .data[[sector_col]]) %>%
    summarise(counts = n(), .groups = "drop")
  

  # FAN COLOR DATA (absolute counts per plantstate)

  binned_colors <- df %>%
    filter(!is.na(.data[[sector_col]])) %>%
    group_by(.data[[group_col]], .data[[sector_col]], .data[[state_col]]) %>%
    summarise(counts = n(), .groups = "drop")
  
  
  # resultant vector helpers
  
  compass_angles <- setNames(seq(0, 337.5, by = 22.5), compass_labels)
  
  nearest_sector <- function(angle_deg) {
    angle_deg <- angle_deg %% 360
    diffs <- abs(unlist(compass_angles) - angle_deg)
    names(compass_angles)[which.min(diffs)]
  }
  
  
  # OVERALL resultant vector (red)

  res_total <- df %>%
    mutate(theta = (.data[[aspect_col]] %% 360) * pi / 180) %>%
    group_by(.data[[group_col]]) %>%
    summarise(
      x = mean(sin(theta), na.rm = TRUE),
      y = mean(cos(theta), na.rm = TRUE),
      R = sqrt(x^2 + y^2),
      mean_deg = (atan2(x, y) * 180 / pi) %% 360,
      .groups = "drop"
    ) %>%
    mutate(R_sector = sapply(mean_deg, nearest_sector)) %>%
    left_join(
      binned_fans %>%
        group_by(.data[[group_col]]) %>%
        summarise(max_count = max(counts), .groups = "drop"),
      by = group_col
    ) %>%
    mutate(arrow_length = R * max_count)
  
  # resultant vectors (colored)
 
  res_state <- df %>%
    mutate(theta = (.data[[aspect_col]] %% 360) * pi / 180) %>%
    group_by(.data[[group_col]], .data[[state_col]]) %>%
    summarise(
      x = mean(sin(theta), na.rm = TRUE),
      y = mean(cos(theta), na.rm = TRUE),
      R = sqrt(x^2 + y^2),
      mean_deg = (atan2(x, y) * 180 / pi) %% 360,
      .groups = "drop"
    ) %>%
    mutate(R_sector = sapply(mean_deg, nearest_sector)) %>%
    left_join(
      binned_fans %>%
        group_by(.data[[group_col]]) %>%
        summarise(max_count = max(counts), .groups = "drop"),
      by = group_col
    ) %>%
    mutate(arrow_length = R * max_count)
  
  
  # PLOT
  
ggplot( binned_colors, aes_string( x = sector_col, y = "counts", fill = state_col ) )+
  geom_bar(stat = "identity", position = "stack",color="grey40", width = 1,alpha=0.5) +
  scale_x_discrete(drop = FALSE, labels = compass_labels) +
  coord_polar(start = -((dirres/2)/360) * 2 * pi) +
  facet_wrap(vars(.data[[group_col]]), ncol = 1, scales = "free") +
  labs(title = "Exposition Rose", x = NULL, y = "Count") +
  theme_minimal(base_size = 12) +
  
  # labels
  scale_fill_manual(values = state_colors, labels = state_labels) +
  scale_color_manual(values = state_colors, labels = state_labels) +
  
  # resultant vector
  geom_segment(
    data = res_total,
    aes(x = R_sector, xend = R_sector, y = 0, yend = arrow_length),
    inherit.aes = FALSE,
    linewidth = 1.3,
    color = "red",
    arrow = arrow(length = unit(0.3, "cm"))
  ) +
  # plant state vectors
#  geom_segment(
#      data = res_state,
#      aes(x = R_sector, xend = R_sector, y = 0, yend = arrow_length, color = plantstate),
#      inherit.aes = FALSE, linewidth = 1.1,
#      arrow = arrow(length = unit(0.25, "cm"))
#    )
  geom_text(
      data = res_total,
      aes(x = R_sector, y = arrow_length + 2,
          label = paste0("Mean = ", round(mean_deg, 1), "°; R = ", round(R, 2))),
      inherit.aes = FALSE, color = "black", size = 3, hjust = 0
    )

}


#pdf("expositionrose.pdf",height=9,width=6)
plot_aspect_rose(aspectdata)
```

```
#dev.off()
```

#### Correlation Matrix and Table

```
fields <- c("Arica", "Oyarbide", "Caldera")

#pdf("corrmatrix.pdf",height=5,width=10)
par(mfrow = c(1, 3))

for (f in fields) {
  # Subset by field
  corrdata <- Sanddata %>%
    filter(fieldID == f) %>%
    select(X, Y, elevation, aspect,
           grainsize_mean, sortingidx, clay, sand, silt)
  
  # Convert to numeric
  corrdata[] <- lapply(corrdata, as.numeric)
  
  # Correlation test
  testRes <- cor.mtest(corrdata, conf.level = 0.95,method="spearman")
  
  # Plot
  corrplot(cor(corrdata, use = "pairwise.complete.obs"),
           col = COL2("PiYG"),
           p.mat = testRes$p,
           sig.level = c(0.001, 0.01, 0.05),
           insig = "label_sig",
           pch.cex = 0.9,
           title = paste(f),
           mar=c(1,1,2,1),
           tl.col="black",
           tl.pos = "lt",
           order="original",
           type="lower",
           method="ellipse")
  corrplot(cor(corrdata, use = "pairwise.complete.obs"),
           col = COL2("PiYG"),
           pch.cex = 0.9,
           mar=c(1,1,2,1),
           tl.pos="n",
           order="original",
           add=T,
           type="upper",
           method="number")
  recordPlot()
}
```

```
#dev.off()

# no field filtering
corrdata <- Sanddata %>%
    select(X, Y, elevation, aspect,
           grainsize_mean, sortingidx, clay, sand, silt)
  
# Convert to numeric
  corrdata[] <- lapply(corrdata, as.numeric)
  
  # Correlation test
  testRes <- cor.mtest(corrdata, conf.level = 0.95,method="spearman")
  
  # Plot
  corrplot(cor(corrdata, use = "pairwise.complete.obs"),
           col = COL2("PiYG"),
           p.mat = testRes$p,
           sig.level = c(0.001, 0.01, 0.05),
           insig = "label_sig",
           pch.cex = 0.9,
           title = paste("Combined Fields"),
           mar=c(1,1,2,1),
           tl.col="black",
           tl.pos = "lt",
           order="original",
           type="lower",
           method="ellipse")
  corrplot(cor(corrdata, use = "pairwise.complete.obs"),
           col = COL2("PiYG"),
           pch.cex = 0.9,
           mar=c(1,1,2,1),
           tl.pos="n",
           order="original",
           add=T,
           type="upper",
           method="number")
```

```
library(dplyr)
library(tidyr)
library(purrr)
library(openxlsx)

fields <- c("Arica", "Oyarbide", "Caldera")

# Choose correction method: "none", "bonferroni", "holm"
correction_method <- "holm"

# Choose p-value display: "threshold" or "stars"
p_display <- "stars"

format_p <- function(p) {
  if (p_display == "threshold") {
    if (p < 0.0001) return("p<0.0001")
    if (p < 0.001) return("p<0.001")
    if (p < 0.01)  return("p<0.01")
    if (p < 0.05)  return("p<0.05")
    return("ns")
  }
  if (p_display == "stars") {
    if (p < 0.0001) return("****")
    if (p < 0.001) return("***")
    if (p < 0.01)  return("**")
    if (p < 0.05)  return("*")
    return("ns")
  }
}

wide_tables <- map(fields, function(f) {
  
  corrdata <- Sanddata %>%
    filter(fieldID == f) %>%
    select(X, Y, elevation, aspect,
           grainsize_mean, sortingidx, clay, sand, silt)
  
  corrdata[] <- lapply(corrdata, as.numeric)
  
  corr_mat <- cor(corrdata, use = "pairwise.complete.obs", method = "spearman")
  testRes  <- cor.mtest(corrdata, conf.level = 0.95, method = "spearman")
  p_mat    <- testRes$p
  
  # Apply multiple-testing correction
  p_vec <- as.vector(p_mat)
  p_adj <- p.adjust(p_vec, method = correction_method)
  p_mat_adj <- matrix(p_adj, nrow = nrow(p_mat), dimnames = dimnames(p_mat))
  
  # Format cells
  formatted <- matrix(
    paste0(
      sprintf("%.3f", corr_mat),
      " (", sapply(p_mat_adj, format_p), ")"
    ),
    nrow = nrow(corr_mat),
    dimnames = dimnames(corr_mat)
  )
  
  df <- as.data.frame(formatted)
  df <- tibble::rownames_to_column(df, var = "variable")
  df$field <- f
  df
})

wide_table_all <- bind_rows(wide_tables)

# Export to Excel
wb <- createWorkbook()

for (df in wide_tables) {
  addWorksheet(wb, unique(df$field))
  writeData(wb, unique(df$field), df)
}

saveWorkbook(wb, "correlation_tables.xlsx", overwrite = TRUE)
```

```
# Create a combined dataset (no field filtering)
corrdata_all <- Sanddata %>%
  select(X, Y, elevation, aspect,
         grainsize_mean, sortingidx, clay, sand, silt)

# Convert to numeric
corrdata_all[] <- lapply(corrdata_all, as.numeric)

# Correlation + p-values
corr_mat_all <- cor(corrdata_all, use = "pairwise.complete.obs", method = "spearman")
testRes_all  <- cor.mtest(corrdata_all, conf.level = 0.95, method = "spearman")
p_mat_all    <- testRes_all$p

# Apply the same correction method you chose earlier
p_vec_all <- as.vector(p_mat_all)
p_adj_all <- p.adjust(p_vec_all, method = correction_method)
p_mat_adj_all <- matrix(p_adj_all, nrow = nrow(p_mat_all), dimnames = dimnames(p_mat_all))

# Format p-values using your chosen display function
formatted_all <- matrix(
  paste0(
    sprintf("%.3f", corr_mat_all),
    " (", sapply(p_mat_adj_all, format_p), ")"
  ),
  nrow = nrow(corr_mat_all),
  dimnames = dimnames(corr_mat_all)
)

wide_all <- as.data.frame(formatted_all)
wide_all <- tibble::rownames_to_column(wide_all, var = "variable")
wide_all$field <- "AllData"

wide_table_all <- bind_rows(wide_tables, list(wide_all))

addWorksheet(wb, "AllData")
writeData(wb, "AllData", wide_all)

saveWorkbook(wb, "correlation_tables.xlsx", overwrite = TRUE)
```
